# Divergent venoms among two closely related co-distributed centipede species, *Scolopendra morsitans* and *S. hardwickei* in tropical Asia

**DOI:** 10.64898/2026.04.01.715817

**Authors:** Aditi, Pragyadeep Roy, Richard Parikh, Aniruddha Marathe, Karunakar Majhi, Ronald Jenner, Jahnavi Joshi

## Abstract

Venom is an important functional trait that helps predatory animals capture prey. Centipede predatory venoms are complex cocktails of multiple proteins, such as neurotoxins (scoloptoxins), cytotoxins, β-pore-forming toxins, and enzymes. We examined venom phenotypes in two closely related and co-occurring centipede species, *Scolopendra morsitans* (n=28) and *S. hardwickei* (n=11), in peninsular India to determine whether their venoms are similar or dissimilar. An integrated proteo-transcriptomic approach was used to characterise the venom phenotypes of the two species across multiple individuals in peninsular India. We used species occurrence records and species distribution models to assess the distributional overlap among these species within the peninsular Indian region. The species showed significant overlap in their current and projected geographical ranges, corresponding with their co-occurrence. We characterised the venom profiles of both species and found that the venoms were cocktails of enzymes, β-pore-forming toxins, and neurotoxins comprising 110 and 84 proteins in *S. morsitans* and *S. hardwickei,* respectively. However, the venom composition of both species differed significantly in toxin abundance and species-specific protein repertoires. This indicates trait divergence in venom phenotypes, suggesting that distinct venom compositions may facilitate coexistence among ecologically similar predatory centipedes. The observed variation in venom phenotypes among co-distributed species opens up important avenues for future research into their ecological roles and functional significance. In this study, we provided a detailed account of venom composition across multiple individuals from the species’ geographic range and highlighted the importance of investigating the role of venom as a trait that could influence species interactions and shape communities in these diverse tropical forests.

## Introduction

Functional traits such as morphological, behavioural, physiological, or biochemical characteristics are crucial for determining species’ ecological roles and interactions, providing critical insights into the mechanisms underlying species coexistence (Chesson, 2000; Nock et al., 2016). Examples include beak size in birds (Navalón et al., 2019; Olsen, 2017), nasal bones in rodents (Martinez et al., 2018), and proboscis length in insects, which influence dietary specialisation and feeding strategies (Gilbert & Jervis, 1998). One key mechanism examined across taxa is how shared ecological pressures among coexisting species can lead to either convergence or divergence in functional traits. Coexisting species may be functionally more dissimilar than expected by chance due to niche differentiation (trait divergence). Conversely, trait convergence occurs when species living in the same habitat evolve similar phenotypes (Adler et al., 2013; Borics et al., 2020; De Solan et al., 2022; Grime, 2006).

Venoms, complex cocktails of bioactive molecules, including neurotoxic peptides and enzymes, represent a key adaptive trait that has evolved independently across diverse animal lineages (Schendel et al., 2019). As potent biochemical traits, venoms play crucial roles in predation, defence, and competition, influencing ecological interactions across taxa (Casewell et al., 2013; Fry et al., 2009). However, understanding the role of venoms in ecological interactions remains limited to relatively few studies. Theory suggests that predatory venoms enable smaller organisms to efficiently subdue larger or faster prey (Arbuckle, 2015; Casewell et al., 2013). At the same time, ecological competition may be mediated by venom specialisation, allowing coexisting species to exploit distinct dietary niches. While venom specialisation has been proposed as a mechanism facilitating ecological differentiation and coexistence, empirical evidence across taxa remains limited and inconclusive. Comparative studies reveal taxonomic variation in the drivers of venom differentiation. In spiders and parasitoid wasps, venom differentiation often reflects prey or host specialisation (Mathé-Hubert et al., 2019; Pekár et al., 2022). In snakes, venom variation is influenced by a combination of diet, environment, and ontogeny (Hirst et al., 2024; Holding et al., 2021; Zancolli et al., 2019). Cone snails exhibit clear venom divergence associated with dietary breadth (Weese & Duda, 2019). Despite these insights, the role of venom in species coexistence remains largely unknown in many venomous groups, especially in arthropods.

Although venom composition and function can now be precisely analysed using proteomic and transcriptomic tools (Mouchbahani-Constance & Sharif-Naeini, 2021; von Reumont et al., 2022), venoms remain understudied in many taxa. Challenges include the difficulty of extracting sufficient venom volumes for individual-level quantification and, importantly, the lack of related ecological data on species’ geographic ranges, abundances, interactions, and natural history, all of which are essential for studying the role of venoms in species coexistence (Sunagar et al., 2016; Von Reumont et al., 2014).

In this study, we assess venom composition in two coexisting centipede species in the tropical forests of Asia. Centipedes are among the oldest terrestrial venomous predators, and their venoms contain neurotoxins that immobilise prey, including ion channel modulators (targeting sodium, potassium, and calcium channels) critical for neuromuscular incapacitation; enzymes such as metalloproteases, hyaluronidases, and phospholipases that facilitate tissue penetration and digestion; and cytolytic toxins, including β-pore-forming toxins (β-PFTx) (Jenner et al., 2019; Liu et al., 2012; Undheim et al., 2014, 2015; Undheim & Jenner, 2021; Undheim & King, 2011). Scolopendrid centipedes are large-bodied, highly venomous arthropods with global distribution, playing crucial roles as apex invertebrate predators in soil ecosystems. Their ancient evolutionary history and ecological significance make them an excellent system for studying venom evolution and species interactions (Arbuckle, 2015, 2020).

We examine venom phenotypes in two coexisting and closely related centipede species, *Scolopendra morsitans* and *S. hardwickei*. Specifically, we test whether these species have similar venom compositions due to shared ecological pressures (e.g., prey types or habitats), or whether ecological competition (e.g., for diet) promotes divergence in venom composition. The genus *Scolopendra*, comprising approximately 100 species distributed worldwide, includes some of the largest centipedes (reaching up to 300 mm in length). These centipedes are significant predators in soil ecosystems and hold cultural and medicinal importance (Siriwut et al., 2015, 2016). *S. morsitans* is widely distributed across tropical forests in Africa, Australia, and South and Southeast Asia, with its type locality in India. *S. hardwickei*, commonly known as the Indian tiger centipede, is one of the largest species and is endemic to the Indian subcontinent. These species have been documented to be coexisting across habitats in peninsular India (Jangi & Dass, 1984).

Trait similarity or dissimilarity may be influenced by evolutionary history. To control for this, we first constructed a phylogenetic tree demonstrating that these species are evolutionarily closely related within the genus *Scolopendra*. Systematic and biogeographic work conducted on scolopendrid centipedes in peninsular India over the last decade has enabled us to quantify spatial overlap among these species. We built Species Distribution Models (SDMs) using extensive field survey data, supplemented by published records, to predict potential species distributions based on bioclimatic niches. While SDMs do not directly indicate species interactions, they can serve as a starting point for investigating potential coexistence in soil-dwelling predatory organisms, particularly where abundance data are scarce. *S. morsitans* and *S. hardwickei*, which have persisted in the landscape over long evolutionary timescales (Joshi & Karanth, 2011), exhibit overlapping predicted distributions that can be used as a proxy for coexistence across habitats in peninsular India.

These species exhibit similar body sizes, a key trait influencing resource use and ecological interactions among coexisting predators. Body size has been linked to variation in venom potency and functional composition across taxa. For example, larger snakes exhibit lineage-specific shifts in venom composition, with certain toxin classes becoming more prevalent rather than increasing in potency universally (Siqueira-Silva et al., 2021). Conversely, smaller scorpions with slender chelae exhibit markedly more potent venoms, reflecting an evolutionary trade-off between mechanical strength and venom investment (Forde et al., 2022).

Given their evolutionary relatedness, similar body sizes, and co-occurrence, *S. morsitans* and *S. hardwickei* provide an ideal setting to investigate the role of venoms in species coexistence among arthropod predators. By analysing venom composition across multiple individuals of both species, we assess patterns of similarity and divergence in venom phenotypes. We test whether habitat similarity has led to similar venom profiles or whether interspecific competition has driven venom divergence. We predicted that high similarity in venom phenotypes would indicate trait similarity, whereas significant differences would suggest divergence in response to ecological conditions.

To test this, we analysed venoms from multiple individuals of both species using proteomics. We also generated venom gland transcriptomes for each species. Proteomic analyses provided a direct representation of the expressed venom phenotype, while transcriptomic data were used to improve protein identification. An integrated approach like this is essential because proteomic analyses alone can miss certain venom components due to protein solubility constraints and ionisation efficiency, as reported in previous studies (Jenner et al., 2019; Madio et al., 2017; Smith & Undheim, 2018). Similarly, transcriptome-based venom profiling can sometimes yield inflated or incomplete estimates of venom diversity. Performing transcriptome analyses alone can overestimate toxin diversity because it fails to distinguish toxin sequences from non-toxin homologues. Also, assembly fragmentation due to limited sequencing depth can further obscure the true representation of expressed toxins (Jenner et al., 2019; Smith & Undheim, 2018). Therefore, an integrated proteo-transcriptomic approach was employed to accurately assess the venom composition of the two species.

## 2. Materials and Methods

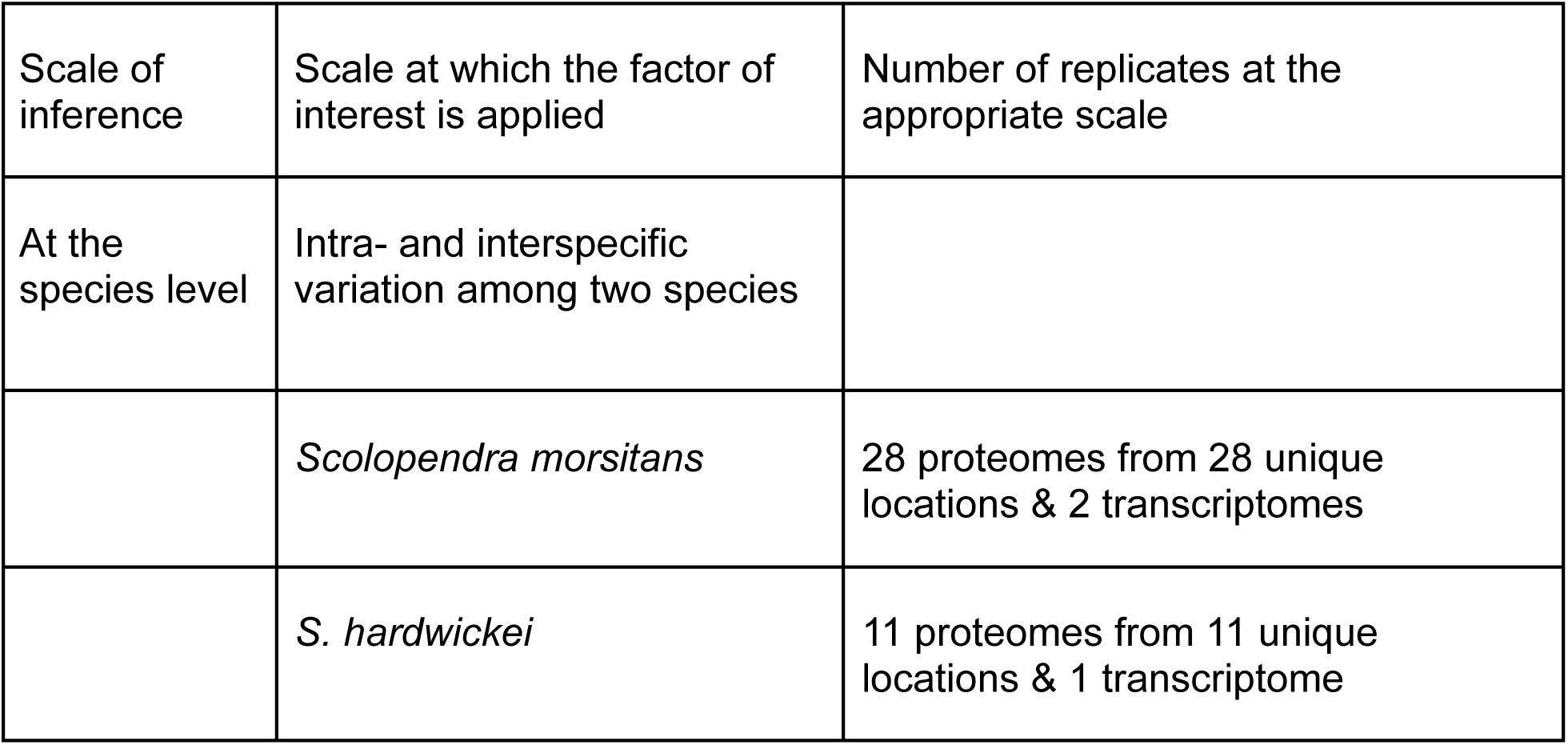

| Scale of inference | Scale at which the factor of interest is applied | Number of replicates at the appropriate scale |
| --- | --- | --- |
| At the species level | Intra- and interspecific variation among two species |  |
|  | *Scolopendra morsitans* | 28 proteomes from 28 unique locations & 2 transcriptomes |
|  | *S. hardwickei* | 11 proteomes from 11 unique locations & 1 transcriptome |

### 2.1. Taxon sampling, phylogenetic, geographic and climatic niche overlap analyses

#### Taxon sampling

*S. morsitans* and *S. hardwickei* individuals were collected from multiple populations (n = 59) during the July-September monsoon period (2021-2024) (Fig. 1). The sampling was targeted across the Western and Eastern Ghats of peninsular India in various habitat types, including wet forests, dry forests, and savannahs. 35 of 59 samples from both species were from the same locality, suggesting that these species co-occur and occupy similar habitats. The species occurrence data were collected through fieldwork, supplemented by published literature (Jangi & Dass, 1984; Joshi & Edgecombe, 2018; Joshi & Karanth, 2011; Rathinasabapathy, 1997; Vazifdar, 2021).

**Figure 1.**
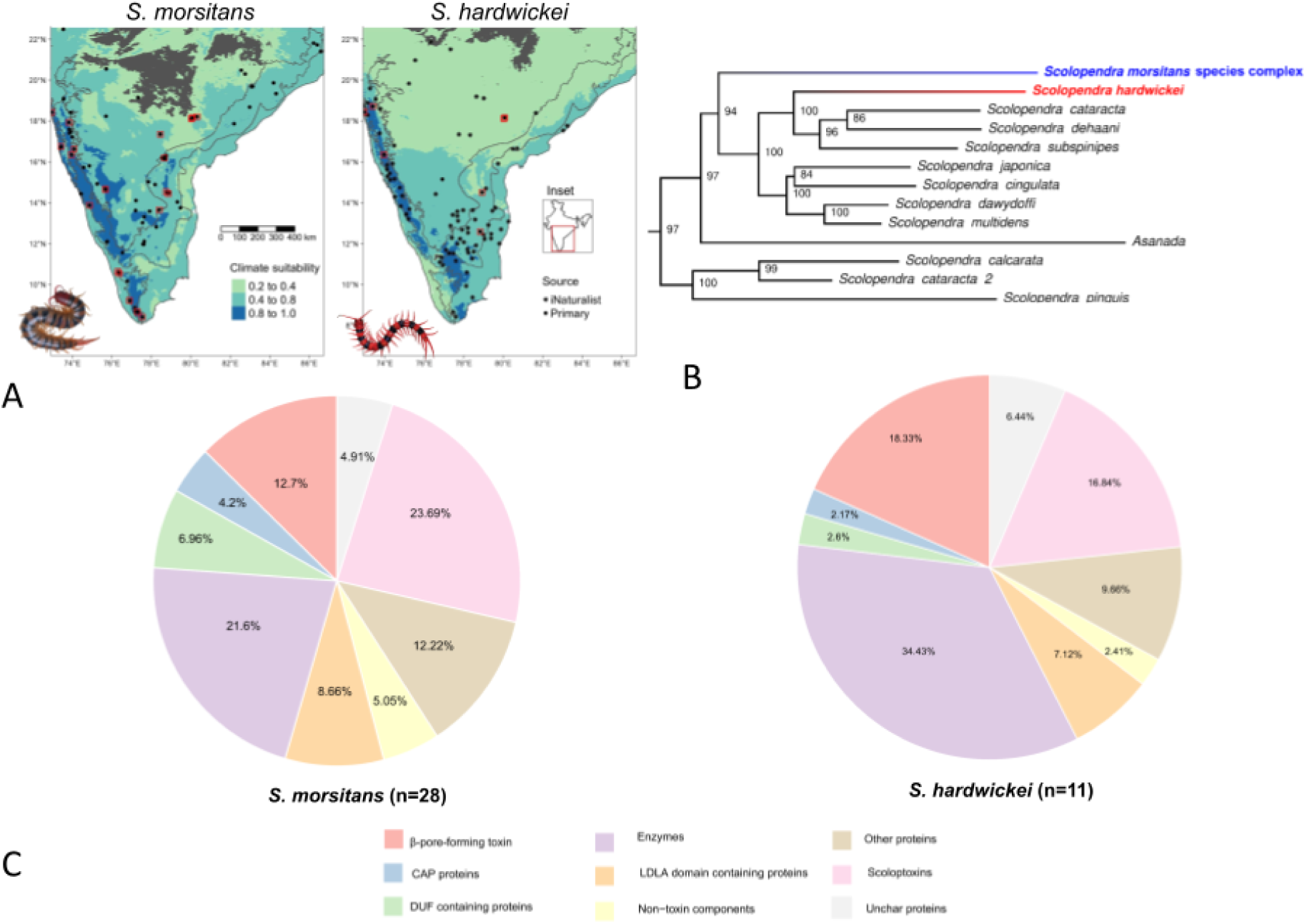
A. The habitat suitability and climate niche space occupied by *S. morsitans* and *S. hardwickei* in peninsular India. *Scolopendra* picture credits: Karunakar Majhi. B. Maximum Likelihood tree indicating the phylogenetic position of *S. morsitans* and *S. hardwickei,* numbers at each node indicate bootstrap support. C. Proteomic venom composition of *S. morsitans* (n=28) and *S. hardwickei* (n=11).

#### Molecular phylogenetic analyses

The molecular phylogenetic tree for tropical Asian *Scolopendra* species was recently reconstructed (Siriwut et al., 2016). However, *S. hardwickei* was missing from the dataset. Also, earlier phylogenetic analysis indicated paraphyly in *S. morsitans.* Therefore, we reconstructed the molecular phylogenetic tree, including *S. hardwickei* and extensive samples of *S. morsitans* across its range (n=54, Fig. 1). We used two mitochondrial DNA (mtDNA - COI & 16S) loci and one nuclear DNA (nuDNA) locus (28S) in a likelihood framework to ascertain their evolutionary relationships. Additionally, we performed a species delimitation analysis to identify species boundaries within *S. morsitans*. The details of the taxon sampling, tree reconstruction and species delimitation analyses are described in SI (Supplementary Information 1).

#### Geographical distribution

We used Species Distribution Models (SDMs) to determine the geographic range and overlap between *S. morsitans* and *S. hardwickei.* We mapped occurrences alongside predicted climatic suitability to visualise the relative extents of the two species across peninsular India. For *S. hardwickei*, we augmented the occurrence data with 192 research-grade records from iNaturalist. To reduce clustering, we thinned the occurrences by removing records within 20 km of one another using the R package spThin v0.2. Climate predictors for the model were variables that describe the mean and seasonality of temperature and precipitation across the peninsular Indian region. We obtained the climate variables from Worldclim (Fick & Hijmans, 2017) at a 30-arc-second resolution. We used MaxEnt, which models species distributions using the species’ known occurrence records and associated climatic variables. This allows prediction of suitable habitat across the landscape based on the environmental conditions at presence locations. It was implemented in the Enmeval v2.0 package (Kass et al., 2021) in R v4.4 to assess climatic suitability for both species, following the approach and parameters detailed in (Bharti et al., 2021) for scolopendrid centipedes of the Western Ghats, India (Fig. 1).

### 2.2. Characterising venom phenotype

By integrating proteomic and transcriptomic data, we characterised venom phenotypes at the individual level for both species. The live animals were brought to the lab, kept in a cool, dark place, and aired regularly for proper ventilation. Venom was extracted from 28 *S. morsitans* and 11 *S. hardwickei* individuals using a commercial Transcutaneous Electrical Nerve Stimulation (TENS) unit *via* electrostimulation and stored at −80 °C. All extractions were performed at the individual level. The venom glands were dissected 4 days after venom milking, stored in RNAlater at −80°C, and total RNA was extracted manually using the phenol-chloroform method (Jenner et al., 2019). We sequenced the transcriptomes for *S. morsitans* and *S. hardwickei* from a single adult individual of each species. The assembled and annotated venom gland transcriptomes were used as reference sequence databases for protein identification in proteomic analyses.

#### Transcriptome generation and assembly

mRNA sequencing was performed at the Central NGS facility of the Centre for Cellular & Molecular Biology, Hyderabad, India, using the Illumina NovaSeq 6000 platform. The raw reads were processed with quality checks (FastQC) before and after trimming low-quality reads (Phred score < 30) and adapter sequences using TrimGalore version 0.6.6. Trimmed reads were assembled into transcripts using the Trinity assembler, which reconstructs transcriptomes *de novo* through Inchworm, Chrysalis, and Butterfly modules to generate full-length transcripts by capturing transcript sequence diversity, including alternative splicing and isoforms (Grabherr et al., 2011). Finally, Cd-hit was used to reduce redundancy within the assembled transcriptomes (Li et al., 2001, 2002; Li & Godzik, 2006). We used Kallisto to quantify gene expression levels in the venom gland transcriptomes, estimating transcript abundances in Transcripts Per Million (TPM) (Bray et al., 2016). *De novo* assembled and annotated venom gland transcriptomes were used as reference databases for mass spectrometry scan searches. The annotated transcriptomes were first translated *in silico* using the Galaxy tool “Get ORF” (Evans et al., 2012; Rice et al., 2000) with an open reading frame (ORF) cut-off of 50 and default settings.

#### Venom proteomics

We employed a bottom-up approach to identify venom proteins (Jenner et al., 2019). The venom proteomes of 28 individuals of *S. morsitans* and 11 of *S. hardwickei* were generated in the Central Proteomics facility, CCMB, Hyderabad, India. After the in-solution digestion of venom proteins, we analysed the generated peptides using the Thermo Scientific Easy-nLC 1200 system coupled to a Q-Exactive mass spectrometer (QE). Peptides were resolved on a Thermo Scientific PepMap RSLC (rapid separation liquid chromatography) C18, 3μm, 100 Å, 75 μm x 15 cm column (ES900) using a 5% - 25% - 45% - 95% −95%-3% Solvent B gradient followed at 0.00, 35.00, 44.00, 50.00, 55.00 and 60.00 minutes respectively (Solvent A: 5% acetonitrile-0.2% formic acid; Solvent B: 90% acetonitrile-0.2% formic acid), in DDA mode (Data Dependent Mode). The nLC eluent was then sprayed into the QE with a spray voltage of 2.2 kV (kilovolts). The mass spectrometer was operated in positive ion mode, with the following MS1 settings: scan range, 400-1650 m/z; automatic gain control (AGC) target, 3 × 10^6^; resolution, 70,000; and a filter for the minimum charge state, set to two. The MS2 data were acquired with the following set of parameters: resolution, 17,500; normalised collision energy, 30; topN, 10; and peptides with charge states 2-5. The resulting data was visualised in Thermo Xcalibur Qual B 3.0.63. Raw data were processed using Proteome Discoverer v2.2.0.388 (Thermo Scientific). Spectra were searched against UniProt databases and custom protein databases derived from de novo-assembled venom gland transcriptomes generated in this study. The following parameters were set for each run: enzyme name: Trypsin (full), minimum peptide length: 6, maximum peptide length: 144, maximum missed cleavages allowed: 2, precursor mass tolerance: 15 ppm, fragment mass tolerance: 0.02 Da, lowest charge state: 2, highest charge state: 6, target FDR strict 0.01, relaxed 0.05.

Carbamidomethylation of cysteine (+57.021 Da) was taken as the static modification, and oxidation of methionine (+15.995 Da) was taken as the dynamic modification. All other parameters were set to their default values.

#### Defining venom phenotypes

Proteins identified with medium and high confidence were retained for downstream analyses. Protein identifications were validated by their presence in the proteomic dataset, using false discovery rate (FDR) and cross-correlation (Xcorr) score thresholds as implemented in Proteome Discoverer v2.2.0.388 (Thermo Fisher Scientific). These criteria ensured the inclusion of proteins supported by high-quality peptide-spectrum matches and minimised the risk of false positives. We used Jaccard’s and Bray-Curtis dissimilarity indices to compare the venom profiles of the two species. Jaccard’s dissimilarity index measured differences in venom composition based on the presence or absence of proteins, while the Bray-Curtis dissimilarity index accounted for the abundance of venom proteins. The differences were tested with the Kruskal-Wallis test, followed by Dunn’s test, which is the post-hoc test for pairwise comparisons. We employed ordination methods, including non-metric Multidimensional Scaling (nMDS) and Principal Component Analysis (PCA), to assess whether venom protein composition and abundance differ between the two species. PCA, which uses Euclidean distance, was applied to identify the major axes of variation in venom phenotypes, while nMDS preserved pairwise dissimilarities using Bray-Curtis distance, providing a more detailed visualisation of similarities and differences in venom composition. We then used permutational ANOVA (permanova) and ANOSIM to assess the significance of venom profile patterns. Pie charts were generated to depict the venom phenotypes of both species based on cumulative protein copy numbers across broad categories, including enzymes, scoloptoxins (SLPTX), β-pore-forming toxins (β-PFTx), LDLA domain-containing proteins, CAP proteins, DUF-containing proteins, uncharacterised proteins (UNCHAR), other proteins, and non-toxin proteins.

## 3. Results

### Venom gland transcriptomes of *S. morsitans* and *S. hardwickei*

The total raw reads processed to generate the assembled transcriptomes were 50,240,329 and 36,447,924 to generate 223,104 (N50=619) and 111,540 (N50=2191) transcripts for *S. morsitans* and *S. hardwickei*, respectively (Table 1). Transcriptomic data were primarily used as reference databases for proteomic analyses. However, we also examined expression patterns using Kallisto to generate the venom gland transcriptional profiles. Among the highly expressed transcripts (TPM ≥ 1000) in the venom gland of *Scolopendra morsitans,* the abundance was dominated by non-toxin genes associated with structural and metabolic functions, including troponin, tropomyosin, actin, myosin, and arginine kinase. A few toxin-related transcripts, such as SLPTX15, CAP proteins, and γ-glutamyl transpeptidase (GGT), were also detected among the highly expressed transcripts. In *Scolopendra hardwickei*, a similar trend was observed, with high expression of muscle-associated proteins, including troponin, tropomyosin, and actin, as well as metabolic enzymes such as arginine kinase and fructose-bisphosphate aldolase. Among toxin-related transcripts, several scoloptoxins (SLPTX01, SLPTX14, SLPTX17, SLPTX28) and β-pore-forming toxins (BPFTx), as well as γ-Glutamyl transpeptidases (GGT), were present within the highly expressed range. These patterns reflect general transcriptional activity in venom gland tissues.

**Table 1.** Summary of venom gland transcriptome assembly statistics and transcript expression categories for *Scolopendra morsitans* and *S. hardwickei*. TPM = Transcripts Per Million; ORFs = Open Reading Frames.

| Parameter | *S. morsitans* | *S. hardwickei* |
| --- | --- | --- |
| Raw reads processed | 50,240,329 | 36,447,924 |
| Total assembled transcripts | 223,104 | 111,540 |
| N50 (bp) | 619 | 2,191 |
| Total predicted ORFs (in silico translated proteome) | 3,370,904 | 2,712,876 |
| Rare ($\leq 10$ TPM) | 215,081 (Avg. TPM = 1.806) | 104,855 (Avg. TPM = 1.911) |
| Low ($>10 - <100$ TPM) | 7,064 (Avg. TPM = 27.068) | 5,832 (Avg. TPM = 25.808) |
| Moderate ($>100 - <1000$ TPM) | 890 (Avg. TPM = 241.757) | 719 (Avg. TPM = 277.790) |
| High ($\geq 1000$ TPM) | 69 (Avg. TPM = 2,971.015) | 134 (Avg. TPM = 3,353.519) |

### Venom proteomics

The venoms were characterised by a greater proportion of high molecular weight proteins ≥ 10kDa (*S. morsitans*: *μ*=172±34.35; *S. hardwickei: μ*=140±37.95) than low molecular weight proteins (*S. morsitans μ*=35±10.70; *S. hardwickei μ*=29±6.09). On average, *S. morsitans* (*μ*=49±8.37; *μ*=183.35±34.09) exhibited higher venom toxin richness and abundance compared to *S. hardwickei* (*μ*=41±7.72; *μ*=146.81±33.26). Of 95 identified toxin proteins, 24 were unique to *S. morsitans*, while four were found uniquely in *S. hardwickei* (Supplementary Table 3). This suggests that more species-specific toxin proteins are detected in *S. morsitans* than in *S. hardwickei,* based on proteomic analyses.

#### Venom phenotypes of *S. morsitans* and *S. hardwickei*

Species distribution models showed that *S. morsitans* and *S. hardwickei* had overlapping distributions with high predicted suitability (p > 0.8) in the central and northern Western Ghats (Fig. 1). Intraspecific venom variation was lower than interspecific according to Jaccard’s dissimilarity index (H-statistic=190.376, df=2, *p*-value <0.001, Fig. 2), based on the unique proteins’ presence. Similarly, intraspecific comparisons using the Bray-Curtis dissimilarity index had lower values than interspecific comparisons (H-statistic = 314.220, df = 2, *p*-value < 0.001), which incorporated both proteins and their abundance. Interspecific comparisons exhibited higher mean dissimilarity values, reflecting distinct venom compositions between *S. morsitans* and *S. hardwickei*. Also, the variance in interspecific distances was lower than within-species variance, indicating that although the two species are compositionally well separated, individuals within each species were also variable. Dunn’s post-hoc test with correction for the indices revealed that *S. hardwickei* and *S. morsitans* venom composition profiles were significantly different from each other (*p* = 0.001; Fig. 2). Non-metric multidimensional scaling (nMDS) based on Bray–Curtis distances revealed distinct patterns in venom composition between *S. morsitans* and *S. hardwickei* (stress = 0.155; non-metric fit R² = 0.9762; linear fit R² = 0.907). There was a significant difference in venom composition as indicated by permANOVA between *S. morsitans* and *S. hardwickei* (pseudo-F=15.36, p=0.001) and ANOSIM (R=0.42, p = 0.001). In the PCA plot, the venom profiles did not overlap, with PC1 explaining 31.5% and PC2 28.5% of the variation. The PC1 loadings were highest for β-PFTx (0.57), followed by enzyme GGT (0.40) and CUB domain-containing metalloproteases (0.31), indicating that the two species diverge significantly in the relative abundance of these toxin proteins. PC2 was dominated by SLPTX15 (0.46), SLPTX11 (0.37), and β-PFTx (0.37) (Supplementary Table 4).

**Figure 2.**
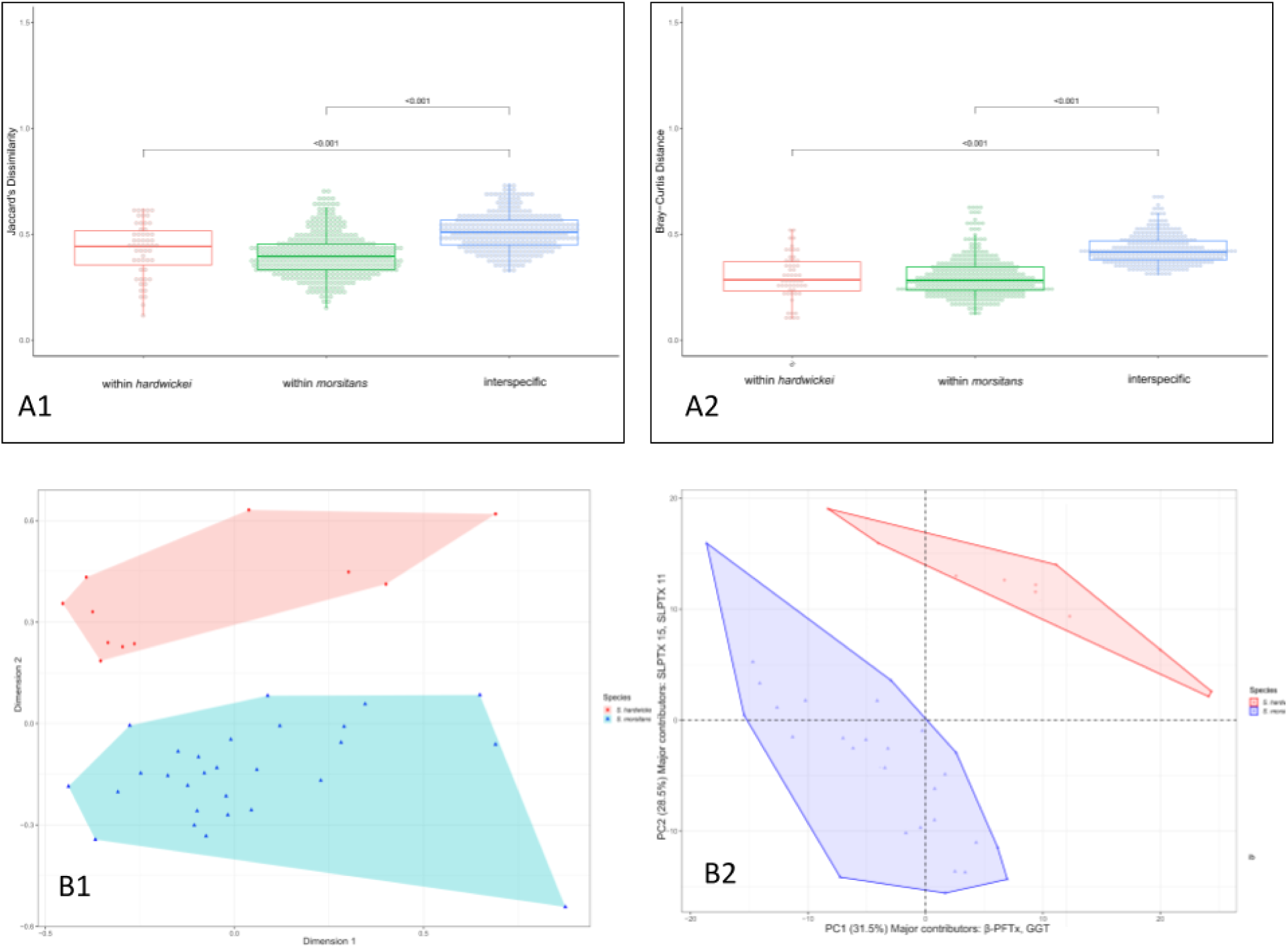
(A1) Jaccard’s dissimilarity and (A2) Bray–Curtis distance comparisons of venom protein profiles within and between *Scolopendra morsitans* (n = 28) and *S. hardwickei* (n = 11). (B1) Non-metric Multidimensional Scaling (nMDS) and (B2) Principal Component Analysis (PCA) plots showing clustering of venom compositions by species. Blue indicates *S. morsitans* and red indicates *S. hardwickei*. In both indices, intraspecific dissimilarity values were significantly lower than interspecific values (p < 0.001), indicating distinct venom compositions between species.

## Discussion

Our comparative analysis of venom phenotypes in *Scolopendra morsitans* and *S. hardwickei* provides one of the first proteomic analyses of centipede venoms from the tropical forests of India and includes the first comprehensive venom profile for *S. hardwickei*. Despite their close evolutionary relationship, overlapping distributions, and similar body size, the two species exhibit distinct venom phenotypes. These differences are primarily driven by variation in toxin abundance and the presence of species-specific proteins. Notably, *S. morsitans* had a more complex venom profile, with 24 unique toxin proteins, compared with just 4 in *S. hardwickei* (Supplementary Table 3). This difference in venom composition could help minimize interspecific competition (Pfennig & Pfennig, 2009, 2010; Stuart et al., 2017), a hypothesis that could now be further tested. Both species shared a core set of venom components, with an approximate 65.5% similarity in protein presence/absence, but differed significantly in their relative abundance and diversity. This could be due to species-specific ecological strategies, such as dietary preferences or habitat use, that likely contributed to their functional divergence, which could be studied further. Our species distribution models also indicate a broader geographic niche for *S. morsitans* than for *S. hardwickei*.

### Compositional and functional diversity of *Scolopendra morsitans* and *S. hardwickei* venoms

Proteo-transcriptomic analysis revealed a complex venom cocktail in both *S. morsitans* and *S. hardwickei*, dominated by enzymes, β-pore-forming toxins (β-PFTx), and scoloptoxins, highlighting the importance of cytolytic and neurotoxic functions in prey capture (Table 2). The β-PFTx is a major non-enzymatic component of centipede venom (Fig. 1C), originating from the bacterial aerolysin-like β-pore-forming toxin superfamily, known for forming transmembrane pores that induce cytolysis (Podobnik et al., 2017; Undheim et al., 2015; Undheim & Jenner, 2021). Their presence in centipede venoms is thought to be facilitated by horizontal gene transfer from bacteria, at least twice (Undheim & Jenner, 2021). These toxins are known to cause myotoxic and edematic effects (Malta et al., 2008), and their dominance across multiple *Scolopendra* species (*S. viridis, S. polymorpha, S. heros*) suggests a conserved functional role (Ellsworth et al., 2024). β-PFTx also had the highest loading in PC1, indicating that interspecific differences are driven mainly by variation in its abundance (Supplementary Table 4).

**Table 2:** Dominant toxin proteins found in the venoms of *Scolopendra* (Undheim et al., 2015; Undheim & King, 2011)

| Protein name | Type | Function |
| --- | --- | --- |
| SLPTX 15 | 4–6 cysteines<br>C–C–CxC | Unknown; K<sub>v</sub> channel antagonists (KC144556); NaV antagonists (KC144793); CaV channel antagonists (KC145039) |
| SLPTX 11 | 4–18 cysteines<br>C–C–CxC–C–C–<br>C–CxC–<br>C–C–C–CxC–C–<br>C (e.g.,<br>KC144104);<br>C–CxC–C (e.g.,<br>JN646116) | Unknown; KV channel antagonists (e.g., JN646116, KC144104); Anticoagulant (KC144430) |
| SLPTX 04 | 4 cysteines;<br>transcripts may<br>encode additional<br>linear peptides<br>upstream of<br>cysteine-rich<br>peptide<br>C–C–C–C | Unknown; KV channel antagonist (KC144226); putative synergistic mode of action for peptides encoded by multidomain transcripts (e.g., U-SLPTX4-Er1.1 and U-SLPTX4-Er1.2 from KF130724) |
| β-PFTx | β-Pore-Forming<br>Toxin | Potentially cytotoxic *via*<br>formation of polymeric pore |
|  |  | structures in cell membranes |
| GGT | γ-Glutamyl<br>transpeptidases | Platelet aggregating activity,<br>hemolytic to mouse and rabbit<br>hemocytes |
| LDLA | LDLA-repeat<br>domain containing<br>protein | Unknown, receptor-mediated<br>endocytosis of specific ligands<br>(Daly et al., 1995)<br>Moderation of immune<br>responses in crustaceans<br>(Huang et al., 2014)<br>Antibacterial (Qu et al., 2016)<br>Antiviral function in shrimps (Xiu<br>et al., 2015, 2016) |
| CentiCAP2 | cysteine rich<br>secretory proteins<br>(CRISPs), antigen<br>5 (Ag5) and<br>pathogenesis-rela<br>ted 1 (Pr-1)<br>proteins (Ward &<br>Rokyta, 2018) | CaV channel antagonist<br>(KC144967);<br>Trypsin inhibitor (KC144061) |

Scoloptoxins, another key non-enzymatic group, are cysteine-rich neurotoxins that modulate voltage-gated Na⁺, K⁺, Ca²⁺, and TRP channels, contributing to prey paralysis, pain, and systemic disruption (Ellsworth et al., 2024; Luo et al., 2022). Some also function as chitinases, aiding in digestion and possessing pharmacological potential (Ellsworth et al., 2024; Hu et al., 2023). SLPTX11 and SLPTX15 showed high loadings on PC2, indicating their importance in distinguishing species-specific venom phenotypes.

CAP proteins, including CRISPs, antigen 5, and PR-1 proteins, were abundant in both venoms. Though their functions in centipedes remain unclear, they are known in other taxa to act as ion channel modulators, vasodilators, and allergens (Fry et al., 2009; Gibbs et al., 2008; Moran et al., 2013). Their high expression levels suggest they may contribute to envenomation symptoms, such as inflammation and hypersensitivity.

CUB domain-containing metalloproteases, zinc-dependent enzymes found in many venomous taxa, were also present. In snakes, they induce haemorrhage and apoptosis, and aid venom spread by degrading the extracellular matrix (Cerdà-Costa & Xavier Gomis-Rüth, 2014; Gutiérrez, 2000). Their abundance in both species and possible role in symptoms such as edema and skin damage (Undheim et al., 2014, 2015; Undheim & King, 2011) highlight their functional importance in centipede venoms. Interestingly, their abundances also differed between the species.

γ-Glutamyl transpeptidases (GGTs) are cell surface enzymes that play a crucial role in oxidative stress regulation and xenobiotic detoxification by catalysing the transfer of γ-glutamyl moieties to amino acids (Danneels et al., 2010). These enzymes have been implicated in hepatocyte damage in snakes (Al-Quraishy et al., 2014) and liver toxicity in bees following venom injection (Ivas et al., 2011). GGT induces apoptosis in host ovaries in parasitoid wasps through oxidative stress (Falabella et al., 2007). They also contribute to inflammation, swelling, nausea and fever due to their effects on hepatic tissues. GGT are widely present in centipede venoms, reported from species such as *S. subspinipes dehaani*, *S. morsitans*, and *T. longicornis*. In *S. subspinipes dehaani*, GGT has been shown to induce platelet aggregation and red blood cell hemolysis (Liu et al., 2012). Their widespread occurrence in scolopendrid venoms suggests they play a key role in envenomation, likely contributing to oxidative stress, allergic responses, and cell death. We also found GGT at high abundance in both species, although the numbers were distinct between them.

Interestingly, we find a high number of LDLA domain-containing proteins, DUF-containing proteins, and uncharacterised proteins (UNCHAR) in the venoms of both *Scolopendra* species. Despite their undefined functions, their high expression levels in both venom gland transcriptomes and venom proteomes suggest a possible role in venom and/or gland maintenance, which warrants further exploration. We also found that *Scolopendra* venoms were dominated by proteins with molecular weights ≥ 10 kDa, a pattern previously reported for other *Scolopendra* species (Liu et al., 2020) and these proteins could be potential therapeutic targets (Han et al., 2022; Yadav & Upadhyay, 2022). Although most components of centipede venom remain functionally uncharacterized, we assigned putative functions to the identified proteins based on homologies, using BLAST searches and literature surveys. The integration of proteomic and transcriptomic data was crucial for identifying isoform diversity and improving functional annotations, especially for poorly represented taxa like centipedes.

### Distinct venom phenotypes among coexisting *Scolopendra* species

The divergence in venom composition between *Scolopendra morsitans* and *S. hardwickei* likely reflects ecological niche differentiation driven by factors such as diet, microhabitat use, or interspecific competition. Yet it is also possible that such compositional divergence arose from neutral or environmentally mediated processes, as co-occurring species frequently experience similar abiotic conditions without necessarily interacting directly (Blanchet et al., 2020). Functional differentiation may arise even in the absence of strong biotic interactions, highlighting the need for future experimental or dietary studies to directly test the ecological consequences of venom variation. While the two species shared ∼ 65-70% similarity in venom protein richness, the relative abundance of shared toxins varied substantially, potentially influencing venom potency and prey specificity. For instance, variation in the copy number of β-pore-forming toxins (β-PFTx) suggests quantitative differences in venom functionality.

*S. morsitans* exhibited a more complex venom profile, with 24 unique toxin proteins compared to just four in *S. hardwickei*, ‘unique’ referring to proteins exclusively detected in one or more individuals of one species (Supplementary table 3). Despite this diversity, ∼90% of the venom phenotype in both species was driven by the abundance of just 30% of the total detected proteins, indicating that a few highly expressed toxins dominate venom composition. Notably, toxin classes such as β-PFTx and CUB domain-containing metalloproteases showed extensive isoform diversity, consistent with patterns of functional streamlining, in which key toxin classes are selectively optimised for specific ecological roles and are prominently represented within venom phenotypes. This mirrors trends seen in snakes, where reduced inter-class redundancy is offset by intra-class diversification (Jackson et al., 2016).

This divergence could also reflect character displacement, an evolutionary mechanism promoting trait divergence between closely related species in sympatry to minimise competition and reproductive interference (Pfennig & Pfennig, 2009, 2010). Similar patterns have been observed in *Crotalus* and *Micrurus* snakes, as well as *in Conus* cone snails, in which venom variation is thought to be an adaptive response to ecological overlap (Barghi et al., 2015; Smith et al., 2023). In our study, while *S. morsitans* and *S. hardwickei* share a core venom proteome, differences in expression levels may reflect variation in venom composition between species, consistent with patterns of trait divergence rather than complete compositional turnover.

### The value of the proteo-transcriptomic approach for non-model organisms

Our study emphasises the significance of integrating transcriptomic and proteomic data to precisely characterise venom composition in non-model organisms. While RNA-seq is effective for identifying toxin genes, it can introduce challenges such as false positives, the inability to detect post-translational modifications, and the misrepresentation of functional protein diversity. Homology-based methods (e.g., BLAST) may also overestimate venom complexity by mis-annotating non-toxin homologs as venom components (Smith & Undheim, 2018), especially in taxa where toxins evolve from non-toxic ancestral proteins. Proteomic analyses provide direct evidence of expressed venom components but can be limited by the representation in databases, especially in taxa such as centipedes that are underrepresented in resources like UniProt (Sunagar et al., 2016; Von Reumont et al., 2014). In our study, only 2.12% of *S. hardwickei* and 1.22% of *S. morsitans* proteins matched UniProt entries. Nevertheless, species-specific transcriptomes greatly enhanced toxin detection; for example, only 9 SLPTX proteins were identified by proteomics alone, whereas 26 were detected using the integrated approach. The proteo-transcriptomic approach was particularly effective for uncovering isoform diversity within major toxin classes, such as SLPTX. This integrated approach is helpful for accurately characterising venoms in understudied lineages, where reliance on a single method may overlook key components of functional and evolutionary significance.

## Conclusion

The first comparative characterisation of venoms from *S. morsitans* and *S. hardwickei* reveals species-specific venom phenotypes shaped by differential protein expression and possibly driven by ecological niche differences. These findings emphasise the adaptive importance of venom variation in influencing interspecific trait divergence, underlining the ecological and evolutionary significance of venoms in arthropods.

## Ethical statement

The authors declare that no ethics clearance was required for this study.

## Declaration of generative AI in scientific writing

The authors declare that no generative AI was used in the writing of this manuscript.

## CRediT authorship contribution statement

Aditi: Conceptualisation, Data curation, Formal analysis, Investigation, Methodology, Visualisation, Writing – review & editing.

Jahnavi Joshi: Conceptualisation, Supervision, Funding acquisition, Resources, Writing – review & editing.

Pragyadeep Roy: Data curation, Formal analysis, Investigation, Methodology, Visualisation, Resources

Richard Parikh: Data curation, Formal analysis, Methodology

Karunakar Majhi: Data curation, Formal analysis, Methodology

## Data Availability

All data used for analyses are presented in the supplementary material and are explained in detail in the methodology section. Also, available here - https://doi.org/10.5281/zenodo.17785917.

## Acknowledgements

We would like to thank Dr. D. K. Bharti, Dr. Mihir Kulkarni, Nehal Gurung, Pooja Pawar, Abhishek Gopal, Maya Manivannan, Payal Dash, Sudhanshu Kumar and Marwa Razak for their assistance with field and laboratory work. We thank Komal Verma, Deep, Sharvani Deshpande, Shriya, Shivika, Diya, SS Dhanush for helping with the venom extractions. We thank Dr Eivind Undheim for sharing his annotated centipede protein database for proteome analyses. We thank Tulasi Nagabandi, Md. Jafurulla and all the team members at the Central Next Generation Sequencing facility, Centre for Cellular & Molecular Biology (CCMB), Hyderabad (Telangana, India). We also thank Y Kameshwari, B Raman, and K Ranjith Kumar for their guidance and support during our work at the Central Proteomics facility at the Centre for Cellular & Molecular Biology (CCMB), Hyderabad, India. We thank Onkar Kulkarni and Payel Mukherjee for their help with transcriptome analyses. We also thank state forest departments and their staff for their help and support throughout fieldwork (Andhra Pradesh - 21049/1/2024 /WL-2, Tamil Nadu - WL5/AV12865/2022, Kerala - KFDHQ-2030/2021-CWW/WL-10, Karnataka - PCCF(WL)/E2/CR-08/2021-22, Maharashtra - CR-29(21-22)/1260/22-23, and Odisha 4637/4WL-413/2023).

