## supplementary files for "Divergent venoms among two closely related co-distributed centipede species, *Scolopendra morsitans* and *S. hardwickei* in tropical Asia"

**This PDF includes:**

Supplementary Information 1

Supplementary Information 2

Supplementary Tables 1 to 3

**Supplementary Information** 1

**Phylogenetic and species delimitation analysis of the genus *Scolopendra***

Taxon sampling and DNA data generation: We sampled *Scolopendra morsitans* and *S. hardwickei* species from the Western and Eastern Ghats across peninsular India. After venom extraction, these specimens were preserved in 70% Ethanol. Genomic DNA was extracted from either leg or segment tissue of each specimen using the standard protocol provided in the QIAGEN DNeasy Blood & Tissue Kit (QIAGEN, Germany). Two mitochondrial markers, COI and 16S rDNA, and a nuclear marker, 28S rDNA, were amplified using the PCR protocol described by Joshi & Karanth (2011). The forward and reverse DNA sequences of each marker were visualised and cleaned using the Chromas 2.6.6 software, based on a close visual inspection of the chromatogram peaks. A consensus sequence was obtained from the forward and reverse sequences by aligning them with MUSCLE in MEGA12 (Tamura et al., 2021; Kumar et al., 2024).

Molecular Phylogenetic Analyses: The molecular phylogenetic tree of *Scolopendra morsitans* and *Scolopendra hardwickei,* along with other *Scolopendra* species found across their ranges, was reconstructed using a concatenated dataset of two mitochondrial DNA (mtDNA) and one nuclear DNA (nuDNA) loci in a Maximum Likelihood (ML) and Bayesian Inference (BI) framework. We used a command-line version of IQTree2 with the default number of ultrafast bootstrap replicates. ModelFinder was used to determine a substitution model for the partitioned concatenated dataset (Minh et al., 2020; Trifinopoulos et al., 2016; Kalyaanamoorthy et al., 2017; Hoang et al., 2018). MrBayes 3.2.7a was used for Bayesian phylogenetic analyses (Huelsenbeck et al., 2001). The concatenated dataset of 16s-28s-COI loci was partitioned into five blocks: one for the 16s gene, one for the 28s gene, and three for each COI codon position. We identified appropriate substitution models for each partition block using ModelFinder in IQTree2, specifying a command to restrict the search to models supported by MrBayes. ModelFinder chose the following models based on the Bayesian Information Criterion (BIC): GTR + I + G for the 16S and first codon position of the COI gene block, GTR + G for the second and third codon positions of the COI gene blocks, and SYM + I + G for the 28S gene block. We performed two runs of 300 million generations, sampling every 500 generations, and stopped the runs once the standard deviation of split frequencies reached below 0.008. We then pruned the first 25% of trees to obtain the consensus tree.

The phylogenetic trees generated by ML and BI were similar, so the ML tree was used for further species delimitation analyses. A Scolopendrinae subfamily-level dataset comprising several species from the genera *Scolopendra*, *Asanada*, and *Cormocephalus* was used for phylogeny reconstruction, with the subfamily *Otostigminae* serving as the outgroup to root the tree.

The sequence dataset was obtained from two sources: DNA sequences generated in the lab (n=23) and DNA sequences available in the NCBI public repository. The ML trees were constructed from the individual datasets of the COI and 16S loci, as well as concatenated datasets of 16S-COI (n = 189) and 16S-28S-COI (n = 189). The single-locus tree-based approach to species delimitation, mPTP (multi-rate Poisson Tree Processes), was used to delineate the species boundaries of S. morsitans, given that it may be a species complex (Kapli et al., 2017). Species delimitation was performed on both individual mtDNA and concatenated mtDNA datasets. Upon reconstructing the subfamily-level ML tree, the subclade of the *S. morsitans* species complex was extracted from it. Correspondingly, the multiple sequence alignment of only *S. morsitans* was generated for mPTP, performed on the command line of Ubuntu Linux. A posterior MCMC support of 1 was used to get a conservative estimate of the distinct units in the *S. morsitans* species complex. The results from mPTP were mapped onto the ML tree of the concatenated 16s-28s-COI loci dataset visualised in the FigTree(v1.4.4) software.

### Supplementary information 2

| Sr. No. | ID | Genus | Species | COI | 16s | 28s | Location | Reference |
|---|---|---|---|---|---|---|---|---|
| 1 | CES091025 | *Ethmostigmus* | *praveeni* | MH908724 | MH908688 | MH908705 | Sharavathi Wildlife Sanctuary, Karnataka, India | Joshi & Edgecombe, 2018 |
| 2 | CES091335 | *Ethmostigmus* | *agasthyamalaiensis* | MH908729 | MH908697 | MH908712 | Kottavasal Reserve Forest, Kerala, India | Joshi & Edgecombe, 2018 |
| 3 | CUMZ 00432 | *Rhysida* | *longipes* | MF167833 | MF167766 | MF167900 | PK-2, Bahin village, Myanmar | Siriwut et al., 2018 |
| 4 | CES091082 | *Rhysida* | *sp.* | MK273266 | MK273384 | MK273486 | Rosemla forest, Shendurney Wildlife Sanctuary, Kerala, India | Joshi & Edgecombe, 2018 |
| 5 | CUMZ 00530 | *Otostigmus* | *scaber* | MF167802 | MF167735 | MF167869 | Wat Ban Hu, Luang Phrabang, Laos | Siriwut et al. 2018 |
| 6 | CUMZ 00518 | *Otostigmus* | *aculeatus* | MF167796 | MF167729 | MF167863 | Wat Phra-Ong Thom, Siam Riep, Cambodia | Siriwut et al. 2018 |
| 7 | CES091033 | *Digitipes* | *barnabasi* | JX531868 | JX531738 | JX531808 | Periyar Tiger Reserve, Kerala, India | Joshi and Karanth 2012 |
| 8 | CES091088 | *Digitipes* | *coonoorensis* | JX531880 | JX531750 | JX531817 | Pandimatte, Shendurney Wildlife Sanctuary, Kerala, India | Joshi and Karanth 2012 |
| 9 | MCZDNA104671 | *Hemiscolopendra* | *marginata* | HQ402548 | HQ402496 | - | Virginia, USA | Vahtera et al. 2012 |
| 10 | CCMB41 | *Scolopendra* | *morsitans 1c* | + | + | + | Nehru Zoo, Telangana | This study |
| 11 | CCMB752 | *Scolopendra* | *morsitans 1a* | + | + | + | Panchgani Plateau no. 2, Maharashtra | This study |
| 12 | CCMB803 | *Scolopendra* | *morsitans 1c* | + | + | + | Ranebennur WLS, Karnataka | This study |
| 13 | CCMB805 | *Scolopendra* | *morsitans 1c* | + | + | + | Ranebennur WLS, Karnataka | This study |
| 14 | CCMB818 | *Scolopendra* | *morsitans 1c* | + | + | + | Sateri Hill near Kolhapur, Maharashtra | This study |
| 15 | CCMB845 | *Scolopendra* | *morsitans 1c* | - | + | + | Ranebennur WLS, Karnataka | This study |
| 16 | CCMB1245 | *Scolopendra* | *hardwickei* | + | + | + | Lankamalla, Andhra Pradesh | This study |
| 17 | CCMB1325 | *Scolopendra* | *morsitans 1b* | + | + | + | Horsley Hills, Andhra Pradesh | This study |
| 18 | CCMB1696 | *Scolopendra* | *morsitans 1c* | + | + | + | Laknavaram, Mulugu division, Telangana | This study |
| 19 | CCMB1725 | *Scolopendra* | *morsitans 1c* | - | + | + | Tadvai, Telangana | This study |
| 20 | CCMB1766 | *Scolopendra* | *morsitans 1c* | + | - | - | Sarvapur beat, Mulugu range, Telangana | This study |
| 21 | CCMB1769 | *Scolopendra* | *morsitans 1c* | + | + | + | Sarvapur beat, Mulugu range, Telangana | This study |
| 22 | CCMB1776 | *Scolopendra* | *hardwickei* | - | - | + | Sarvapur beat, Mulugu range, Telangana | This study |
| 23 | CCMB2197 | *Scolopendra* | *hardwickei* | + | + | + | Radhanagari WLS, Maharashtra | This study |
| 24 | CCMB2382 | *Scolopendra* | *morsitans 1c* | - | + | + | Amrabad Tiger Reserve, Telangana | This study |
| 25 | CCMB3125 | *Scolopendra* | *hardwickei* | - | + | - | Jawadhu hills starting of the Ghat section, Tamil Nadu | This study |
| 26 | CCMB3772 | *Scolopendra* | *morsitans 1b* | + | + | + | Thirukkurungudi, Tamil Nadu | This study |
| 27 | CCMB3868 | *Scolopendra* | *morsitans 1b* | + | + | + | KMTR, Tamil Nadu | This study |
| 28 | CCMB4114 | *Scolopendra* | *morsitans 1b* | + | + | + | Injikuzhi, Tamil Nadu | This study |
| 29 | CCMB4216 | *Scolopendra* | *morsitans 1b* | + | + | + | Kumanoor, Kerala | This study |
| 30 | CCMB4407 | *Scolopendra* | *morsitans 1b* | + | + | + | Peechi, Kerala | This study |
| 31 | CCMB4441 | *Scolopendra* | *morsitans 1b* | + | + | + | Vazhani, Kerala | This study |
| 32 | CCMB4454 | *Scolopendra* | *morsitans 1b* | + | + | + | Vazhani, Kerala | This study |
| 33 | CES07101 | *Scolopendra* | *morsitans 1b* | - | MK273303 | - | Ramnagar, Banaglore, Karnataka, India | Joshi & Edgecombe, 2018 |
| 34 | CES07103 | *Scolopendra* | *morsitans 1b* | - | MK273304 | MK273428 | Ramnagar, Banaglore, Karnataka, India | Joshi & Edgecombe, 2018 |
| 35 | CES07105 | *Scolopendra* | *morsitans 1b* | - | MK273305 | MK273429 | Ramnagar, Banaglore, Karnataka, India | Joshi & Edgecombe, 2018 |
| 36 | CES07106 | *Scolopendra* | *morsitans 1b* | JN004006 | JN003895 | JN003951 | Ramnagar, Banaglore, Karnataka, India | Joshi and Karanth 2011 |
| 37 | CES07107 | *Scolopendra* | *morsitans 1b* | JN004007 | JN003896 | JN003952 | Ramnagar, Banaglore, Karnataka, India | Joshi and Karanth 2011 |
| 38 | CES07111 | *Scolopendra* | *morsitans 1b* | MK273201 | MK273306 | MK273430 | Ramnagar, Banaglore, Karnataka, India | Joshi & Edgecombe, 2018 |
| 39 | CES07116 | *Scolopendra* | *morsitans 1c* | - | MK273307 | - | Narsimha Devera Betta, Chikkaballapur, Karnataka, India | Joshi & Edgecombe, 2018 |
| 40 | CES07118 | *Scolopendra* | *morsitans 1c* | - | MK273308 | - | Narsimha Devera Betta, Chikkaballapur, Karnataka, India | Joshi & Edgecombe, 2018 |
| 41 | CES07120 | *Scolopendra* | *morsitans 1c* | MK273202 | MK273309 | MK273431 | Narsimha Devera Betta, Chikkaballapur, Karnataka, India | Joshi & Edgecombe, 2018 |
| 42 | CES07152 | *Scolopendra* | *morsitans 1c* | MK273205 | MK273313 | MK273435 | Pawgad, Karnataka, India | Joshi & Edgecombe, 2018 |
| 43 | CES07155 | *Scolopendra* | *morsitans 1b* | MK273206 | MK273314 | - | Chatancode, Vidura, Kerala, India | Joshi & Edgecombe, 2018 |
| 44 | CES07203 | *Scolopendra* | *morsitans 1a* | JN004008 | JN003897 | JN003953 | Devarayanadurga, Tumkur district, Karnataka, India | Joshi and Karanth 2011 |

| | | | | | | | | |
|---|---|---|---|---|---|---|---|---|
| 45 | CES07204 | *Scolopendra* | *morsitans 1a* | JN004009 | JN003898 | JN003954 | Devarayanadurga, Tumkur district, Karnataka, India | Joshi and Karanth 2011 |
| 46 | CES07206 | *Scolopendra* | *morsitans 1b* | - | MK273319 | MK273439 | Ramnagar, Banaglore, Karnataka, India | Joshi & Edgecombe, 2018 |
| 47 | CES07211 | *Scolopendra* | *morsitans 1c* | MK273212 | MK273320 | MK273440 | Bopdev ghat, Pune District, Maharashtra, India | Joshi & Edgecombe, 2018 |
| 48 | CES07212 | *Scolopendra* | *morsitans 1c* | JN004010 | JN003899 | JN003955 | Bopdev ghat, Pune District, Maharashtra, India | Joshi and Karanth 2011 |
| 49 | CES07213 | *Scolopendra* | *morsitans 1c* | MK273213 | MK273321 | - | Bopdev ghat, Pune District, Maharashtra, India | Joshi & Edgecombe, 2018 |
| 50 | CES07229 | *Scolopendra* | *morsitans 1b* | - | MK273324 | - | Anashi-Dandeli Tiger Reserve, Karwar District, Karnataka, India | Joshi & Edgecombe, 2018 |
| 51 | CES07235 | *Scolopendra* | *morsitans 1c* | JN004011 | JN003900 | JN003956 | Kumta, Uttara Kannada District, Karnataka, India | Joshi and Karanth 2011 |
| 52 | CES07241 | *Scolopendra* | *morsitans 1c* | - | MK273328 | MK273446 | Dughsagar Water Falls, Goa-Karnataka border, India | Joshi & Edgecombe, 2018 |
| 53 | CES07250 | *Scolopendra* | *morsitans 1b* | - | MK273330 | MK273447 | Mudumalai Tiger Reserve, Nilgiri District, Tamil Nadu, India | Joshi & Edgecombe, 2018 |
| 54 | CES07251 | *Scolopendra* | *morsitans 1c* | - | MK273331 | MK273448 | Kumta, Uttara Kannada District, Karnataka, India | Joshi & Edgecombe, 2018 |
| 55 | CES07252 | *Scolopendra* | *morsitans 1c* | JN004012 | JN003901 | JN003957 | Kumta, Uttara Kannada District, Karnataka, India | Joshi and Karanth 2011 |
| 56 | CES07259 | *Scolopendra* | *morsitans 1c* | MK273224 | MK273334 | MK273450 | Ranebennur, Haveri district, Karnataka, India | Joshi & Edgecombe, 2018 |
| 57 | CES07266 | *Scolopendra* | *morsitans 1c* | MK273225 | MK273335 | MK273451 | Ranebennur, Haveri district, Karnataka, India | Joshi & Edgecombe, 2018 |
| 58 | CES07267 | *Scolopendra* | *morsitans 1c* | MK273226 | MK273336 | MK273452 | Ranebennur, Haveri district, Karnataka, India | Joshi & Edgecombe, 2018 |
| 59 | CES07268 | *Scolopendra* | *morsitans 1c* | MK273227 | MK273337 | MK273453 | Ranebennur, Haveri district, Karnataka, India | Joshi & Edgecombe, 2018 |
| 60 | CES07269 | *Scolopendra* | *morsitans 1c* | MK273228 | MK273338 | MK273454 | Ranebennur, Haveri district, Karnataka, India | Joshi & Edgecombe, 2018 |
| 61 | CES07285 | *Scolopendra* | *hardwickei* | MK273231 | MK273341 | - | Bellari, Karnataka, India | Joshi & Edgecombe, 2018 |
| 62 | CES07287 | *Scolopendra* | *morsitans 1b* | MK273232 | MK273342 | MK273457 | Pooyyamkutty, Ernakulam district, Kerala, India | Joshi & Edgecombe, 2018 |
| 63 | CES08959 | *Scolopendra* | *hardwickei* | - | MK273356 | MK273466 | Amboli, Sindhudurg Dsitrict, Maharashtra, India | Joshi & Edgecombe, 2018 |
| 64 | CES091041 | *Scolopendra* | *morsitans 1c* | MK273256 | MK273372 | MK273480 | Agastya campus, Andhra Pradesh, India | Joshi & Edgecombe, 2018 |
| 65 | CES091392 | *Scolopendra* | *morsitans 2a* | MK273284 | - | MK273504 | Devmali, Orissa, India | Joshi & Edgecombe, 2018 |
| 66 | CES091411 | *Scolopendra* | *morsitans 2b* | MK273292 | MK273416 | MK273511 | Zara, Gujrath, India | Joshi & Edgecombe, 2018 |
| 67 | CES091412 | *Scolopendra* | *morsitans 2b* | MK273293 | - | MK273512 | Rampar, Gujrath, India | Joshi & Edgecombe, 2018 |
| 68 | CES091424 | *Scolopendra* | *morsitans 2b* | MK273294 | MK273417 | MK273513 | Champaner, Gujrath, India | Joshi & Edgecombe, 2018 |
| 69 | CES091459 | *Scolopendra* | *morsitans 2a* | MK273300 | MK273424 | MK273517 | Gudem, Andhra Pradesh, India | Joshi & Edgecombe, 2018 |
| 70 | CES091464 | *Scolopendra* | *morsitans 2c* | MK273302 | MK273426 | MK273519 | Tattapani, Himachal Pradesh, India | Joshi & Edgecombe, 2018 |
| 71 | CES091465 | *Scolopendra* | *morsitans 2c* | - | MK273427 | MK273520 | Sujanpur, Himachal Pradesh, India | Joshi & Edgecombe, 2018 |
| 72 | MCZ DNA103969 | *Scolopendra* | *afer* | KF676520 | KF676478 | KF676378 | Jakobsen's Beach, Tanzania | Vahtera et al. 2013 |
| 73 | MCZ DNA104817 | *Scolopendra* | *alternans* | KF676521 | KF676479 | - | Monroe County, Florida, USA | Vahtera et al. 2013 |
| 74 | CUMZ 00312 | *Scolopendra* | *calcarata* | KR705650 | KR705588 | - | Kanchanaburi, Thailand | Siriwut et al. 2015b |
| 75 | CUMZ 00417 | *Scolopendra* | *calcarata* | Can't use this s | KU512632 | - | Wat Mae Long, Mae Chaem, Chiang Mai, Thailand | Siriwut et al., 2016 |
| 76 | CUMZ 00418 | *Scolopendra* | *calcarata* | KU512630 | KU512633 | - | Lan Sang Waterfall, Mueang, Tak, Thailand | Siriwut et al., 2016 |
| 77 | MCZ DNA102462 | *Scolopendra* | *canidens* | KF676522 | KF676480 | KF676379 | Uzun district, Uzbekistan | Vahtera et al. 2013 |
| 78 | - | *Scolopendra* | *canidens* | JN688441 | JN688394 | - | Greece, Serifos | Oeyen et al., 2014 |
| 79 | CUMZ 00316 | *Scolopendra* | *cataracta* | KR705672 | KR705610 | - | Pakse, Champasak, Laos | Siriwut et al., 2015b |
| 80 | CUMZ 00317 | *Scolopendra* | *cataracta* | KR705633 | KR705571 | - | Tat Pha Yueang, Mueang Sing, Luang Namtha, Laos | Siriwut et al. 2015b |
| 81 | NHMUK010305528 | *Scolopendra* | *cataracta* | KU512631 | KU512634 | - | Kao Sok National Park, Surat Thani, Thailand | Siriwut et al., 2016 |
| 82 | MCZ DNA | *Scolopendra* | *cingulata* | HM453310 | HM453220 | AF000782 | Catalunya, Spain | Vahtera et al. 2013 |
| 83 | ZFMK-Sco-1 | *Scolopendra* | *cingulata* | KJ812067 | KJ812046 | - | Austria, Brugenland, Leitha Mts. | Oeyen et al., 2014 |
| 84 | Myr 01559 | *Scolopendra* | *cingulata* | KJ812077 | KJ812057 | - | Hungary, Ve´rtes Mts., Csa´kbere´ny, Bucka | Oeyen et al., 2014 |
| 85 | ZFMK-Sco-14/ZFMK-MYR-00585 | *Scolopendra* | *cingulata* | KJ812080 | KJ812060 | - | Greece, Kavala | Oeyen et al., 2014 |
| 86 | ZFMK-Sco-13 | *Scolopendra* | *cingulata* | KJ812082 | KJ812061 | - | Greece, Nestos Delta, Port Lagos | Oeyen et al., 2014 |
| 87 | ZFMK-Sco-11 | *Scolopendra* | *cingulata* | KJ812086 | KJ812065 | - | Romania, Anina | Oeyen et al., 2014 |
| 88 | ZFMK-Sco-12 | *Scolopendra* | *cingulata* | KJ812084 | KJ812063 | - | Turkey, Troy | Oeyen et al., 2014 |
| 89 | - | *Scolopendra* | *cretica* | JN688440 | JN688393 | - | Greece, Crete | Oeyen et al., 2014 |
| 90 | CUMZ 00272 | *Scolopendra* | *dawydoffi* | KR705618 | KR705680 | - | Saphan Hin Waterfall, Trad | Siriwut et al. 2015b |
| 91 | CUMZ 00290 | *Scolopendra* | *dawydoffi* | KR705654 | KR705592 | - | Sakearat, Nakhon Ratchasima | Siriwut et al. 2015b |
| 92 | CUMZ 00294.1 | *Scolopendra* | *dawydoffi* | KR705635 | KR705573 | - | Wat Thang Biang, Pak Chong, Nakhon Ratchasima | Siriwut et al. 2015b |
| 93 | CUMZ 00294.1 | *Scolopendra* | *dawydoffi* | KR705634 | KR705572 | - | Wat Thang Biang, Pak Chong, Nakhon Ratchasima | Siriwut et al. 2015b |
| 94 | CUMZ 00243 | *Scolopendra* | *dehaani* | KR705632 | KR705570 | - | Hub Pa-Tat, Lansak, Uthaithani | Siriwut et al. 2015b |

| | | | | | | | | |
|---|---|---|---|---|---|---|---|---|
| 95 | CUMZ 00247 | *Scolopendra* | *dehaani* | KR705655 | KR705593 | - | Ban Dongsavanh, Phang Khon, Sakon Nakhon | Siriwut et al. 2015b |
| 96 | CUMZ 00248 | *Scolopendra* | *dehaani* | KR705652 | KR705590 | - | Kaeng Lamduan, Namyeun, Ubon Ratchathani | Siriwut et al. 2015b |
| 97 | CUMZ 00251 | *Scolopendra* | *dehaani* | KR705640 | KR705578 | - | Sairung Waterfall, Takua Pa, Phangnga | Siriwut et al. 2015b |
| 98 | CUMZ 00252 | *Scolopendra* | *dehaani* | KR705683 | KR705621 | - | Sichang Island, Chonburi | Siriwut et al. 2015b |
| 99 | CUMZ 00253 | *Scolopendra* | *dehaani* | KR705630 | KR705568 | - | Tham Khao Bin, Ratchaburi | Siriwut et al. 2015b |
| 100 | CUMZ 00256 | *Scolopendra* | *dehaani* | KR705688 | KR705626 | - | Sapanthai, Bangban, Ayutthaya | Siriwut et al. 2015b |
| 101 | CUMZ 00262 | *Scolopendra* | *dehaani* | KR705637 | KR705575 | - | JPR Stone Park, Kraburi, Ranong | Siriwut et al. 2015b |
| 102 | | *Scolopendra* | *hardwickei* | KF676523 | KF676482 | KF676381 | | Vahtera et al. 2013 |
| 103 | CUMZ 00319 | *Scolopendra* | *japonica* | KR705617 | KR705679 | - | Shinshu University, Matsumoto, Japan | Siriwut et al. 2015b |
| 104 | CUMZ 00298-1 | *Scolopendra* | *japonica* | KR705671 | KR705609 | - | Plain of Jar, Xieang Khouang, Laos | Siriwut et al. 2015b |
| 105 | CUMZ 00298-2 | *Scolopendra* | *japonica* | KR705670 | KR705608 | - | Plain of Jar, Xieang Khouang, Laos | Siriwut et al. 2015b |
| 106 | CUMZ 00297-1 | *Scolopendra* | *japonica* | KR705675 | KR705613 | - | Phu Fah Mountain, Phongsali, Laos | Siriwut et al. 2015b |
| 107 | CUMZ 00297-2 | *Scolopendra* | *japonica* | KR705674 | KR705612 | - | Phu Fah Mountain, Phongsali, Laos | Siriwut et al. 2015b |
| 108 | MCZ DNA103954 | *Scolopendra* | *laeta* | KF676541 | KF676483 | KF676382 | Cowra Shire, New South Wales, Australia | Vahtera et al. 2013 |
| 109 | MCZ DNA120857 | *Scolopendra* | *leki* | KF676524 | KF676484 | - | Southern Carnarvon Range, Western Australia, Australia | Vahtera et al. 2013 |
| 110 | MCZ DNA104707 | *Scolopendra* | *morsitans* | HQ402553 | HQ402501 | - | E. of Sntiou Malem, Senegal | Vahtera et al. 2012 |
| 111 | CUMZ 00339 | *Scolopendra* | *morsitans* | KR705662 | KR705600 | - | Ban Dan Chang, Ta Kantho, Khonkaen | Siriwut et al. 2015b |
| 112 | CUMZ 00340 | *Scolopendra* | *morsitans* | KR705661 | KR705599 | - | Lainan, Weing Sa, Nan | Siriwut et al. 2015b |
| 113 | CUMZ 00341 | *Scolopendra* | *morsitans* | KR705660 | KR705598 | - | Juang Island, Sattahip, Chonburi | Siriwut et al. 2015b |
| 114 | CUMZ 00342 | *Scolopendra* | *morsitans* | KR705666 | KR705604 | - | Ban Khok Pho, Prasat, Surin | Siriwut et al. 2015b |
| 115 | CUMZ 00343 | *Scolopendra* | *morsitans* | KR705665 | KR705603 | - | Hui Hong Khrai, Chiangmai | Siriwut et al. 2015b |
| 116 | CUMZ 00344 | *Scolopendra* | *morsitans* | KR705664 | KR705602 | - | Tha Kra Bak Reservoir, Srakaeo | Siriwut et al. 2015b |
| 117 | CUMZ 00345 | *Scolopendra* | *morsitans* | KR705663 | KR705601 | - | Wat Phanombak, Srisophon, Cambodia | Siriwut et al. 2015b |
| 118 | MCZ DNA107083 | *Scolopendra* | *multidens* | KF676540 | KF676485 | - | Qiang Binh, Vietnam | Vahtera et al. 2013 |
| 119 | MCZ DNA106873 | *Scolopendra* | *oraniensis* | - | KF676486 | - | Nuoro, Sardinia, Italy | Vahtera et al. 2013 |
| 120 | Myr 00568 | *Scolopendra* | *oraniensis* | KJ812088 | KJ812066 | - | Marokko, Prov. Nador, Atlas Mts. | Oeyen et al., 2014 |
| 121 | CUMZ 00303 | *Scolopendra* | *pinguis* | KR705645 | KR705584 | - | Wat Tham Lijia, Sangkhlaburi, Kanchanaburi | Siriwut et al. 2015b |
| 122 | CUMZ 00304 | *Scolopendra* | *pinguis* | KR705642 | KR705580 | - | Wiang Thong Hotspring, Mueang Ieam, Huaphan, Laos | Siriwut et al. 2015b |
| 123 | CUMZ 00305 | *Scolopendra* | *pinguis* | KR705647 | KR705585 | - | Phusang Waterfall, Phayao | Siriwut et al. 2015b |
| 124 | CUMZ 00306 | *Scolopendra* | *pinguis* | KR705643 | KR705581 | - | Ban Na-Ton, Muang Khun, Xieng Khouang, Laos | Siriwut et al. 2015b |
| 125 | CUMZ 00307 | *Scolopendra* | *pinguis* | KR705644 | KR705582 | - | Hui Nam-Un, Wiangkhum, Nan | Siriwut et al. 2015b |
| 126 | CUMZ 00309 | *Scolopendra* | *pinguis* | KR705646 | KR705584 | - | Khao Rao Cave, Bokaeo, Laos | Siriwut et al. 2015b |
| 127 | CUMZ 00313 | *Scolopendra* | *pinguis* | KR705649 | KR705587 | - | Ban Pang Pan, Maetaeng, Chiangmai | Siriwut et al. 2015b |
| 128 | CUMZ 00314 | *Scolopendra* | *pinguis* | KR705648 | KR705586 | - | Phamone Cave, Pangmapha, Maehongson | Siriwut et al. 2015b |
| 129 | MCZ DNA10158 | *Scolopendra* | *polymorpha* | KF676525 | KF676487 | KF676384 | Las Cruces, New Mexico, USA | Vahtera et al. 2013 |
| 130 | CUMZ 00315 | *Scolopendra* | *subspinipes* | KR705636 | KR705574 | - | Singapore | Siriwut et al. 2015b |
| 131 | MCZ DNA106501/IZ-130685 | *Scolopendra* | *subspinipes subspini* | KF676528 | KF676488 | KF676387 | Weam, Western Province, Papua New Guinea | Vahtera et al. 2013 |
| 132 | MCZ DNA104694 | *Scolopendra* | *subspinipes subspini* | HQ402554 | HQ402502 | HQ402538 | Martinique | Vahtera et al. 2012 |
| 133 | MCAZ DNA100675 | *Scolopendra* | *viridis* | DQ201431 | DQ201425 | DQ222134 | Gila Cliff Dwellings, New Mexico, USA | Vahtera et al. 2012 |
| 134 | TS-20180301-01 | *Scolopendra* | *alcyona* | LC504083 | LC504073 | - | Yona, Okinawa-jima Island, Okinawa | Tsukamoto et al., 2021 |
| 135 | TS-20180912-01 | *Scolopendra* | *alcyona* | LC504090 | LC504081 | - | Yona, Okinawa-jima Island, Okinawa | Tsukamoto et al., 2021 |
| 136 | CES07225 | *Asanada* | *sp.* | JN004013 | JN003902 | JN003958 | Anashi-Dandeli Tiger Reserve, Karwar district, Karnataka, India | Joshi and Karanth 2011 |
| 137 | CES08951 | *Asanada* | *sp.* | JN004014 | JN003903 | - | Amboli, Sindhudurg district, Maharashtra, India | Joshi and Karanth 2011 |
| 138 | CES08954 | *Asanada* | *sp.* | MK273244 | MK273355 | MK273465 | Amboli, Sindhudurg district, Maharashtra, India | Joshi and Edgecombe 2017 |
| 139 | CES091369 | *Asanada* | *sp.* | MK273280 | MK273403 | MK273500 | Kolli hills, Namakkal district , Tamil Nadu, India | Joshi and Edgecombe 2018 |
| 140 | | *Asanada* | *breviconis* | HQ402540 | HQ402489 | - | Nongbua, Nakhon Sawan, Thailand | Vahtera et al. 2012 |
| 141 | | *Asanada* | *socotrana* | - | HQ402490 | HQ402524 | Senegal | Vahtera et al. 2012 |
| 142 | CES07124 | *Cormocephalus* | *sp3* | MK273203 | MK273310 | MK273432 | Silent Valley National Park, Kerala, India | Joshi & Edgecombe, 2018 |
| 143 | CES07129 | *Cormocephalus* | *sp5* | JN003997 | JN003886 | JN003942 | Silent Valley National Park, Palakkad district, Kerala, India | Joshi and Karanth 2011 |
| 144 | CES07136 | *Cormocephalus* | *sp5* | JN003998 | JN003887 | JN003943 | Silent Valley National Park, Palakkad district, Kerala, India | Joshi and Karanth 2011 |

| | | | | | | | | |
|---|---|---|---|---|---|---|---|---|
| 145 | CES07139 | *Cormocephalus* | *sp5* | JN003999 | JN003888 | JN003944 | Silent Valley National Park, Palakkad district, Kerala, India | Joshi and Karanth 2011 |
| 146 | CES07147 | *Cormocephalus* | *sp5* | - | MK273312 | MK273434 | Silent Valley National Park, Kerala, India | Joshi & Edgecombe, 2018 |
| 147 | CES07163 | *Cormocephalus* | *sp5* | MK273209 | MK273317 | - | Neyyar Wildlife Sanctuary, Thiruvananthapuram District, Kerala, | Joshi & Edgecombe, 2018 |
| 148 | CES07199 | *Cormocephalus* | *sp3* | MK273210 | MK273318 | - | Ponmudi Reserve Forest, Thiruvananthapuram District, Kerala, I | Joshi & Edgecombe, 2018 |
| 149 | CES07200 | *Cormocephalus* | *sp2* | JN004000 | JN003889 | JN003945 | Ponmudi Reserve Forest, Thiruvananthapuram District, Kerala, I | Joshi and Karanth 2011 |
| 150 | CES07205 | *Cormocephalus* | *sp3* | JN004001 | JN003890 | JN003946 | Ramanagar, Ramanagara district, Karnataka, India | Joshi and Karanth 2011 |
| 151 | CES07220 | *Cormocephalus* | *sp3* | JN004002 | JN003891 | JN003947 | Anashi-Dandeli Tiger Reserve, Karwar district, Karnataka, India | Joshi and Karanth 2011 |
| 152 | CES07243 | *Cormocephalus* | *sp3* | MK273220 | MK273329 | - | Dughsagar Water Falls, Goa-Karnataka border, India | Joshi & Edgecombe, 2018 |
| 153 | CES07254 | *Cormocephalus* | *sp3* | MK273222 | MK273332 | MK273449 | Ranebennur, Haveri district, Karnataka, India | Joshi & Edgecombe, 2018 |
| 154 | CES07256 | *Cormocephalus* | *sp3* | JN004003 | JN003892 | JN003948 | Ranebennur, Haveri district, Karnataka, India | Joshi and Karanth 2011 |
| 155 | CES07257 | *Cormocephalus* | *sp3* | MK273223 | MK273333 | - | Ranebennur, Haveri district, Karnataka, India | Joshi & Edgecombe, 2018 |
| 156 | CES07258 | *Cormocephalus* | *sp3* | JN004004 | JN003893 | JN003949 | Devarayanadurga, Tumkur district, Karnataka, India | Joshi & Karanth, 2011 |
| 157 | CES07265 | *Cormocephalus* | *sp3* | JN004005 | JN003894 | - | Devarayanadurga, Tumkur district, Karnataka, India | Joshi and Karanth 2011 |
| 158 | CES07296 | *Cormocephalus* | *sp3* | MK273233 | - | - | Dimbam, Periyar, Tamil Nadu, India | Joshi & Edgecombe, 2018 |
| 159 | CES08923 | *Cormocephalus* | *sp6* | MK273241 | MK273351 | - | Tadoli, Kudremukh National Park, India | Joshi & Edgecombe, 2018 |
| 160 | CES091022 | *Cormocephalus* | *sp4* | - | MK273365 | - | Bisale Ghat Reserve Forest, Hassan district, Karnataka, India | Joshi & Edgecombe, 2018 |
| 161 | CES091027 | *Cormocephalus* | *sp4* | - | MK273366 | - | Kodagu district, Karnataka, India | Joshi & Edgecombe, 2018 |
| 162 | CES091051 | *Cormocephalus* | *sp2* | MK273257 | MK273373 | - | Parambikulam Tiger Reserve, Kerala, India | Joshi & Edgecombe, 2018 |
| 163 | CES091053 | *Cormocephalus* | *sp5* | MK273258 | MK273374 | - | Sholayar Reserve Forest, Kerala, India | Joshi & Edgecombe, 2018 |
| 164 | CES091054 | *Cormocephalus* | *sp5* | MK273259 | MK273375 | MK273481 | Sholayar Reserve Forest, Ernakulam district, Kerala, India | Joshi & Edgecombe, 2018 |
| 165 | CES091055 | *Cormocephalus* | *sp2* | MK273260 | MK273376 | - | Sholayar Reserve Forest, Ernakulam district, Kerala, India | Joshi & Edgecombe, 2018 |
| 166 | CES091059 | *Cormocephalus* | *sp2* | MK273261 | MK273377 | - | Athirapally, Thrissur district, Kerala, India | Joshi & Edgecombe, 2018 |
| 167 | CES091060 | *Cormocephalus* | *sp2* | MK273262 | MK273378 | MK273482 | Athirapally, Thrissur district, Kerala, India | Joshi & Edgecombe, 2018 |
| 168 | CES091074 | *Cormocephalus* | *sp5* | MK273263 | MK273379 | MK273483 | Parakkadavu, Shendurney Wildlife Sanctuary, Kollam district, Ke | Joshi & Edgecombe, 2018 |
| 169 | CES091075 | *Cormocephalus* | *sp2* | - | MK273380 | - | Parakkadavu, Shendurney Wildlife Sanctuary, Kerala, India | Joshi & Edgecombe, 2018 |
| 170 | CES091084 | *Cormocephalus* | *sp5* | MK273267 | MK273385 | MK273487 | Rosemla forest, Shendurney Wildlife Sanctuary, Kerala, India | Joshi & Edgecombe, 2018 |
| 171 | CES091093 | *Cormocephalus* | *sp5* | - | MK273387 | - | Pandimatte, Shendurney Wildlife Sanctuary, Kerala, India | Joshi & Edgecombe, 2018 |
| 172 | CES091097 | *Cormocephalus* | *sp2* | MK273269 | MK273389 | MK273490 | Pandimatte, Shendurney Wildlife Sanctuary, Kollam district, Kera | Joshi & Edgecombe, 2018 |
| 173 | CES091312 | *Cormocephalus* | *sp2* | MK273271 | MK273393 | - | Vellithode, Periyar Tiger Reserve, Kerala, India | Joshi & Edgecombe, 2018 |
| 174 | CES091314 | *Cormocephalus* | *sp2* | MK273273 | MK273395 | - | Vellithode, Periyar Tiger Reserve, Kerala, India | Joshi & Edgecombe, 2018 |
| 175 | CES091317 | *Cormocephalus* | *sp5* | MK273274 | MK273396 | MK273495 | Vellithode, Periyar Tiger Reserve, Kerala, India | Joshi & Edgecombe, 2018 |
| 176 | CES091332 | *Cormocephalus* | *sp5* | - | MK273398 | - | Siruvani Reserve Forest, Kerala, India | Joshi & Edgecombe, 2018 |
| 177 | CES091340 | *Cormocephalus* | *sp5* | MK273277 | MK273399 | MK273497 | Pasikada, Aachankovil Reserve Forest, Kerala, India | Joshi & Edgecombe, 2018 |
| 178 | CES091374 | *Cormocephalus* | *sp6* | - | MK273404 | - | Kolli hills, Namakkal district , Tamil Nadu, India | Joshi & Edgecombe, 2018 |
| 179 | CES091383 | *Cormocephalus* | *sp* | - | MK273406 | - | Javadi hills, Tiruvannamalai, Tamil Nadu, India | Joshi & Edgecombe, 2018 |
| 180 | MCZ DNA100679 | *Cormocephalus* | *anceps* | KF676535 | KF676494 | KF676399 | Western Cape, South Africa | Vahtera et al. 2013 |
| 181 | MCZ DNA103951 | *Cormocephalus* | *aurantiipes* | HQ402543 | HQ402492 | KF676388 | Western Australia, Australia | Vahtera et al. 2012 |
| 182 | MCZ DNA103964 | *Cormocephalus* | *bonaerius* | KF676530 | KF676489 | KF676389 | Parque de Cahuita, Costa Rica | Vahtera et al. 2013 |
| 183 | MCZ DNA106754 | *Cormocephalus* | *elegans* | KF676536 | KF676495 | KF676400 | KwaZulu-Natal, South Africa | Vahtera et al. 2013 |
| 184 | MCZ DNA104642 | *Cormocephalus* | *gervaisianus* | - | KF676490 | KF676390 | Tunis District, Tunisia | Vahtera et al. 2013 |
| 185 | MCZ DNA100377 | *Cormocephalus* | *hartmeyeri* | KF676531 | KF676491 | KF676391 | Pemberton, Western Australia, Australia | Vahtera et al. 2013 |
| 186 | MCZ DNA100274 | *Cormocephalus* | *monteithi* | - | AF370861 | AF173280 | Kenilworth State Forest, Queensland, Australia | Vahtera et al. 2012 |
| 187 | MCZ DNA106751 | *Cormocephalus* | *nitidus* | KF676533 | KF676492 | KF676397 | KwaZulu-Natal, South Africa | Vahtera et al. 2013 |
| 188 | MCZ DNA106752 | *Cormocephalus* | *pseudopunctatus* | KF676534 | KF676493 | KF676398 | KwaZulu-Natal, South Africa | Vahtera et al. 2013 |
| 189 | MCZ DNA106510 | *Cormocephalus* | *westwoodi* | KF676537 | KF676496 | KF676401 | Armidale, New South Wales, Australia | Vahtera et al. 2013 |

**Supplementary information 3**

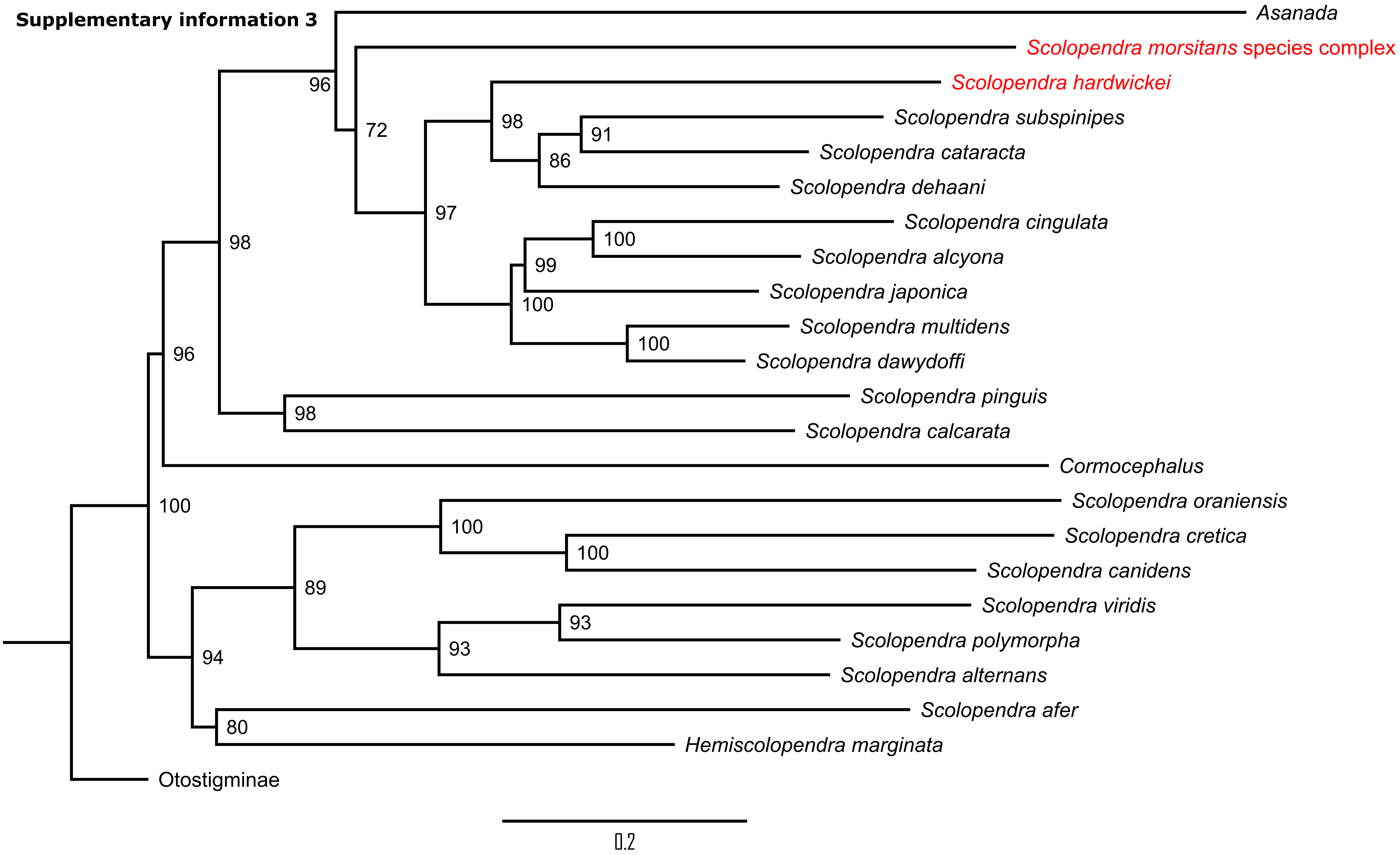

Note: Bootstrap values are indicated at the nodes in the above ML tree

### Supplementary Tables

| | | | | | | | |
|---|---|---|---|---|---|---|---|
| MS Title: | Divergent venoms among two closely related co-distributed centipede species, *Scolopendra morsitans* and *S. hardwickei* in tropical Asia | | | | | | |
| Authors: | Aditi | Pragyadeep Roy | Richard Parikh | Aniruddha Marathe | Karunakar Majhi | Ronald Jenner | Jahnavi Joshi* |
| Affiliations | CSIR-Centre for Cellular and Molecular Biology, Hyderabad, India | CSIR-Centre for Cellular and Molecular Biology, Hyderabad, India | CSIR-Centre for Cellular and Molecular Biology, Hyderabad, India | CSIR-Centre for Cellular and Molecular Biology, Hyderabad, India | CSIR-Centre for Cellular and Molecular Biology, Hyderabad, India | Natural History Museum, London, United Kingdom | CSIR-Centre for Cellular and Molecular Biology, Hyderabad, India. |
| | Academy of Scientific and Innovative Research, Ghaziabad, Uttar Pradesh, India | Academy of Scientific and Innovative Research, Ghaziabad, Uttar Pradesh, India | Indian Institute of Science, Education and Research, Pune, India | | | | Academy of Scientific and Innovative Research, Ghaziabad, Uttar Pradesh, India. |
| *Email ID for correspondence | | | | | | | |
| Table legends | | | | | | | |
| Table S1 (Page 9) | **Transcriptome:** Venom gland Transcriptomes assembly information of both *S. morsitans* and *S. hardwickei* | | | | | | |
| Tables S2 (Pages 10 - 12) | **General summary**: This series of sheets contains the sampling information, matrix generated, index calculations and the list of unique proteins identified in both the species | | | | | | |
| Tables S3 (Pages 13 - 14) | **PCA data**: This series of sheets contains information about the PCN matrix used to build the PCA plot and the result loadings | | | | | | |

**N.B. The contents might not be visible at the first look, but please zoom them and they would be legible.**

| Species | *S. hardwickei* | *S. morsitans* |
|---|---|---|
| Vocher ID | CCMB1692 | CCMB845 |
| Location | Laknavaram, Telangana | Ranebennur WLS, Karnataka |
| Latitude | 18.16381 | 14.666814 |
| Longitude | 80.06749 | 75.662201 |
| Elevation | 201 | 589 |
| Total raw reads | 36,447,924 | 50,240,329 |
| Reads with adapters | 19,100,903 | 44,468,575 |
| Adapter sequence | AGATCGGAAGAGC | AGATCGGAAGAGC |
| Total bp processed for trimming | 5,503,636,524 | 7,586,289,679 |
| Quality trimmed bp | 106,148,480 | 213,101,356 |
| Filtered bp | 5,199,671,023 | 4,616,941,559 |
| Reads after trimming | 35,680,848 | 49614068 |
| GC percentage | 42 | 42 |
| Sequence length | 30-151 | 20-151 |
| Total transcripts | 111,540 | 223,104 |
| N50 | 2191 | 619 |
| Unidentified transcripts (TPM >1) | 78675 | 154203 |
| Non-toxin transcripts (TPM>1) | 1444 | 1303 |
| Toxin transcripts (TPM >1) | 2323 | 2186 |

**SI Table 2.a: Sampling Information**

| Specimen | Species name | Location | Longitude | Latitude | Elevation |
|---|---|---|---|---|---|
| CCMB41 | *Scolopendra morsitans 1c* | Nehru Zoo, Telangana | 78.452145 | 17.350547 | 533 |
| CCMB752 | *Scolopendra morsitans 1a* | Panchgani, Maharashtra | 73.821493 | 17.929342 | 1,320 |
| CCMB803 | *Scolopendra morsitans 1c* | Ranebennur WLS, Karnataka | 75.664520 | 14.649282 | 617 |
| CCMB805 | *Scolopendra morsitans 1c* | Ranebennur WLS, Karnataka | 75.664520 | 14.649282 | 617 |
| CCMB818 | *Scolopendra morsitans 1c* | Kolhapur, Maharashtra | 74.080209 | 16.631231 | 568 |
| CCMB845 | *Scolopendra morsitans 1c* | Ranebennur WLS, Karnataka | 75.662201 | 14.666814 | 589 |
| CCMB1159 | *Scolopendra morsitans* | Mookambika WLS, Karnataka | 74.864420 | 13.878260 | 627 |
| CCMB1209 | *Scolopendra morsitans 1c* | Kadappa city, Andhra Pradesh | 78.855990 | 14.444500 | 159 |
| CCMB1245 | *Scolopendra hardwickei* | Lankamalla, Andhra Pradesh | 78.925990 | 14.501780 | 147 |
| CCMB1325 | *Scolopendra morsitans 1b* | Horsley Hills, Andhra Pradesh | 78.396890 | 13.648710 | 1,251 |
| CCMB1691 | *Scolopendra hardwickei* | Laknavaram, Telangana | 80.07301 | 18.16355 | 198 |
| CCMB1692 | *Scolopendra hardwickei* | Laknavaram, Telangana | 80.06749 | 18.16381 | 201 |
| CCMB1693 | *Scolopendra morsitans* | Laknavaram, Telangana | 80.06749 | 18.16381 | 201 |
| CCMB1695 | *Scolopendra morsitans* | Laknavaram, Telangana | 80.06749 | 18.16381 | 201 |
| CCMB1696 | *Scolopendra morsitans 1c* | Laknavaram, Telangana | 80.06749 | 18.16381 | 201 |
| CCMB1725 | *Scolopendra morsitans 1c* | Tadvai, Telangana | 80.31848 | 18.20765 | 200 |
| CCMB1766 | *Scolopendra morsitans 1c* | Sarvapur beat, Telanagana | 80.032620 | 18.117620 | 292 |
| CCMB1769 | *Scolopendra morsitans 1c* | Sarvapur beat, Telanagana | 80.030270 | 18.119400 | 224 |
| CCMB1776 | *Scolopendra hardwickei* | Sarvapur beat, Telanagana | 80.032620 | 18.117620 | 292 |
| CCMB1803 | *Scolopendra morsitans* | Gandikota, Andhra Pradesh | 78.746060 | 14.531070 | 146 |
| CCMB2184 | *Scolopendra morsitans* | Devihasol, Maharashtra | 73.424560 | 16.745440 | 127 |
| CCMB2194 | *Scolopendra hardwickei* | Radhanagari WLS, Maharashtra | 73.955370 | 16.348910 | 953 |
| CCMB2197 | *Scolopendra hardwickei* | Radhanagari WLS, Maharashtra | 73.955370 | 16.348910 | 953 |
| CCMB2198 | *Scolopendra morsitans* | Radhanagari WLS, Maharashtra | 73.945690 | 16.346030 | 1,004 |
| CCMB2382 | *Scolopendra morsitans 1c* | Amrabad Tiger Reserve, Telangana | 78.655160 | 16.185760 | 809 |
| CCMB3125 | *Scolopendra hardwickei* | Jawadhu hills, TamilNadu | 78.774305 | 12.376218 | 485 |
| CCMB3265 | *Scolopendra morsitans* | Phansad, Maharashtra | 72.944260 | 18.425710 | 299 |
| CCMB3279 | *Scolopendra hardwickei* | Phansad, Maharashtra | 72.934490 | 18.438570 | 299 |
| CCMB3280 | *Scolopendra hardwickei* | Phansad, Maharashtra | 72.941630 | 18.430640 | 288 |
| CCMB3281 | *Scolopendra hardwickei* | Phansad, Maharashtra | 72.934490 | 18.438570 | 299 |
| CCMB3338 | *Scolopendra hardwickei* | Lonavala, Maharashtra | 73.389270 | 18.710590 | 769 |
| CCMB3772 | *Scolopendra morsitans 1b* | Thirukkurungudi, TamilNadu | 77.545380 | 8.405420 | 195 |
| CCMB3868 | *Scolopendra morsitans 1b* | KMTR, TamilNadu | 77.282250 | 8.663210 | 283 |
| CCMB4114 | *Scolopendra morsitans 1b* | Injikuzhi, TamilNadu | 77.286280 | 8.659820 | 285 |
| CCMB4216 | *Scolopendra morsitans 1b* | Kumanoor, Kerala | 76.892930 | 9.209570 | 71 |
| CCMB4224 | *Scolopendra morsitans* | Kumanoor, Kerala | 76.892930 | 9.209570 | 71 |
| CCMB4407 | *Scolopendra morsitans 1b* | Peechi, Kerala | 76.37338 | 10.53165 | 81 |
| CCMB4441 | *Scolopendra morsitans 1b* | Vazhani, Kerala | 76.303530 | 10.630280 | 83 |
| CCMB4454 | *Scolopendra morsitans 1b* | Vazhani, Kerala | 76.302280 | 10.627840 | 145 |

**SI Table 2.b: The Protein Copy Number Matrix generated using the Protein Discoverer results**

| ID | species | Acid-phosphatase | Alpha-2-macroglobulin | ALPHA-MANNOSIDASE | Apolipoprotein D | ARYLSULFATASE_ | BPFTx | C-type_Lectin | CAA-like | Calycin-Lipocalin | CAP | Carbon-nitrogen_hydrolase | Carboxylic ester hydrolase | Cathepsin | Chitinase | Chondroitinase | COEsteraseB | CUB-domai_protein | Cubilin_ | Cyclase-like_protein | Cystatin | Cysteine-rich venom protein | DUF1397 | DUF3472 | Fibronectin_or_immunoglobulin-like | GDH | GGT | GH18 | Haem-binding_protein | Icarapin-like | Ige-ESP-like |
|---|---|---|---|---|---|---|---|---|---|---|---|---|---|---|---|---|---|---|---|---|---|---|---|---|---|---|---|---|---|---|---|
| CCMB1159 | *morsitans* | 0 | 2 | 1 | 0 | 0 | 18 | 1 | 0 | 0 | 11 | 0 | 0 | 1 | 1 | 1 | 1 | 13 | 1 | 0 | 1 | 0 | 0 | 12 | 0 | 3 | 1 | 0 | 0 | 1 | 0 |
| CCMB1209 | *morsitans* | 0 | 0 | 2 | 0 | 0 | 26 | 1 | 0 | 0 | 8 | 0 | 0 | 1 | 2 | 1 | 5 | 16 | 3 | 0 | 1 | 0 | 0 | 12 | 0 | 3 | 1 | 0 | 0 | 0 | 0 |
| CCMB1325 | *morsitans* | 0 | 1 | 2 | 0 | 0 | 31 | 2 | 0 | 0 | 11 | 0 | 0 | 0 | 1 | 1 | 1 | 16 | 3 | 0 | 1 | 0 | 0 | 8 | 0 | 3 | 0 | 0 | 0 | 0 | 0 |
| CCMB1693 | *morsitans* | 0 | 3 | 2 | 0 | 0 | 31 | 3 | 0 | 0 | 13 | 1 | 0 | 0 | 2 | 2 | 6 | 14 | 3 | 0 | 4 | 0 | 0 | 8 | 0 | 2 | 1 | 0 | 1 | 1 | 0 |
| CCMB1695 | *morsitans* | 0 | 0 | 0 | 0 | 0 | 27 | 0 | 0 | 0 | 12 | 0 | 0 | 0 | 1 | 1 | 4 | 12 | 2 | 0 | 2 | 0 | 0 | 4 | 0 | 2 | 1 | 0 | 0 | 0 | 0 |
| CCMB1696 | *morsitans* | 1 | 0 | 1 | 0 | 0 | 32 | 2 | 0 | 0 | 8 | 0 | 0 | 0 | 1 | 2 | 5 | 15 | 3 | 0 | 2 | 0 | 0 | 7 | 0 | 2 | 0 | 0 | 0 | 1 | 1 |
| CCMB1725 | *morsitans* | 0 | 12 | 0 | 1 | 0 | 21 | 1 | 0 | 2 | 8 | 7 | 0 | 0 | 3 | 1 | 2 | 11 | 3 | 0 | 2 | 0 | 1 | 2 | 0 | 1 | 0 | 4 | 1 | 1 | 0 |
| CCMB1766 | *morsitans* | 0 | 0 | 1 | 0 | 0 | 32 | 2 | 0 | 0 | 14 | 0 | 0 | 0 | 2 | 2 | 4 | 19 | 3 | 0 | 2 | 1 | 0 | 8 | 1 | 2 | 0 | 0 | 0 | 1 | 1 |
| CCMB1769 | *morsitans* | 0 | 1 | 2 | 0 | 0 | 33 | 2 | 0 | 0 | 11 | 0 | 0 | 0 | 2 | 2 | 5 | 16 | 3 | 0 | 2 | 0 | 0 | 10 | 0 | 1 | 0 | 0 | 0 | 1 | 0 |
| CCMB1803 | *morsitans* | 0 | 2 | 2 | 0 | 0 | 27 | 1 | 0 | 0 | 8 | 0 | 0 | 1 | 1 | 1 | 2 | 16 | 3 | 0 | 2 | 0 | 0 | 9 | 0 | 2 | 3 | 0 | 0 | 0 | 0 |
| CCMB2184 | *morsitans* | 0 | 1 | 1 | 0 | 0 | 26 | 2 | 0 | 0 | 8 | 0 | 0 | 0 | 2 | 1 | 2 | 17 | 4 | 0 | 2 | 0 | 0 | 11 | 0 | 1 | 0 | 0 | 0 | 1 | 0 |
| CCMB2198 | *morsitans* | 0 | 0 | 1 | 0 | 0 | 26 | 1 | 0 | 0 | 7 | 0 | 0 | 1 | 2 | 1 | 2 | 15 | 3 | 0 | 2 | 0 | 0 | 9 | 0 | 2 | 0 | 0 | 0 | 1 | 0 |
| CCMB2382 | *morsitans* | 0 | 6 | 0 | 0 | 0 | 20 | 2 | 0 | 1 | 11 | 2 | 0 | 0 | 1 | 1 | 3 | 11 | 6 | 0 | 3 | 1 | 0 | 3 | 0 | 1 | 0 | 0 | 1 | 0 | 0 |
| CCMB3265 | *morsitans* | 0 | 13 | 1 | 0 | 0 | 17 | 3 | 0 | 2 | 6 | 4 | 0 | 0 | 2 | 1 | 1 | 9 | 2 | 0 | 1 | 1 | 1 | 4 | 0 | 1 | 2 | 1 | 1 | 0 | 0 |
| CCMB3772 | *morsitans* | 0 | 13 | 1 | 0 | 1 | 19 | 2 | 0 | 2 | 10 | 0 | 0 | 1 | 2 | 1 | 4 | 12 | 3 | 1 | 2 | 1 | 0 | 9 | 0 | 2 | 0 | 0 | 2 | 0 | 0 |
| CCMB3868 | *morsitans* | 0 | 0 | 1 | 0 | 0 | 19 | 0 | 0 | 0 | 6 | 0 | 0 | 0 | 1 | 1 | 1 | 12 | 3 | 0 | 3 | 0 | 0 | 7 | 0 | 2 | 0 | 1 | 1 | 0 | 1 |
| CCMB41 | *morsitans* | 0 | 5 | 0 | 0 | 0 | 5 | 0 | 0 | 0 | 7 | 1 | 0 | 0 | 3 | 0 | 0 | 5 | 3 | 0 | 1 | 0 | 0 | 2 | 0 | 0 | 0 | 0 | 0 | 0 | 0 |
| CCMB4114 | *morsitans* | 0 | 0 | 0 | 0 | 0 | 20 | 0 | 2 | 0 | 5 | 0 | 0 | 0 | 1 | 1 | 2 | 14 | 1 | 0 | 0 | 0 | 0 | 7 | 1 | 2 | 0 | 0 | 0 | 0 | 2 |
| CCMB4216 | *morsitans* | 0 | 8 | 1 | 1 | 1 | 22 | 1 | 0 | 1 | 6 | 1 | 0 | 0 | 2 | 1 | 4 | 13 | 4 | 1 | 2 | 1 | 0 | 8 | 17 | 2 | 0 | 0 | 1 | 0 | 1 |
| CCMB4224 | *morsitans* | 0 | 3 | 1 | 0 | 1 | 25 | 0 | 0 | 1 | 6 | 0 | 0 | 1 | 1 | 1 | 2 | 13 | 3 | 0 | 3 | 2 | 0 | 7 | 0 | 2 | 0 | 0 | 1 | 0 | 1 |
| CCMB4407 | *morsitans* | 0 | 8 | 1 | 1 | 1 | 22 | 1 | 0 | 2 | 4 | 1 | 0 | 0 | 2 | 1 | 4 | 12 | 3 | 0 | 2 | 0 | 0 | 5 | 17 | 3 | 0 | 2 | 1 | 0 | 0 |
| CCMB4441 | *morsitans* | 0 | 0 | 1 | 0 | 0 | 23 | 0 | 0 | 0 | 3 | 0 | 0 | 0 | 1 | 1 | 3 | 13 | 4 | 0 | 1 | 0 | 0 | 4 | 0 | 3 | 0 | 0 | 0 | 0 | 0 |
| CCMB4454 | *morsitans* | 0 | 0 | 1 | 0 | 0 | 20 | 0 | 0 | 0 | 3 | 0 | 0 | 0 | 1 | 1 | 2 | 11 | 4 | 0 | 1 | 0 | 0 | 7 | 0 | 2 | 0 | 0 | 0 | 0 | 0 |
| CCMB752 | *morsitans* | 0 | 4 | 1 | 0 | 0 | 22 | 2 | 0 | 0 | 8 | 0 | 0 | 1 | 1 | 1 | 3 | 13 | 3 | 0 | 1 | 1 | 0 | 11 | 0 | 2 | 1 | 0 | 0 | 1 | 0 |
| CCMB803 | *morsitans* | 0 | 2 | 1 | 0 | 0 | 26 | 1 | 0 | 0 | 7 | 1 | 0 | 1 | 1 | 1 | 3 | 13 | 3 | 0 | 0 | 1 | 0 | 3 | 0 | 3 | 1 | 0 | 1 | 1 | 1 |
| CCMB805 | *morsitans* | 0 | 0 | 1 | 0 | 0 | 32 | 2 | 0 | 0 | 5 | 0 | 0 | 1 | 1 | 2 | 3 | 16 | 5 | 0 | 2 | 1 | 0 | 7 | 0 | 2 | 0 | 1 | 0 | 1 | 0 |
| CCMB818 | *morsitans* | 0 | 1 | 1 | 0 | 0 | 29 | 1 | 0 | 0 | 8 | 2 | 0 | 0 | 1 | 1 | 3 | 12 | 5 | 0 | 1 | 0 | 0 | 7 | 0 | 2 | 0 | 0 | 0 | 1 | 0 |
| CCMB845 | *morsitans* | 0 | 0 | 1 | 0 | 1 | 35 | 2 | 0 | 0 | 13 | 0 | 0 | 1 | 1 | 2 | 3 | 18 | 5 | 0 | 1 | 0 | 0 | 7 | 0 | 3 | 0 | 0 | 0 | 1 | 0 |
| CCMB1245 | *hardwickei* | 0 | 0 | 2 | 0 | 0 | 39 | 1 | 0 | 0 | 4 | 0 | 1 | 1 | 1 | 1 | 2 | 28 | 1 | 0 | 0 | 0 | 0 | 4 | 0 | 0 | 14 | 0 | 0 | 1 | 0 |
| CCMB1691 | *hardwickei* | 0 | 0 | 2 | 0 | 0 | 42 | 1 | 0 | 0 | 3 | 0 | 0 | 1 | 1 | 1 | 2 | 23 | 1 | 0 | 0 | 0 | 1 | 3 | 0 | 1 | 14 | 0 | 0 | 1 | 0 |
| CCMB1692 | *hardwickei* | 0 | 0 | 1 | 0 | 0 | 36 | 1 | 0 | 0 | 3 | 0 | 0 | 1 | 0 | 1 | 1 | 19 | 1 | 0 | 0 | 0 | 0 | 3 | 0 | 1 | 15 | 0 | 0 | 0 | 0 |
| CCMB1776 | *hardwickei* | 0 | 11 | 0 | 1 | 1 | 29 | 3 | 0 | 2 | 3 | 5 | 0 | 2 | 0 | 1 | 1 | 20 | 1 | 0 | 1 | 0 | 0 | 3 | 0 | 1 | 14 | 0 | 1 | 1 | 1 |
| CCMB2194 | *hardwickei* | 0 | 0 | 1 | 0 | 0 | 24 | 1 | 0 | 0 | 3 | 0 | 1 | 0 | 0 | 1 | 2 | 12 | 0 | 0 | 0 | 1 | 0 | 4 | 0 | 3 | 18 | 0 | 0 | 0 | 1 |
| CCMB2197 | *hardwickei* | 0 | 0 | 0 | 0 | 0 | 25 | 1 | 0 | 0 | 4 | 0 | 0 | 0 | 0 | 1 | 1 | 13 | 0 | 0 | 0 | 1 | 1 | 4 | 0 | 2 | 13 | 0 | 0 | 0 | 0 |
| CCMB3125 | *hardwickei* | 0 | 0 | 2 | 0 | 0 | 28 | 1 | 0 | 0 | 3 | 0 | 0 | 0 | 0 | 1 | 1 | 15 | 3 | 0 | 0 | 0 | 0 | 4 | 0 | 1 | 13 | 0 | 0 | 0 | 0 |
| CCMB3279 | *hardwickei* | 0 | 0 | 0 | 0 | 0 | 21 | 1 | 0 | 0 | 3 | 0 | 0 | 0 | 0 | 1 | 2 | 13 | 0 | 0 | 0 | 1 | 0 | 4 | 0 | 2 | 11 | 0 | 0 | 0 | 1 |
| CCMB3280 | *hardwickei* | 0 | 9 | 0 | 0 | 1 | 13 | 3 | 0 | 4 | 2 | 1 | 0 | 1 | 0 | 0 | 2 | 12 | 0 | 0 | 1 | 1 | 0 | 4 | 0 | 0 | 6 | 0 | 0 | 1 | 1 |
| CCMB3281 | *hardwickei* | 0 | 4 | 1 | 0 | 0 | 23 | 3 | 0 | 2 | 3 | 1 | 0 | 0 | 0 | 1 | 2 | 14 | 0 | 0 | 1 | 1 | 0 | 4 | 0 | 2 | 10 | 4 | 1 | 1 | 2 |
| CCMB3338 | *hardwickei* | 0 | 6 | 1 | 0 | 0 | 16 | 0 | 0 | 2 | 4 | 0 | 0 | 0 | 0 | 1 | 2 | 13 | 0 | 0 | 1 | 0 | 0 | 3 | 0 | 1 | 6 | 0 | 0 | 0 | 0 |

**Continued**

| ID | species | Kazal_domain | LDLA | Leucine-rich_repeat | Lipid-transport_protein | Lysosomal_lipase | Lysozyme C | M12A | MCSACoa | Neprilysin | NOT_TOXIN_Actin | NOT_TOXIN_Arginine_kinase_ | NOT_TOXIN_ATP_synthase | NOT_TOXIN_Calcium-transporting_ATPase | NOT_TOXIN_Histone_H2A | NOT_TOXIN_Histone_H2B | NOT_TOXIN_INSUFFICIENT_EVIDENCE_Alpha-L-Fucosidase | NOT_TOXIN_INSUFFICIENT_EVIDENCE_Avidin-related | NOT_TOXIN_INSUFFICIENT_EVIDENCE_CUB-tolloid | NOT_TOXIN_INSUFFICIENT_EVIDENCE_NOT_TOXIN_INSUFFICIENT_EVIDENCE_insect_cuticle_uncharacterized_protein_ | NOT_TOXIN_Myosin_heavy_chain_protein | NOT_TOXIN_Myosin_light_chain_protein | NOT_TOXIN_Myosin_tail_1_domain-containing_protein | NOT_TOXIN_NO_SIGNALP_Calponin | NOT_TOXIN_NO_SIGNALP_Pyruvate_kinase | NOT_TOXIN_structural_component | NOT_TOXIN_Tropomyosin_ | NOT_TOXIN_Troponin | NOT_TOXIN_tubulin_alpha_chain_ | NOT_TOXIN_tubulin_beta_chain_ | NOT_TOXIN_Vitellogenin-6 |
|---|---|---|---|---|---|---|---|---|---|---|---|---|---|---|---|---|---|---|---|---|---|---|---|---|---|---|---|---|---|---|---|
| CCMB1159 | *morsitans* | 0 | 13 | 0 | 7 | 0 | 2 | 0 | 0 | 0 | 0 | 0 | 0 | 0 | 0 | 0 | 0 | 0 | 3 | 0 | 0 | 0 | 0 | 0 | 0 | 0 | 0 | 0 | 0 | 0 | 2 |
| CCMB1209 | *morsitans* | 0 | 11 | 0 | 1 | 0 | 2 | 0 | 0 | 0 | 0 | 0 | 0 | 1 | 0 | 0 | 0 | 0 | 4 | 0 | 0 | 0 | 0 | 0 | 0 | 0 | 0 | 0 | 0 | 0 | 2 |
| CCMB1325 | *morsitans* | 0 | 15 | 0 | 0 | 0 | 2 | 0 | 0 | 0 | 0 | 0 | 0 | 0 | 0 | 0 | 0 | 0 | 2 | 0 | 0 | 0 | 0 | 0 | 0 | 0 | 0 | 0 | 0 | 0 | 1 |
| CCMB1693 | *morsitans* | 0 | 19 | 0 | 9 | 0 | 3 | 0 | 0 | 0 | 0 | 0 | 0 | 0 | 0 | 1 | 0 | 1 | 6 | 0 | 0 | 0 | 0 | 0 | 0 | 0 | 0 | 0 | 0 | 0 | 7 |
| CCMB1695 | *morsitans* | 0 | 14 | 0 | 0 | 0 | 3 | 0 | 0 | 0 | 0 | 0 | 0 | 0 | 0 | 0 | 1 | 0 | 1 | 0 | 0 | 0 | 0 | 0 | 0 | 0 | 0 | 0 | 0 | 0 | 0 |
| CCMB1696 | *morsitans* | 0 | 17 | 0 | 0 | 0 | 2 | 0 | 0 | 0 | 0 | 0 | 0 | 0 | 0 | 0 | 0 | 0 | 2 | 1 | 0 | 0 | 0 | 0 | 0 | 0 | 0 | 0 | 0 | 0 | 1 |
| CCMB1725 | *morsitans* | 0 | 11 | 0 | 11 | 0 | 2 | 0 | 0 | 1 | 6 | 3 | 0 | 0 | 1 | 1 | 1 | 0 | 1 | 0 | 1 | 2 | 1 | 0 | 1 | 0 | 0 | 0 | 1 | 0 | 8 |
| CCMB1766 | *morsitans* | 0 | 16 | 0 | 0 | 0 | 4 | 0 | 0 | 0 | 0 | 0 | 0 | 0 | 0 | 0 | 0 | 0 | 5 | 0 | 0 | 0 | 0 | 0 | 0 | 0 | 0 | 0 | 0 | 0 | 2 |
| CCMB1769 | *morsitans* | 0 | 17 | 1 | 4 | 0 | 3 | 0 | 0 | 0 | 0 | 0 | 0 | 0 | 0 | 0 | 0 | 0 | 3 | 0 | 0 | 0 | 0 | 0 | 0 | 0 | 0 | 0 | 1 | 0 | 3 |
| CCMB1803 | *morsitans* | 0 | 13 | 0 | 8 | 0 | 2 | 0 | 0 | 0 | 1 | 0 | 0 | 0 | 0 | 0 | 0 | 0 | 1 | 0 | 0 | 0 | 0 | 0 | 0 | 1 | 0 | 0 | 0 | 0 | 6 |
| CCMB2184 | *morsitans* | 0 | 19 | 1 | 1 | 0 | 2 | 0 | 0 | 0 | 2 | 0 | 0 | 0 | 0 | 0 | 0 | 0 | 3 | 0 | 0 | 0 | 0 | 0 | 0 | 0 | 0 | 0 | 0 | 0 | 3 |
| CCMB2198 | *morsitans* | 0 | 14 | 0 | 0 | 0 | 2 | 0 | 0 | 0 | 0 | 0 | 0 | 0 | 0 | 0 | 0 | 0 | 4 | 0 | 0 | 0 | 0 | 0 | 0 | 0 | 0 | 0 | 0 | 0 | 0 |
| CCMB2382 | *morsitans* | 0 | 10 | 0 | 7 | 0 | 2 | 0 | 0 | 1 | 2 | 0 | 0 | 0 | 0 | 0 | 1 | 0 | 2 | 0 | 0 | 0 | 0 | 0 | 0 | 0 | 0 | 0 | 0 | 0 | 6 |
| CCMB3265 | *morsitans* | 1 | 10 | 0 | 7 | 0 | 2 | 0 | 0 | 1 | 3 | 0 | 1 | 0 | 1 | 1 | 1 | 0 | 2 | 0 | 1 | 1 | 0 | 0 | 0 | 0 | 0 | 0 | 1 | 1 | 9 |
| CCMB3772 | *morsitans* | 0 | 9 | 2 | 10 | 0 | 3 | 0 | 0 | 1 | 5 | 3 | 0 | 0 | 0 | 0 | 1 | 0 | 3 | 0 | 0 | 0 | 0 | 0 | 1 | 0 | 0 | 0 | 0 | 0 | 8 |
| CCMB3868 | *morsitans* | 0 | 13 | 0 | 1 | 0 | 3 | 0 | 0 | 0 | 0 | 0 | 0 | 0 | 0 | 0 | 2 | 0 | 1 | 0 | 0 | 0 | 0 | 0 | 0 | 0 | 0 | 1 | 0 | 0 | 6 |
| CCMB41 | *morsitans* | 0 | 7 | 0 | 11 | 0 | 0 | 0 | 0 | 0 | 1 | 0 | 0 | 0 | 1 | 1 | 0 | 0 | 1 | 0 | 0 | 0 | 0 | 0 | 0 | 0 | 0 | 0 | 0 | 0 | 7 |
| CCMB4114 | *morsitans* | 0 | 13 | 0 | 0 | 0 | 3 | 0 | 0 | 0 | 0 | 0 | 0 | 0 | 0 | 0 | 0 | 0 | 3 | 0 | 0 | 0 | 0 | 0 | 0 | 0 | 0 | 0 | 0 | 0 | 0 |
| CCMB4216 | *morsitans* | 0 | 12 | 1 | 9 | 0 | 3 | 0 | 0 | 0 | 3 | 0 | 0 | 0 | 1 | 0 | 4 | 0 | 2 | 0 | 0 | 0 | 0 | 0 | 0 | 0 | 0 | 0 | 0 | 0 | 9 |
| CCMB4224 | *morsitans* | 0 | 12 | 1 | 8 | 0 | 3 | 0 | 0 | 0 | 0 | 0 | 0 | 0 | 0 | 0 | 3 | 0 | 3 | 0 | 0 | 0 | 0 | 0 | 0 | 0 | 0 | 0 | 0 | 0 | 8 |
| CCMB4407 | *morsitans* | 0 | 13 | 0 | 10 | 0 | 2 | 0 | 0 | 0 | 3 | 0 | 0 | 0 | 0 | 0 | 2 | 0 | 2 | 0 | 0 | 0 | 0 | 0 | 0 | 0 | 0 | 0 | 0 | 0 | 9 |
| CCMB4441 | *morsitans* | 0 | 16 | 0 | 4 | 0 | 2 | 0 | 0 | 0 | 0 | 0 | 0 | 0 | 0 | 0 | 3 | 0 | 2 | 0 | 0 | 0 | 0 | 0 | 0 | 0 | 0 | 0 | 0 | 0 | 5 |
| CCMB4454 | *morsitans* | 0 | 10 | 0 | 1 | 0 | 3 | 0 | 0 | 0 | 0 | 0 | 0 | 0 | 0 | 0 | 0 | 0 | 2 | 0 | 0 | 0 | 0 | 0 | 0 | 0 | 0 | 0 | 0 | 0 | 3 |
| CCMB752 | *morsitans* | 0 | 10 | 2 | 9 | 0 | 2 | 0 | 0 | 1 | 1 | 0 | 0 | 0 | 1 | 1 | 0 | 0 | 4 | 0 | 2 | 0 | 0 | 0 | 0 | 0 | 0 | 1 | 0 | 0 | 5 |
| CCMB803 | *morsitans* | 0 | 18 | 0 | 8 | 0 | 2 | 0 | 0 | 0 | 0 | 0 | 0 | 0 | 0 | 0 | 0 | 0 | 3 | 0 | 0 | 0 | 0 | 0 | 0 | 0 | 0 | 0 | 0 | 0 | 6 |
| CCMB805 | *morsitans* | 0 | 14 | 1 | 0 | 0 | 2 | 0 | 0 | 0 | 0 | 0 | 0 | 0 | 0 | 0 | 0 | 0 | 2 | 0 | 0 | 0 | 0 | 0 | 0 | 0 | 0 | 0 | 0 | 0 | 0 |
| CCMB818 | *morsitans* | 0 | 14 | 0 | 0 | 0 | 2 | 1 | 0 | 0 | 0 | 0 | 0 | 0 | 0 | 0 | 0 | 0 | 2 | 0 | 0 | 0 | 0 | 0 | 0 | 0 | 0 | 0 | 0 | 0 | 2 |
| CCMB845 | *morsitans* | 0 | 16 | 0 | 0 | 0 | 2 | 0 | 0 | 0 | 0 | 0 | 0 | 0 | 0 | 0 | 0 | 0 | 1 | 0 | 0 | 0 | 0 | 0 | 0 | 0 | 0 | 2 | 0 | 0 | 0 |
| CCMB1245 | *hardwickei* | 0 | 15 | 0 | 0 | 0 | 3 | 0 | 1 | 0 | 0 | 0 | 0 | 0 | 0 | 1 | 0 | 0 | 2 | 0 | 0 | 0 | 0 | 0 | 0 | 0 | 0 | 0 | 0 | 0 | 0 |
| CCMB1691 | *hardwickei* | 0 | 15 | 0 | 0 | 0 | 3 | 0 | 0 | 0 | 0 | 0 | 0 | 0 | 0 | 0 | 0 | 0 | 0 | 0 | 0 | 0 | 0 | 0 | 0 | 0 | 0 | 0 | 0 | 0 | 1 |
| CCMB1692 | *hardwickei* | 0 | 13 | 0 | 0 | 0 | 3 | 0 | 0 | 0 | 0 | 0 | 0 | 0 | 0 | 0 | 0 | 0 | 0 | 0 | 0 | 0 | 0 | 0 | 0 | 0 | 0 | 0 | 0 | 0 | 0 |
| CCMB1776 | *hardwickei* | 0 | 16 | 0 | 7 | 1 | 3 | 0 | 0 | 1 | 0 | 0 | 0 | 0 | 0 | 0 | 1 | 0 | 0 | 0 | 1 | 0 | 0 | 0 | 0 | 0 | 0 | 0 | 0 | 0 | 3 |
| CCMB2194 | *hardwickei* | 0 | 9 | 0 | 0 | 0 | 0 | 0 | 0 | 0 | 0 | 0 | 0 | 0 | 0 | 0 | 0 | 0 | 0 | 0 | 0 | 0 | 0 | 0 | 0 | 0 | 0 | 0 | 0 | 0 | 0 |
| CCMB2197 | *hardwickei* | 0 | 7 | 0 | 0 | 0 | 0 | 0 | 0 | 0 | 0 | 1 | 0 | 0 | 0 | 0 | 0 | 0 | 0 | 0 | 0 | 0 | 0 | 0 | 0 | 0 | 0 | 0 | 0 | 0 | 0 |
| CCMB3125 | *hardwickei* | 0 | 11 | 0 | 0 | 0 | 3 | 0 | 0 | 0 | 0 | 0 | 0 | 0 | 0 | 0 | 0 | 0 | 0 | 0 | 0 | 0 | 0 | 0 | 0 | 0 | 0 | 0 | 0 | 0 | 2 |
| CCMB3279 | *hardwickei* | 0 | 9 | 0 | 0 | 0 | 0 | 0 | 0 | 0 | 0 | 0 | 0 | 0 | 0 | 0 | 0 | 0 | 0 | 0 | 0 | 0 | 0 | 0 | 0 | 0 | 0 | 0 | 0 | 0 | 0 |
| CCMB3280 | *hardwickei* | 0 | 6 | 0 | 9 | 0 | 0 | 0 | 0 | 1 | 3 | 0 | 0 | 0 | 0 | 1 | 0 | 0 | 0 | 0 | 0 | 1 | 0 | 1 | 0 | 0 | 0 | 0 | 1 | 1 | 4 |
| CCMB3281 | *hardwickei* | 0 | 11 | 0 | 9 | 0 | 0 | 0 | 0 | 1 | 5 | 0 | 0 | 0 | 1 | 1 | 0 | 0 | 0 | 0 | 0 | 1 | 0 | 0 | 0 | 0 | 1 | 0 | 0 | 1 | 2 |
| CCMB3338 | *hardwickei* | 0 | 3 | 0 | 7 | 0 | 0 | 0 | 0 | 1 | 0 | 0 | 0 | 0 | 0 | 0 | 0 | 0 | 0 | 0 | 0 | 0 | 0 | 0 | 0 | 0 | 0 | 0 | 0 | 0 | 3 |

**Continued**

| ID | species | PAM-like | PEBP-like | Peptidase_M13 | Phosphodiesterase2 | PLA2 | Purple acid phosphatase | S1 | S10 | S8 | Saposin-related | SCTX01 | SCTX02 | Serpine | SLPTx-SSD552 | SLPTx-SSD558 | SLPTx-SSD976 | SLPTX01 | SLPTX02 | SLPTX03 | SLPTX04 | SLPTX05 | SLPTX06 | SLPTX07 | SLPTX08 | SLPTX09 | SLPTX10 | SLPTX11 | SLPTX12 | SLPTX13 | SLPTX14 |
|---|---|---|---|---|---|---|---|---|---|---|---|---|---|---|---|---|---|---|---|---|---|---|---|---|---|---|---|---|---|---|---|
| CCMB1159 | *morsitans* | 1 | 2 | 0 | 1 | 1 | 0 | 3 | 2 | 1 | 0 | 0 | 0 | 0 | 0 | 0 | 0 | 2 | 0 | 1 | 4 | 0 | 2 | 1 | 0 | 1 | 5 | 9 | 1 | 2 | 1 |
| CCMB1209 | *morsitans* | 2 | 2 | 0 | 1 | 4 | 0 | 8 | 1 | 2 | 0 | 0 | 0 | 1 | 0 | 0 | 0 | 2 | 0 | 1 | 3 | 0 | 0 | 1 | 0 | 1 | 3 | 9 | 1 | 2 | 0 |
| CCMB1325 | *morsitans* | 2 | 2 | 0 | 6 | 2 | 0 | 7 | 1 | 2 | 0 | 0 | 0 | 0 | 0 | 0 | 1 | 2 | 0 | 2 | 2 | 1 | 0 | 0 | 0 | 0 | 6 | 17 | 1 | 0 | 8 |
| CCMB1693 | *morsitans* | 1 | 1 | 1 | 2 | 2 | 0 | 6 | 1 | 2 | 0 | 0 | 1 | 1 | 0 | 0 | 0 | 3 | 0 | 1 | 2 | 0 | 2 | 0 | 0 | 0 | 4 | 11 | 1 | 3 | 7 |
| CCMB1695 | *morsitans* | 1 | 0 | 0 | 1 | 3 | 0 | 7 | 1 | 1 | 0 | 0 | 0 | 0 | 0 | 0 | 0 | 2 | 0 | 1 | 4 | 0 | 0 | 1 | 1 | 0 | 4 | 10 | 1 | 3 | 0 |
| CCMB1696 | *morsitans* | 1 | 1 | 0 | 1 | 2 | 0 | 6 | 1 | 6 | 1 | 0 | 0 | 1 | 0 | 0 | 0 | 2 | 0 | 3 | 4 | 0 | 0 | 0 | 1 | 0 | 3 | 17 | 0 | 3 | 0 |
| CCMB1725 | *morsitans* | 0 | 1 | 0 | 1 | 2 | 0 | 9 | 0 | 0 | 1 | 0 | 0 | 3 | 0 | 0 | 0 | 1 | 0 | 1 | 2 | 0 | 0 | 1 | 0 | 0 | 4 | 13 | 0 | 3 | 0 |
| CCMB1766 | *morsitans* | 1 | 1 | 0 | 1 | 3 | 0 | 6 | 1 | 2 | 0 | 0 | 0 | 1 | 0 | 1 | 0 | 3 | 0 | 1 | 3 | 0 | 2 | 0 | 0 | 0 | 5 | 9 | 1 | 0 | 0 |
| CCMB1769 | *morsitans* | 1 | 1 | 0 | 1 | 2 | 0 | 8 | 1 | 1 | 1 | 0 | 0 | 0 | 0 | 0 | 0 | 2 | 0 | 2 | 4 | 0 | 2 | 1 | 0 | 0 | 4 | 15 | 1 | 3 | 7 |
| CCMB1803 | *morsitans* | 1 | 2 | 0 | 1 | 1 | 0 | 7 | 1 | 1 | 0 | 0 | 0 | 1 | 0 | 0 | 0 | 2 | 1 | 1 | 1 | 0 | 0 | 1 | 0 | 1 | 2 | 9 | 1 | 0 | 0 |
| CCMB2184 | *morsitans* | 1 | 2 | 0 | 1 | 2 | 1 | 3 | 2 | 2 | 1 | 0 | 0 | 0 | 0 | 0 | 1 | 3 | 0 | 1 | 2 | 0 | 2 | 1 | 0 | 0 | 2 | 11 | 1 | 0 | 0 |
| CCMB2198 | *morsitans* | 1 | 0 | 0 | 1 | 1 | 0 | 6 | 1 | 1 | 0 | 0 | 0 | 1 | 0 | 0 | 1 | 3 | 0 | 1 | 1 | 0 | 0 | 1 | 0 | 0 | 1 | 8 | 1 | 0 | 0 |
| CCMB2382 | *morsitans* | 1 | 0 | 1 | 1 | 2 | 0 | 12 | 2 | 1 | 0 | 0 | 0 | 3 | 0 | 0 | 0 | 2 | 0 | 3 | 4 | 0 | 0 | 1 | 0 | 0 | 4 | 15 | 0 | 3 | 9 |
| CCMB3265 | *morsitans* | 0 | 0 | 1 | 1 | 3 | 0 | 7 | 0 | 0 | 0 | 0 | 0 | 2 | 0 | 0 | 1 | 3 | 1 | 2 | 1 | 0 | 0 | 1 | 0 | 0 | 1 | 10 | 0 | 0 | 0 |
| CCMB3772 | *morsitans* | 1 | 2 | 3 | 2 | 1 | 0 | 11 | 1 | 2 | 0 | 0 | 0 | 3 | 0 | 1 | 0 | 2 | 1 | 1 | 1 | 0 | 0 | 1 | 0 | 0 | 5 | 10 | 1 | 0 | 0 |
| CCMB3868 | *morsitans* | 1 | 2 | 0 | 2 | 2 | 0 | 7 | 2 | 1 | 0 | 0 | 1 | 0 | 0 | 0 | 0 | 4 | 1 | 1 | 1 | 0 | 0 | 0 | 1 | 0 | 3 | 6 | 1 | 0 | 0 |
| CCMB41 | *morsitans* | 0 | 0 | 0 | 0 | 0 | 0 | 4 | 0 | 0 | 0 | 0 | 0 | 1 | 0 | 0 | 0 | 1 | 0 | 1 | 0 | 0 | 0 | 0 | 0 | 0 | 1 | 2 | 0 | 0 | 0 |
| CCMB4114 | *morsitans* | 1 | 3 | 0 | 2 | 2 | 0 | 6 | 2 | 2 | 1 | 0 | 0 | 1 | 0 | 0 | 0 | 2 | 0 | 3 | 2 | 0 | 0 | 0 | 0 | 0 | 1 | 16 | 0 | 0 | 0 |
| CCMB4216 | *morsitans* | 1 | 2 | 1 | 2 | 2 | 0 | 10 | 1 | 1 | 0 | 0 | 0 | 3 | 0 | 0 | 0 | 2 | 0 | 1 | 1 | 1 | 0 | 0 | 0 | 0 | 3 | 7 | 0 | 0 | 0 |
| CCMB4224 | *morsitans* | 1 | 2 | 2 | 2 | 2 | 0 | 9 | 1 | 1 | 0 | 0 | 0 | 4 | 0 | 0 | 0 | 2 | 0 | 1 | 1 | 0 | 0 | 0 | 0 | 0 | 3 | 8 | 0 | 0 | 0 |
| CCMB4407 | *morsitans* | 1 | 2 | 2 | 2 | 2 | 0 | 9 | 1 | 1 | 0 | 0 | 0 | 1 | 0 | 0 | 0 | 2 | 1 | 1 | 0 | 0 | 0 | 0 | 0 | 0 | 3 | 14 | 0 | 0 | 0 |
| CCMB4441 | *morsitans* | 1 | 2 | 0 | 2 | 2 | 0 | 7 | 1 | 1 | 0 | 0 | 0 | 0 | 0 | 0 | 0 | 3 | 0 | 1 | 0 | 1 | 0 | 0 | 0 | 0 | 2 | 14 | 0 | 0 | 0 |
| CCMB4454 | *morsitans* | 1 | 2 | 0 | 2 | 1 | 0 | 5 | 0 | 0 | 0 | 0 | 0 | 0 | 0 | 0 | 0 | 2 | 0 | 1 | 0 | 0 | 0 | 0 | 0 | 0 | 1 | 6 | 0 | 0 | 0 |
| CCMB752 | *morsitans* | 1 | 2 | 3 | 0 | 2 | 0 | 7 | 2 | 1 | 0 | 0 | 0 | 0 | 0 | 0 | 1 | 3 | 0 | 1 | 2 | 0 | 0 | 0 | 0 | 0 | 2 | 9 | 1 | 0 | 1 |
| CCMB803 | *morsitans* | 1 | 1 | 1 | 1 | 3 | 0 | 3 | 1 | 2 | 0 | 0 | 0 | 0 | 0 | 0 | 0 | 2 | 0 | 3 | 4 | 0 | 0 | 0 | 0 | 2 | 6 | 18 | 0 | 3 | 3 |
| CCMB805 | *morsitans* | 2 | 1 | 0 | 1 | 1 | 0 | 6 | 1 | 1 | 1 | 1 | 0 | 1 | 0 | 0 | 0 | 2 | 0 | 1 | 4 | 0 | 2 | 1 | 0 | 2 | 6 | 12 | 1 | 0 | 1 |
| CCMB818 | *morsitans* | 1 | 0 | 0 | 1 | 1 | 0 | 6 | 1 | 2 | 0 | 0 | 2 | 0 | 0 | 0 | 0 | 3 | 0 | 1 | 4 | 0 | 2 | 1 | 0 | 2 | 5 | 12 | 1 | 3 | 0 |
| CCMB845 | *morsitans* | 1 | 0 | 0 | 1 | 2 | 0 | 6 | 1 | 1 | 0 | 0 | 0 | 0 | 1 | 0 | 0 | 2 | 0 | 1 | 3 | 0 | 2 | 1 | 0 | 2 | 7 | 10 | 1 | 3 | 1 |
| CCMB1245 | *hardwickei* | 1 | 1 | 1 | 0 | 7 | 0 | 4 | 1 | 0 | 0 | 0 | 0 | 0 | 0 | 0 | 0 | 2 | 0 | 0 | 3 | 0 | 0 | 0 | 0 | 0 | 4 | 3 | 1 | 1 | 2 |
| CCMB1691 | *hardwickei* | 2 | 1 | 0 | 0 | 6 | 0 | 5 | 1 | 2 | 0 | 0 | 0 | 0 | 0 | 0 | 0 | 2 | 0 | 1 | 0 | 1 | 0 | 0 | 1 | 0 | 6 | 3 | 1 | 1 | 1 |
| CCMB1692 | *hardwickei* | 1 | 1 | 1 | 0 | 4 | 0 | 6 | 1 | 1 | 0 | 0 | 0 | 0 | 0 | 0 | 0 | 2 | 0 | 1 | 3 | 0 | 0 | 0 | 0 | 0 | 6 | 3 | 1 | 1 | 1 |
| CCMB1776 | *hardwickei* | 1 | 1 | 4 | 0 | 6 | 0 | 9 | 1 | 2 | 0 | 0 | 2 | 6 | 0 | 0 | 1 | 2 | 0 | 1 | 0 | 1 | 0 | 0 | 0 | 0 | 4 | 3 | 0 | 1 | 0 |
| CCMB2194 | *hardwickei* | 1 | 1 | 0 | 0 | 4 | 0 | 4 | 1 | 1 | 0 | 0 | 0 | 0 | 0 | 0 | 0 | 2 | 0 | 1 | 0 | 0 | 0 | 0 | 1 | 0 | 3 | 4 | 1 | 0 | 1 |
| CCMB2197 | *hardwickei* | 1 | 1 | 2 | 0 | 2 | 0 | 4 | 0 | 0 | 0 | 0 | 0 | 0 | 0 | 0 | 0 | 2 | 0 | 1 | 0 | 0 | 0 | 0 | 0 | 0 | 3 | 4 | 1 | 0 | 1 |
| CCMB3125 | *hardwickei* | 1 | 1 | 1 | 0 | 5 | 0 | 9 | 1 | 1 | 0 | 0 | 0 | 0 | 0 | 0 | 0 | 2 | 0 | 1 | 1 | 0 | 0 | 0 | 0 | 0 | 8 | 3 | 1 | 1 | 1 |
| CCMB3279 | *hardwickei* | 1 | 1 | 0 | 0 | 3 | 0 | 4 | 1 | 0 | 0 | 0 | 0 | 0 | 0 | 0 | 0 | 1 | 0 | 0 | 0 | 0 | 0 | 0 | 1 | 0 | 3 | 4 | 1 | 0 | 1 |
| CCMB3280 | *hardwickei* | 0 | 0 | 1 | 0 | 3 | 0 | 9 | 0 | 0 | 0 | 0 | 1 | 5 | 0 | 0 | 0 | 1 | 0 | 0 | 0 | 0 | 0 | 0 | 0 | 0 | 0 | 3 | 0 | 0 | 1 |
| CCMB3281 | *hardwickei* | 2 | 0 | 2 | 0 | 3 | 0 | 4 | 1 | 0 | 0 | 0 | 0 | 0 | 0 | 0 | 0 | 1 | 0 | 0 | 0 | 0 | 0 | 1 | 1 | 0 | 3 | 4 | 1 | 0 | 1 |
| CCMB3338 | *hardwickei* | 0 | 0 | 1 | 0 | 2 | 0 | 2 | 0 | 1 | 0 | 0 | 1 | 1 | 0 | 0 | 0 | 1 | 0 | 0 | 0 | 0 | 0 | 0 | 1 | 0 | 4 | 4 | 0 | 0 | 1 |

**Continued**

| ID | species | SLPTX15 | SLPTX16 | SLPTX17 | SLPTX19 | SLPTX20 | SLPTX21 | SLPTX29 | SLPTX30 | SLPTX47 | TGFbeta | Thioester-containing protein 4 | Transferrin | Triacylglycerol_lipase_family | Unchar03 | Unchar05 | Unchar06 | Unchar08 | Unchar10 | Unchar11 | Unchar13 | Unchar30 | Unchar31 | Unchar32 | Unchar33 | Vigilin | WAP |
|---|---|---|---|---|---|---|---|---|---|---|---|---|---|---|---|---|---|---|---|---|---|---|---|---|---|---|---|
| CCMB1159 | *morsitans* | 11 | 5 | 2 | 0 | 1 | 1 | 0 | 0 | 0 | 1 | 0 | 2 | 0 | 0 | 7 | 1 | 0 | 3 | 3 | 0 | 0 | 0 | 0 | 0 | 0 | 1 |
| CCMB1209 | *morsitans* | 9 | 4 | 1 | 1 | 2 | 0 | 0 | 0 | 0 | 1 | 0 | 0 | 1 | 0 | 5 | 1 | 1 | 2 | 0 | 0 | 0 | 0 | 0 | 0 | 0 | 0 |
| CCMB1325 | *morsitans* | 13 | 4 | 1 | 1 | 1 | 2 | 0 | 0 | 0 | 0 | 0 | 0 | 0 | 0 | 4 | 1 | 0 | 2 | 1 | 0 | 1 | 0 | 0 | 0 | 0 | 2 |
| CCMB1693 | *morsitans* | 10 | 5 | 4 | 2 | 1 | 1 | 0 | 0 | 0 | 1 | 0 | 7 | 4 | 0 | 4 | 1 | 1 | 2 | 3 | 0 | 0 | 0 | 0 | 0 | 0 | 0 |
| CCMB1695 | *morsitans* | 11 | 6 | 0 | 1 | 1 | 0 | 0 | 1 | 0 | 1 | 0 | 4 | 3 | 0 | 3 | 1 | 0 | 2 | 2 | 0 | 0 | 0 | 0 | 0 | 0 | 0 |
| CCMB1696 | *morsitans* | 13 | 4 | 2 | 2 | 1 | 1 | 0 | 0 | 0 | 1 | 0 | 0 | 1 | 0 | 4 | 1 | 1 | 2 | 1 | 0 | 2 | 0 | 0 | 0 | 0 | 0 |
| CCMB1725 | *morsitans* | 14 | 2 | 2 | 1 | 1 | 0 | 0 | 4 | 0 | 1 | 0 | 9 | 0 | 0 | 1 | 1 | 0 | 2 | 3 | 0 | 1 | 0 | 2 | 1 | 0 | 0 |
| CCMB1766 | *morsitans* | 12 | 5 | 9 | 2 | 1 | 0 | 0 | 0 | 0 | 1 | 0 | 0 | 0 | 0 | 4 | 1 | 1 | 2 | 3 | 0 | 0 | 0 | 0 | 0 | 0 | 0 |
| CCMB1769 | *morsitans* | 15 | 4 | 2 | 1 | 1 | 1 | 0 | 0 | 0 | 1 | 0 | 1 | 0 | 0 | 5 | 1 | 1 | 2 | 1 | 0 | 1 | 0 | 0 | 0 | 0 | 0 |
| CCMB1803 | *morsitans* | 7 | 4 | 0 | 1 | 0 | 0 | 0 | 2 | 0 | 1 | 0 | 1 | 2 | 0 | 4 | 1 | 1 | 3 | 4 | 0 | 0 | 0 | 1 | 0 | 0 | 0 |
| CCMB2184 | *morsitans* | 13 | 5 | 4 | 1 | 0 | 1 | 0 | 1 | 0 | 0 | 0 | 1 | 2 | 0 | 2 | 2 | 1 | 3 | 1 | 0 | 0 | 0 | 0 | 0 | 0 | 0 |
| CCMB2198 | *morsitans* | 7 | 3 | 3 | 1 | 0 | 1 | 0 | 0 | 0 | 0 | 0 | 0 | 1 | 0 | 3 | 1 | 1 | 2 | 2 | 0 | 0 | 0 | 0 | 0 | 0 | 0 |
| CCMB2382 | *morsitans* | 8 | 3 | 0 | 1 | 0 | 0 | 0 | 0 | 0 | 1 | 0 | 7 | 0 | 0 | 2 | 1 | 0 | 2 | 2 | 0 | 1 | 0 | 1 | 0 | 0 | 0 |
| CCMB3265 | *morsitans* | 12 | 6 | 1 | 1 | 1 | 1 | 0 | 2 | 0 | 0 | 1 | 8 | 0 | 1 | 3 | 1 | 1 | 2 | 2 | 0 | 0 | 0 | 2 | 1 | 0 | 0 |
| CCMB3772 | *morsitans* | 8 | 5 | 1 | 1 | 0 | 1 | 0 | 2 | 1 | 0 | 0 | 8 | 0 | 0 | 2 | 1 | 1 | 3 | 4 | 0 | 0 | 0 | 2 | 0 | 0 | 0 |
| CCMB3868 | *morsitans* | 8 | 5 | 0 | 0 | 0 | 0 | 1 | 0 | 0 | 0 | 0 | 2 | 0 | 0 | 3 | 1 | 0 | 1 | 1 | 0 | 0 | 0 | 0 | 0 | 0 | 0 |
| CCMB41 | *morsitans* | 6 | 5 | 0 | 0 | 0 | 0 | 0 | 2 | 0 | 0 | 0 | 3 | 0 | 0 | 0 | 0 | 0 | 1 | 0 | 0 | 0 | 0 | 0 | 2 | 0 | 0 |
| CCMB4114 | *morsitans* | 11 | 3 | 1 | 1 | 0 | 0 | 0 | 1 | 0 | 0 | 0 | 0 | 1 | 0 | 3 | 1 | 0 | 1 | 2 | 0 | 1 | 2 | 0 | 0 | 0 | 0 |
| CCMB4216 | *morsitans* | 10 | 3 | 5 | 1 | 0 | 0 | 0 | 2 | 0 | 0 | 0 | 1 | 1 | 0 | 2 | 1 | 1 | 1 | 4 | 0 | 0 | 0 | 0 | 0 | 0 | 0 |
| CCMB4224 | *morsitans* | 12 | 4 | 7 | 1 | 0 | 0 | 0 | 3 | 0 | 0 | 0 | 1 | 2 | 0 | 3 | 1 | 1 | 1 | 1 | 0 | 0 | 0 | 0 | 0 | 0 | 0 |
| CCMB4407 | *morsitans* | 8 | 1 | 0 | 1 | 0 | 0 | 1 | 1 | 0 | 0 | 0 | 7 | 2 | 0 | 2 | 1 | 0 | 1 | 4 | 0 | 0 | 0 | 0 | 0 | 0 | 0 |
| CCMB4441 | *morsitans* | 8 | 2 | 0 | 1 | 1 | 0 | 0 | 0 | 0 | 0 | 0 | 1 | 2 | 0 | 2 | 1 | 0 | 1 | 1 | 0 | 0 | 1 | 0 | 0 | 0 | 0 |
| CCMB4454 | *morsitans* | 7 | 2 | 0 | 0 | 0 | 0 | 0 | 0 | 0 | 0 | 0 | 0 | 1 | 0 | 2 | 1 | 0 | 1 | 1 | 0 | 0 | 0 | 0 | 0 | 1 | 0 |
| CCMB752 | *morsitans* | 13 | 5 | 3 | 1 | 0 | 1 | 0 | 2 | 0 | 0 | 0 | 2 | 0 | 0 | 4 | 2 | 0 | 2 | 5 | 0 | 0 | 0 | 0 | 0 | 0 | 0 |
| CCMB803 | *morsitans* | 24 | 2 | 1 | 1 | 1 | 1 | 0 | 0 | 0 | 1 | 0 | 5 | 0 | 0 | 2 | 1 | 0 | 2 | 2 | 0 | 1 | 2 | 0 | 0 | 0 | 1 |
| CCMB805 | *morsitans* | 15 | 6 | 0 | 0 | 1 | 2 | 0 | 0 | 0 | 1 | 0 | 0 | 1 | 0 | 4 | 0 | 0 | 2 | 2 | 0 | 0 | 0 | 0 | 0 | 0 | 0 |
| CCMB818 | *morsitans* | 14 | 6 | 0 | 1 | 0 | 2 | 0 | 3 | 0 | 1 | 0 | 5 | 1 | 0 | 6 | 1 | 0 | 1 | 2 | 0 | 0 | 0 | 0 | 0 | 0 | 1 |
| CCMB845 | *morsitans* | 15 | 9 | 0 | 1 | 1 | 2 | 0 | 0 | 0 | 1 | 0 | 0 | 2 | 0 | 5 | 0 | 0 | 1 | 2 | 0 | 0 | 0 | 0 | 0 | 0 | 1 |
| CCMB1245 | *hardwickei* | 1 | 5 | 4 | 0 | 0 | 1 | 0 | 1 | 0 | 0 | 0 | 0 | 0 | 0 | 3 | 1 | 0 | 5 | 2 | 0 | 0 | 0 | 0 | 0 | 0 | 0 |
| CCMB1691 | *hardwickei* | 2 | 5 | 4 | 0 | 0 | 1 | 0 | 1 | 0 | 0 | 0 | 0 | 0 | 0 | 3 | 2 | 0 | 3 | 3 | 0 | 0 | 0 | 1 | 0 | 0 | 0 |
| CCMB1692 | *hardwickei* | 1 | 5 | 5 | 0 | 0 | 1 | 2 | 1 | 0 | 0 | 0 | 0 | 0 | 0 | 1 | 1 | 0 | 5 | 4 | 0 | 0 | 0 | 1 | 0 | 0 | 0 |
| CCMB1776 | *hardwickei* | 1 | 8 | 6 | 0 | 0 | 0 | 0 | 5 | 0 | 0 | 1 | 2 | 0 | 0 | 1 | 1 | 0 | 2 | 3 | 0 | 0 | 0 | 1 | 0 | 0 | 0 |
| CCMB2194 | *hardwickei* | 1 | 6 | 2 | 0 | 0 | 0 | 0 | 1 | 0 | 0 | 0 | 0 | 0 | 0 | 1 | 1 | 0 | 3 | 2 | 0 | 0 | 0 | 1 | 0 | 0 | 0 |
| CCMB2197 | *hardwickei* | 1 | 5 | 2 | 0 | 0 | 0 | 0 | 0 | 0 | 0 | 0 | 0 | 0 | 0 | 1 | 1 | 0 | 3 | 2 | 0 | 0 | 0 | 1 | 0 | 0 | 0 |
| CCMB3125 | *hardwickei* | 1 | 4 | 6 | 0 | 0 | 1 | 0 | 1 | 0 | 0 | 0 | 1 | 0 | 0 | 1 | 1 | 0 | 1 | 4 | 1 | 0 | 0 | 1 | 0 | 0 | 0 |
| CCMB3279 | *hardwickei* | 1 | 5 | 2 | 0 | 0 | 0 | 0 | 0 | 0 | 0 | 0 | 0 | 0 | 0 | 1 | 1 | 0 | 2 | 3 | 0 | 0 | 0 | 0 | 0 | 0 | 0 |
| CCMB3280 | *hardwickei* | 1 | 5 | 0 | 0 | 0 | 0 | 0 | 2 | 0 | 0 | 0 | 3 | 0 | 1 | 1 | 1 | 0 | 1 | 5 | 0 | 0 | 0 | 1 | 0 | 0 | 0 |
| CCMB3281 | *hardwickei* | 1 | 6 | 2 | 1 | 0 | 0 | 0 | 0 | 1 | 0 | 0 | 3 | 0 | 1 | 1 | 1 | 0 | 2 | 3 | 0 | 0 | 0 | 1 | 0 | 0 | 0 |
| CCMB3338 | *hardwickei* | 2 | 4 | 2 | 0 | 0 | 0 | 0 | 0 | 0 | 0 | 0 | 3 | 0 | 0 | 1 | 1 | 0 | 1 | 5 | 0 | 0 | 0 | 1 | 1 | 0 | 0 |

**SI Table 2c: Summary Statistics of the venom proteins in *Scolopendra***

| Protein group Cumulative Copy Number | *S. morsitans* (n=28) | *S. hardwickei* (n=11) |
|---|---|---|
| Enzymes | 1167 | 556 |
| Scoloptoxins | 1280 | 272 |
| Beta pore forming toxins | 686 | 296 |
| Unchar proteins | 265 | 104 |
| LDLA domain containing proteins | 468 | 115 |
| DUF containing proteins | 376 | 42 |
| CAP proteins | 227 | 35 |
| Other proteins | 660 | 156 |
| Non-toxin components | 273 | 39 |

**For_Jaccard_Index**

| Genus | species | Acid-phosphatase | Alpha-2-macroglobulin | ALPHA-MANNOSIDASE | Apolipoprotein D | ARYLSULFATASE_ | BPFTx | C-type_Lectin | CAA-like | Calycin-Lipocalin | CAP | Carbon-nitrogen_hydrolase | Carboxylic ester hydrolase | Cathepsin | Chitinase | Chondroitinase | COEsteraseB | CUB-domai_protein | Cubilin_ | Cyclase-like_protein | Cystatin | Cysteine-rich venom protein | DUF1397 | DUF3472 | Fibronectin_or_immunoglobulin-like | GDH | GGT | GH18 | Haem-binding_protein | Icarapin-like |
|---|---|---|---|---|---|---|---|---|---|---|---|---|---|---|---|---|---|---|---|---|---|---|---|---|---|---|---|---|---|---|
| *Scolopendra* | *morsitans* | 1 | 1 | 1 | 1 | 1 | 1 | 1 | 1 | 1 | 1 | 1 | 0 | 1 | 1 | 1 | 1 | 1 | 1 | 1 | 1 | 1 | 1 | 1 | 1 | 1 | 1 | 1 | 1 | 1 |
| *Scolopendra* | *hardwickei* | 0 | 1 | 1 | 1 | 1 | 1 | 1 | 0 | 1 | 1 | 1 | 1 | 1 | 1 | 1 | 1 | 1 | 1 | 0 | 1 | 1 | 1 | 1 | 0 | 1 | 1 | 1 | 1 | 1 |
| | shared | 0 | 1 | 1 | 1 | 1 | 1 | 1 | 0 | 1 | 1 | 1 | 0 | 1 | 1 | 1 | 1 | 1 | 1 | 0 | 1 | 1 | 1 | 1 | 0 | 1 | 1 | 1 | 1 | 1 |

**Continued**

| Genus | species | Ige-ESP-like | Kazal_domain | LDLA | Leucine-rich_repeat | Lipid-transport_protein | Lysosomal_lipase | Lysozyme C | M12A | MCSACoa | Neprilysin | NOT_TOXIN_Actin | NOT_TOXIN_Arginine_kinase_ | NOT_TOXIN_ATP_synthase | NOT_TOXIN_Calcium-transporting_ATPase | NOT_TOXIN_Histone_H2A | NOT_TOXIN_Histone_H2B | NOT_TOXIN_INSUFFICIENT_EVIDENCE_Alpha-L-Fucosidase | NOT_TOXIN_INSUFFICIENT_EVIDENCE_Avidin-related | NOT_TOXIN_INSUFFICIENT_EVIDENCE_CUB-tolloid | NOT_TOXIN_INSUFFICIENT_EVIDENCE_NOT_TOXIN_INSUFFICIENT_EVIDENCE_insect_cuticle_uncharacterized_protein_ | NOT_TOXIN_Myosin_heavy_chain_protein | NOT_TOXIN_Myosin_light_chain_protein | NOT_TOXIN_Myosin_tail_1_domain-containing_protein | NOT_TOXIN_NO_SIGNALP_Calponin | NOT_TOXIN_NO_SIGNALP_Pyruvate_kinase | NOT_TOXIN_structural_component | NOT_TOXIN_Tropomyosin_ | NOT_TOXIN_Troponin | NOT_TOXIN_tubulin_alpha_chain_ |
|---|---|---|---|---|---|---|---|---|---|---|---|---|---|---|---|---|---|---|---|---|---|---|---|---|---|---|---|---|---|---|
| *Scolopendra* | *morsitans* | 1 | 1 | 1 | 1 | 1 | 0 | 1 | 1 | 0 | 1 | 1 | 1 | 1 | 1 | 1 | 1 | 1 | 1 | 1 | 1 | 1 | 1 | 1 | 0 | 1 | 1 | 0 | 1 | 1 |
| *Scolopendra* | *hardwickei* | 1 | 0 | 1 | 0 | 1 | 1 | 1 | 0 | 1 | 1 | 1 | 1 | 0 | 0 | 1 | 1 | 1 | 0 | 1 | 0 | 1 | 1 | 0 | 1 | 0 | 0 | 1 | 0 | 1 |
| | shared | 1 | 0 | 1 | 0 | 1 | 0 | 1 | 0 | 0 | 1 | 1 | 1 | 0 | 0 | 1 | 1 | 1 | 0 | 1 | 0 | 1 | 1 | 0 | 0 | 0 | 0 | 0 | 0 | 1 |

**Continued**

| Genus | species | NOT_TOXIN_tubulin_beta_chain_ | NOT_TOXIN_Vitellogenin-6 | PAM-like | PEBP-like | Peptidase_M13 | Phosphodiesterase 2 | PLA2 | Purple acid phosphatase | S1 | S10 | S8 | Saposin-related | SCTX01 | SCTX02 | Serpine | SLPTX-SSD552 | SLPTX-SSD558 | SLPTX-SSD976 | SLPTX01 | SLPTX02 | SLPTX03 | SLPTX04 | SLPTX05 | SLPTX06 | SLPTX07 | SLPTX08 | SLPTX09 | SLPTX10 | SLPTX11 |
|---|---|---|---|---|---|---|---|---|---|---|---|---|---|---|---|---|---|---|---|---|---|---|---|---|---|---|---|---|---|---|
| *Scolopendra* | *morsitans* | 1 | 1 | 1 | 1 | 1 | 1 | 1 | 1 | 1 | 1 | 1 | 1 | 1 | 1 | 1 | 1 | 1 | 1 | 1 | 1 | 1 | 1 | 1 | 1 | 1 | 1 | 1 | 1 | 1 |
| *Scolopendra* | *hardwickei* | 1 | 1 | 1 | 1 | 1 | 0 | 1 | 0 | 1 | 1 | 1 | 0 | 0 | 1 | 1 | 0 | 0 | 1 | 1 | 0 | 1 | 1 | 1 | 0 | 1 | 1 | 0 | 1 | 1 |
| | shared | 1 | 1 | 1 | 1 | 1 | 0 | 1 | 0 | 1 | 1 | 1 | 0 | 0 | 1 | 1 | 0 | 0 | 1 | 1 | 0 | 1 | 1 | 1 | 0 | 1 | 1 | 0 | 1 | 1 |

**Continued**

| Genus | species | SLPTX12 | SLPTX13 | SLPTX14 | SLPTX15 | SLPTX16 | SLPTX17 | SLPTX19 | SLPTX20 | SLPTX21 | SLPTX29 | SLPTX30 | SLPTX47 | TGFbeta | Thioester-containing protein 4 | Transferrin | Triacylglycerol_lipase_family | Unchar03 | Unchar05 | Unchar06 | Unchar08 | Unchar10 | Unchar11 | Unchar13 | Unchar30 | Unchar31 | Unchar32 | Unchar33 | Vigilin | WAP |
|---|---|---|---|---|---|---|---|---|---|---|---|---|---|---|---|---|---|---|---|---|---|---|---|---|---|---|---|---|---|---|
| *Scolopendra* | *morsitans* | 1 | 1 | 1 | 1 | 1 | 1 | 1 | 1 | 1 | 1 | 1 | 1 | 1 | 1 | 1 | 1 | 1 | 1 | 1 | 1 | 1 | 1 | 0 | 1 | 1 | 1 | 1 | 1 | 1 |
| *Scolopendra* | *hardwickei* | 1 | 1 | 1 | 1 | 1 | 1 | 1 | 0 | 1 | 1 | 1 | 1 | 0 | 1 | 1 | 0 | 1 | 1 | 1 | 0 | 1 | 1 | 1 | 0 | 0 | 1 | 1 | 0 | 0 |
| | shared | 1 | 1 | 1 | 1 | 1 | 1 | 1 | 0 | 1 | 1 | 1 | 1 | 0 | 1 | 1 | 0 | 1 | 1 | 1 | 0 | 1 | 1 | 0 | 0 | 0 | 1 | 1 | 0 | 0 |

| | |
|---|---|
| Proteins in *Scolopendra morsitans* (A) = 110 | |
| Proteins in *Scolopendra hardwickei* (B) = 84 | 0.7052631579 |
| Shared proteins (A∩B) = 67 | |
| J=67/95=0.705 | |

**For general summary**

| ID | species | Total number of protein hits (medium and high) | number of hits (medium and high confidence) greater than or equal to 10kDa | number of hits (medium and high confidence) less than 10kDa | protein abundance of knowns | protein richness |
|---|---|---|---|---|---|---|
| CCMB1159 | *morsitans* | 185 | 163 | 22 | 174 | 51 |
| CCMB1209 | *morsitans* | 184 | 158 | 26 | 175 | 48 |
| CCMB1325 | *morsitans* | 196 | 165 | 31 | 196 | 45 |
| CCMB1693 | *morsitans* | 260 | 221 | 39 | 241 | 57 |
| CCMB1695 | *morsitans* | 176 | 151 | 25 | 164 | 42 |
| CCMB1696 | *morsitans* | 237 | 193 | 44 | 195 | 50 |
| CCMB1725 | *morsitans* | 284 | 234 | 50 | 223 | 63 |
| CCMB1766 | *morsitans* | 218 | 190 | 28 | 204 | 47 |
| CCMB1769 | *morsitans* | 223 | 181 | 42 | 217 | 53 |
| CCMB1803 | *morsitans* | 201 | 175 | 26 | 178 | 52 |
| CCMB2184 | *morsitans* | 222 | 185 | 39 | 186 | 52 |
| CCMB2198 | *morsitans* | 172 | 146 | 26 | 147 | 43 |
| CCMB2382 | *morsitans* | 228 | 173 | 55 | 197 | 51 |
| CCMB3265 | *morsitans* | 260 | 215 | 45 | 196 | 66 |
| CCMB3772 | *morsitans* | 273 | 220 | 53 | 223 | 62 |
| CCMB3868 | *morsitans* | 173 | 139 | 34 | 143 | 44 |
| CCMB41 | *morsitans* | 104 | 86 | 18 | 90 | 28 |
| CCMB4114 | *morsitans* | 183 | 152 | 31 | 149 | 41 |
| CCMB4216 | *morsitans* | 241 | 191 | 50 | 213 | 58 |
| CCMB4224 | *morsitans* | 215 | 182 | 33 | 188 | 51 |
| CCMB4407 | *morsitans* | 233 | 190 | 43 | 205 | 52 |
| CCMB4441 | *morsitans* | 154 | 130 | 24 | 147 | 40 |
| CCMB4454 | *morsitans* | 111 | 94 | 17 | 109 | 33 |
| CCMB752 | *morsitans* | 251 | 206 | 45 | 193 | 55 |
| CCMB803 | *morsitans* | 225 | 184 | 41 | 209 | 55 |
| CCMB805 | *morsitans* | 185 | 160 | 25 | 183 | 49 |
| CCMB818 | *morsitans* | 210 | 176 | 34 | 188 | 50 |
| CCMB845 | *morsitans* | 207 | 167 | 40 | 201 | 48 |
| CCMB1245 | *hardwickei* | 183 | 147 | 36 | 176 | 41 |
| CCMB1691 | *hardwickei* | 198 | 171 | 27 | 175 | 43 |
| CCMB1692 | *hardwickei* | 165 | 132 | 33 | 159 | 40 |
| CCMB1776 | *hardwickei* | 258 | 219 | 39 | 211 | 55 |
| CCMB2194 | *hardwickei* | 135 | 112 | 23 | 124 | 36 |
| CCMB2197 | *hardwickei* | 116 | 93 | 23 | 112 | 32 |
| CCMB3125 | *hardwickei* | 155 | 128 | 27 | 148 | 41 |
| CCMB3279 | *hardwickei* | 115 | 92 | 23 | 105 | 30 |
| CCMB3280 | *hardwickei* | 198 | 164 | 34 | 135 | 44 |
| CCMB3281 | *hardwickei* | 199 | 166 | 33 | 160 | 53 |
| CCMB3338 | *hardwickei* | 142 | 120 | 22 | 110 | 37 |

| | Average number of protein hits (medium and high) | Average number of hits (medium and high confidence) greater than or equal to 10kDa | Average number of hits (medium and high confidence) less than 10kDa | richness | abundance |
|---|---|---|---|---|---|
| morsitans | 207.53571 | 172.39285 | 35.214285 | 49.5 | 183.35714 |
| hardwickei | 169.45454 | 140.36363 | 29.090909 | 41.090909 | 146.81818 |

| | STDEV number of protein hits (medium and high) | STDEV number of hits (medium and high confidence) greater than or equal to 10kDa | STDEV number of hits (medium and high confidence) less than 10kDa | STDEV richness | STDEV abundance |
|---|---|---|---|---|---|
| morsitans | 43.122978 | 34.353650 | 10.702568 | 8.3776598 | 34.092699 |
| hardwickei | 43.033390 | 37.955955 | 6.0902306 | 7.7259891 | 33.265051 |

**Toxin proteins richness**

| Genus | species | Acid-phosphatase | Alpha-2-macroglobulin | ALPHA-MANNOSIDASE | Apolipoprotein D | ARYLSULFATASE_ | BPFTx | C-type_Lectin | CAA-like | Calycin-Lipocalin | CAP | Carbon-nitrogen_hydrolase | Carboxylic ester hydrolase | Cathepsin | Chitinase | Chondroitinase | COEsteraseB | CUB-domai_protein | Cubilin_ | Cyclase-like_protein | Cystatin | Cysteine-rich venom protein | DUF1397 | DUF3472 | Fibronectin_or_immunoglobulin-like | GDH | GGT | GH18 | Haem-binding_protein | Icarapin-like |
|---|---|---|---|---|---|---|---|---|---|---|---|---|---|---|---|---|---|---|---|---|---|---|---|---|---|---|---|---|---|---|
| *Scolopendra* | *morsitans* | 1 | 1 | 1 | 1 | 1 | 1 | 1 | 1 | 1 | 1 | 1 | 0 | 1 | 1 | 1 | 1 | 1 | 1 | 1 | 1 | 1 | 1 | 1 | 1 | 1 | 1 | 1 | 1 | 1 |
| *Scolopendra* | *hardwickei* | 0 | 1 | 1 | 1 | 1 | 1 | 1 | 0 | 1 | 1 | 1 | 1 | 1 | 1 | 1 | 1 | 1 | 1 | 0 | 1 | 1 | 1 | 1 | 0 | 1 | 1 | 1 | 1 | 1 |

**Continued**

| Genus | species | Ige-ESP-like | Kazal_domain | LDLA | Leucine-rich_repeat | Lipid-transport_protein | Lysosomal_lipase | Lysozyme C | M12A | MCSACoa | Neprilysin | PAM-like | PEBP-like | Peptidase_M13 | Phosphodiesterase 2 | PLA2 | Purple acid phosphatase | S1 | S10 | S8 | Saposin-related | SCTX01 | SCTX02 | Serpine | SLPTX-SSD552 | SLPTX-SSD558 | SLPTX-SSD976 | SLPTX01 | SLPTX02 | SLPTX03 |
|---|---|---|---|---|---|---|---|---|---|---|---|---|---|---|---|---|---|---|---|---|---|---|---|---|---|---|---|---|---|---|
| *Scolopendra* | *morsitans* | 1 | 1 | 1 | 1 | 1 | 0 | 1 | 1 | 0 | 1 | 1 | 1 | 1 | 1 | 1 | 1 | 1 | 1 | 1 | 1 | 1 | 1 | 1 | 1 | 1 | 1 | 1 | 1 | 1 |
| *Scolopendra* | *hardwickei* | 1 | 0 | 1 | 0 | 1 | 1 | 1 | 0 | 1 | 1 | 1 | 1 | 1 | 0 | 1 | 0 | 1 | 1 | 1 | 0 | 0 | 1 | 1 | 0 | 0 | 1 | 1 | 0 | 1 |

**Continued**

| Genus | species | SLPTX04 | SLPTX05 | SLPTX06 | SLPTX07 | SLPTX08 | SLPTX09 | SLPTX10 | SLPTX11 | SLPTX12 | SLPTX13 | SLPTX14 | SLPTX15 | SLPTX16 | SLPTX17 | SLPTX19 | SLPTX20 | SLPTX21 | SLPTX29 | SLPTX30 | SLPTX47 | TGFbeta | Thioester-containing protein 4 | Transferrin | Triacylglycerol_lipase_family | Unchar03 | Unchar05 | Unchar06 | Unchar08 | Unchar10 |
|---|---|---|---|---|---|---|---|---|---|---|---|---|---|---|---|---|---|---|---|---|---|---|---|---|---|---|---|---|---|---|
| *Scolopendra* | *morsitans* | 1 | 1 | 1 | 1 | 1 | 1 | 1 | 1 | 1 | 1 | 1 | 1 | 1 | 1 | 1 | 1 | 1 | 1 | 1 | 1 | 1 | 1 | 1 | 1 | 1 | 1 | 1 | 1 | 1 |
| *Scolopendra* | *hardwickei* | 1 | 1 | 0 | 1 | 1 | 0 | 1 | 1 | 1 | 1 | 1 | 1 | 1 | 1 | 1 | 0 | 1 | 1 | 1 | 1 | 0 | 1 | 1 | 0 | 1 | 1 | 1 | 0 | 1 |

**Continued**

| Genus | species | Unchar11 | Unchar13 | Unchar30 | Unchar31 | Unchar32 | Unchar33 | Vigilin | WAP |
|---|---|---|---|---|---|---|---|---|---|
| *Scolopendra* | *morsitans* | 1 | 0 | 1 | 1 | 1 | 1 | 1 | 1 |
| *Scolopendra* | *hardwickei* | 1 | 1 | 0 | 0 | 1 | 1 | 0 | 0 |

**SI Table 2.d: Summary data**

| | *S. morsitans* (n=28) | *S. hardwickei* (n=11) |
|---|---|---|
| Number of individuals investigated | 28 | 11 |
| Total proteins identified (includes toxins and non toxins) | 110 | 84 |
| Total toxin proteins identified | 91 | 71 |
| Total non toxin proteins identified | 19 | 13 |
| Total unique toxin proteins identified | 24 | 4 |
| Average proteins richness | 49±8.37 | 41±7.72 |
| Average proteins abundance | 183.35±34.09 | 146.81±33.26 |

**SI Table 2e: Unique Proteins found in *Scolopendra***

| *Scolopendra hardwickei* | | | *Scolopendra morsitans* | | |
|---|---|---|---|---|---|
| Toxin proteins (protein group) | Found unique in: Across individuals | Possible roles in venom | Toxin proteins (protein group) | Found unique in: Across individuals | Possible roles in venom |

| | | | | | | |
|---|---|---|---|---|---|---|
| Carboxylic ester hydrolase (Enzyme) | 2/11 | A category of esterases, proposed to play a part in the release of endogenous purines during envenomation, these act as “multitoxins” causing a multitude of pharmacological effects including immobilization through hypotension (Hakim et al., 2015; Undheim et al., 2015) | | Acid-phosphatase (Enzyme) | 1/28 | A category of non-specific esterases, likely facilitate the release of endogenous purines during envenomation, triggering multi toxic effects such as immobilization through hypotension ( Malta et al., 2008; Hakim et al., 2015) |
| Lysosomal lipase (Enzyme) | 1/11 | A category of lipase, unknown function | | M12A Metalloprotease (Enzyme) | 1/28 | A Zinc metalloendopeptidase, A potential spreading factor likely involved in skin damage, blister formation, edema, myonecrosis and inflammation, consistent with some of the centipede bite symptoms (Undheim et al., 2015; Hakim et al., 2015) |
| Unchar 13 (uncharacterised proteins) | 1/11 | Unknown | | Purple acid phosphatase (Enzyme) | 1/28 | A category of non-specific esterases (Scolopendra subspinipes mutilans) (Zhao et al., 2018) |
| MCSACoa (other toxin proteins) | 1/11 | Unknown | | Cyclase like protein (Enzyme) | 2/28 | A category of acyltransferases (most likely Glutaminyl cyclase), involved in post translational processing of numerous bioactive proteins (Pawlak and Kini, 2006; Wang et al., 2014) |
| | | | | Triacylglycerol lipase (Enzyme) | 16/28 | A category of Lipases, thought to be involved in the pre-digestion of prey Dresler et al., 2024) |
| | | | | Phosphodiesterase2 (Enzyme) | 26/28 | Enzymatically acts on the extracellular ATP, likely perturbing the related physiological responses (Pan et al., 2023) |
| | | | | SLPTX 02 (Scoloptoxins) | 5/28 | Defensin-like with 6 cysteines, unknown function (Undheim et al., 2015) |
| | | | | SLPTX 06 (Scoloptoxins) | 8/28 | Unknown function (Undheim et al., 2015) |
| | | | | SLPTX 09 (Scoloptoxins) | 7/28 | Unknown; putative synergistic mode of action for peptides encoded by multidomain transcripts (Undheim et al., 2015) |
| | | | | SLPTX 20 (Scoloptoxins) | 14/28 | Unknown function (Undheim et al., 2015) |
| | | | | Unchar 08 (Uncharacterised proteins) | 12/28 | Unknown function (Undheim et al., 2015) |
| | | | | Unchar 30 (Uncharacterised proteins) | 7/28 | Unknown function |
| | | | | Unchar 31 (Uncharacterised proteins) | 3/28 | Unknown function |
| | | | | Vigilin (other toxin proteins) | 1/28 | Unknown function, hypothesized to be involved in regulating mRNA stability and translation, might be involved in RNA-mediated gene silencing in snake venoms (Rokyta et al., 2012) |
| | | | | WAP proteins (other toxin proteins) | 5/28 | Whey acidic protein domain containing proteins, antibacterial function in snake venoms (Nair et al., 2007) and crustaceans (Smith et al., 2008) |
| | | | | TGF beta (other toxin proteins) | 14/28 | Transforming growth factor proteins, involved in mediating Developmental Signalling Systems (Chipman et al., 2014) |
| | | | | Leucine rich repeat (LRR) domain containing proteins (other toxin proteins) | 7/28 | Shows similarity to the immune active proteins of spider-related animals, hypothesized to possibly be involved in protection of the venom gland against microbial infections (Langenegger et al., 2019), predicted to be involved in protein-ligand interactions in spiders (Haney et al., 2018) and parasitoids (Alvarado et al., 2019) |
| | | | | Kazal domain containing proteins (other toxin proteins) | 1/28 | Inhibit the catalytic activity of proteolytic enzymes in snakes (Fernandez et al., 2016), heteropterans (Walker et al., 2018) and centipede (Luan et al., 2019) venoms. Thought to ensure the regulation of proteinase activity crucial for the survival of venomous animals, possible synergistic role in association with other venom components |
| | | | | Fibronectin or immunoglobulin- like (other toxin proteins) | 4/28 | A large glycoprotein, associated with a multitude of cellular processes (cell adhesion, migration, cytoskeletal organization, morphology, and phagocytosis), interact with other proteins in the extracellular matrix including heparin and cell membrane receptors (Ward, 2020) |
| | | | | Saposin related proteins (other toxin proteins) | 6/28 | Proteins having a saposin like domain, unknown function in centipede venoms, act as macro conotoxins, inhibitor of the sensory neuron calcium channel in marine cone snails (Hackney et al., 2023) |

**Venom protein matrix used to build the PCA plot**

| ID | species | Acid-phosphatase | Alpha-2-macroglobulin | ALPHA-MANNOSIDASE | Apolipoprotein D | ARYLSULFATASE_ | BPFTx | C-type_Lectin | CAA-like | Calycin-Lipocalin | CAP | Carbon-nitrogen_hydrolase | Carboxylic ester hydrolase | Cathepsin | Chitinase | Chondroitinase | COEsterase B | CUB-domai_protein | Cubilin_ | Cyclase-like_protein | Cystatin | Cysteine-rich venom protein | DUF1397 | DUF3472 | Fibronectin_or_immunoglobulin-like | GDH | GGT | GH18 |
|---|---|---|---|---|---|---|---|---|---|---|---|---|---|---|---|---|---|---|---|---|---|---|---|---|---|---|---|---|
| CCMB1159 | *morsitans* | 0 | 2 | 1 | 0 | 0 | 18 | 1 | 0 | 0 | 11 | 0 | 0 | 1 | 1 | 1 | 1 | 13 | 1 | 0 | 1 | 0 | 0 | 12 | 0 | 3 | 1 | 0 |
| CCMB1209 | *morsitans* | 0 | 0 | 2 | 0 | 0 | 26 | 1 | 0 | 0 | 8 | 0 | 0 | 1 | 2 | 1 | 5 | 16 | 3 | 0 | 1 | 0 | 0 | 12 | 0 | 3 | 1 | 0 |
| CCMB1325 | *morsitans* | 0 | 1 | 2 | 0 | 0 | 31 | 2 | 0 | 0 | 11 | 0 | 0 | 0 | 1 | 1 | 1 | 16 | 3 | 0 | 1 | 0 | 0 | 8 | 0 | 3 | 0 | 0 |
| CCMB1693 | *morsitans* | 0 | 3 | 2 | 0 | 0 | 31 | 3 | 0 | 0 | 13 | 1 | 0 | 0 | 2 | 2 | 6 | 14 | 3 | 0 | 4 | 0 | 0 | 8 | 0 | 2 | 1 | 0 |
| CCMB1695 | *morsitans* | 0 | 0 | 0 | 0 | 0 | 27 | 0 | 0 | 0 | 12 | 0 | 0 | 0 | 1 | 1 | 4 | 12 | 2 | 0 | 2 | 0 | 0 | 4 | 0 | 2 | 1 | 0 |
| CCMB1696 | *morsitans* | 1 | 0 | 1 | 0 | 0 | 32 | 2 | 0 | 0 | 8 | 0 | 0 | 0 | 1 | 2 | 5 | 15 | 3 | 0 | 2 | 0 | 0 | 7 | 0 | 2 | 0 | 0 |
| CCMB1725 | *morsitans* | 0 | 12 | 0 | 1 | 0 | 21 | 1 | 0 | 2 | 8 | 7 | 0 | 0 | 3 | 1 | 2 | 11 | 3 | 0 | 2 | 0 | 1 | 2 | 0 | 1 | 0 | 4 |
| CCMB1766 | *morsitans* | 0 | 0 | 1 | 0 | 0 | 32 | 2 | 0 | 0 | 14 | 0 | 0 | 0 | 2 | 2 | 4 | 19 | 3 | 0 | 2 | 1 | 0 | 8 | 1 | 2 | 0 | 0 |
| CCMB1769 | *morsitans* | 0 | 1 | 2 | 0 | 0 | 33 | 2 | 0 | 0 | 11 | 0 | 0 | 0 | 2 | 2 | 5 | 16 | 3 | 0 | 2 | 0 | 0 | 10 | 0 | 1 | 0 | 0 |
| CCMB1803 | *morsitans* | 0 | 2 | 2 | 0 | 0 | 27 | 1 | 0 | 0 | 8 | 0 | 0 | 1 | 1 | 1 | 2 | 16 | 3 | 0 | 2 | 0 | 0 | 9 | 0 | 2 | 3 | 0 |
| CCMB2184 | *morsitans* | 0 | 1 | 1 | 0 | 0 | 26 | 2 | 0 | 0 | 8 | 0 | 0 | 0 | 2 | 1 | 2 | 17 | 4 | 0 | 2 | 0 | 0 | 11 | 0 | 1 | 0 | 0 |
| CCMB2198 | *morsitans* | 0 | 0 | 1 | 0 | 0 | 26 | 1 | 0 | 0 | 7 | 0 | 0 | 1 | 2 | 1 | 2 | 15 | 3 | 0 | 2 | 0 | 0 | 9 | 0 | 2 | 0 | 0 |
| CCMB2382 | *morsitans* | 0 | 6 | 0 | 0 | 0 | 20 | 2 | 0 | 1 | 11 | 2 | 0 | 0 | 1 | 1 | 3 | 11 | 6 | 0 | 3 | 1 | 0 | 3 | 0 | 1 | 0 | 0 |
| CCMB3265 | *morsitans* | 0 | 13 | 1 | 0 | 0 | 17 | 3 | 0 | 2 | 6 | 4 | 0 | 0 | 2 | 1 | 1 | 9 | 2 | 0 | 1 | 1 | 1 | 4 | 0 | 1 | 2 | 1 |
| CCMB3772 | *morsitans* | 0 | 13 | 1 | 0 | 1 | 19 | 2 | 0 | 2 | 10 | 0 | 0 | 1 | 2 | 1 | 4 | 12 | 3 | 1 | 2 | 1 | 0 | 9 | 0 | 2 | 0 | 0 |
| CCMB3868 | *morsitans* | 0 | 0 | 1 | 0 | 0 | 19 | 0 | 0 | 0 | 6 | 0 | 0 | 0 | 1 | 1 | 0 | 12 | 3 | 0 | 3 | 0 | 0 | 7 | 0 | 2 | 0 | 1 |
| CCMB41 | *morsitans* | 0 | 5 | 0 | 0 | 0 | 5 | 0 | 0 | 0 | 7 | 1 | 0 | 0 | 3 | 0 | 0 | 5 | 3 | 0 | 1 | 0 | 0 | 2 | 0 | 0 | 0 | 0 |
| CCMB4114 | *morsitans* | 0 | 0 | 0 | 0 | 0 | 20 | 0 | 2 | 0 | 5 | 0 | 0 | 0 | 1 | 1 | 2 | 14 | 1 | 0 | 0 | 0 | 0 | 7 | 1 | 2 | 0 | 0 |
| CCMB4216 | *morsitans* | 0 | 8 | 1 | 1 | 1 | 22 | 1 | 0 | 1 | 6 | 1 | 0 | 0 | 2 | 1 | 4 | 13 | 4 | 1 | 2 | 1 | 0 | 8 | 17 | 2 | 0 | 0 |
| CCMB4224 | *morsitans* | 0 | 3 | 1 | 0 | 1 | 25 | 0 | 0 | 1 | 6 | 0 | 0 | 1 | 1 | 1 | 2 | 13 | 3 | 0 | 3 | 2 | 0 | 7 | 0 | 2 | 0 | 0 |
| CCMB4407 | *morsitans* | 0 | 8 | 1 | 1 | 1 | 22 | 1 | 0 | 2 | 4 | 1 | 0 | 0 | 2 | 1 | 4 | 12 | 3 | 0 | 2 | 0 | 0 | 5 | 17 | 3 | 0 | 2 |
| CCMB4441 | *morsitans* | 0 | 0 | 1 | 0 | 0 | 23 | 0 | 0 | 0 | 3 | 0 | 0 | 0 | 1 | 1 | 3 | 13 | 4 | 0 | 1 | 0 | 0 | 4 | 0 | 3 | 0 | 0 |
| CCMB4454 | *morsitans* | 0 | 0 | 1 | 0 | 0 | 20 | 0 | 0 | 0 | 3 | 0 | 0 | 0 | 1 | 1 | 2 | 11 | 4 | 0 | 1 | 0 | 0 | 7 | 0 | 2 | 0 | 0 |
| CCMB752 | *morsitans* | 0 | 4 | 1 | 0 | 0 | 22 | 2 | 0 | 0 | 8 | 0 | 0 | 1 | 1 | 1 | 3 | 13 | 3 | 0 | 1 | 1 | 0 | 11 | 0 | 2 | 1 | 0 |
| CCMB803 | *morsitans* | 0 | 2 | 1 | 0 | 0 | 26 | 1 | 0 | 0 | 7 | 1 | 0 | 1 | 1 | 1 | 3 | 13 | 3 | 0 | 0 | 1 | 0 | 3 | 0 | 3 | 1 | 0 |
| CCMB805 | *morsitans* | 0 | 0 | 1 | 0 | 0 | 32 | 2 | 0 | 0 | 5 | 0 | 0 | 1 | 1 | 2 | 3 | 16 | 5 | 0 | 2 | 1 | 0 | 7 | 0 | 2 | 0 | 1 |
| CCMB818 | *morsitans* | 0 | 1 | 1 | 0 | 0 | 29 | 1 | 0 | 0 | 8 | 2 | 0 | 0 | 1 | 1 | 3 | 12 | 5 | 0 | 1 | 0 | 0 | 7 | 0 | 2 | 0 | 0 |
| CCMB845 | *morsitans* | 0 | 0 | 1 | 0 | 1 | 35 | 2 | 0 | 0 | 13 | 0 | 0 | 1 | 1 | 2 | 3 | 18 | 5 | 0 | 1 | 0 | 0 | 7 | 0 | 3 | 0 | 0 |
| CCMB1245 | *hardwickei* | 0 | 0 | 2 | 0 | 0 | 39 | 1 | 0 | 0 | 4 | 0 | 1 | 1 | 1 | 1 | 2 | 28 | 1 | 0 | 0 | 0 | 0 | 4 | 0 | 0 | 14 | 0 |
| CCMB1691 | *hardwickei* | 0 | 0 | 2 | 0 | 0 | 42 | 1 | 0 | 0 | 3 | 0 | 0 | 1 | 1 | 1 | 2 | 23 | 1 | 0 | 0 | 0 | 1 | 3 | 0 | 1 | 14 | 0 |
| CCMB1692 | *hardwickei* | 0 | 0 | 1 | 0 | 0 | 36 | 1 | 0 | 0 | 3 | 0 | 0 | 1 | 0 | 1 | 1 | 19 | 1 | 0 | 0 | 0 | 0 | 3 | 0 | 1 | 15 | 0 |
| CCMB1776 | *hardwickei* | 0 | 11 | 0 | 1 | 1 | 29 | 3 | 0 | 2 | 3 | 5 | 0 | 2 | 0 | 1 | 1 | 20 | 1 | 0 | 1 | 0 | 0 | 3 | 0 | 1 | 14 | 0 |
| CCMB2194 | *hardwickei* | 0 | 0 | 1 | 0 | 0 | 24 | 1 | 0 | 0 | 3 | 0 | 1 | 0 | 0 | 1 | 2 | 12 | 0 | 0 | 0 | 1 | 0 | 4 | 0 | 3 | 18 | 0 |
| CCMB2197 | *hardwickei* | 0 | 0 | 0 | 0 | 0 | 25 | 1 | 0 | 0 | 4 | 0 | 0 | 0 | 0 | 1 | 1 | 13 | 0 | 0 | 0 | 1 | 1 | 4 | 0 | 2 | 13 | 0 |
| CCMB3125 | *hardwickei* | 0 | 0 | 2 | 0 | 0 | 28 | 1 | 0 | 0 | 3 | 0 | 0 | 0 | 0 | 1 | 1 | 15 | 3 | 0 | 0 | 0 | 0 | 4 | 0 | 1 | 13 | 0 |
| CCMB3279 | *hardwickei* | 0 | 0 | 0 | 0 | 0 | 21 | 1 | 0 | 0 | 3 | 0 | 0 | 0 | 0 | 1 | 2 | 13 | 0 | 0 | 0 | 1 | 0 | 4 | 0 | 2 | 11 | 0 |
| CCMB3280 | *hardwickei* | 0 | 9 | 0 | 0 | 1 | 13 | 3 | 0 | 4 | 2 | 1 | 0 | 1 | 0 | 0 | 2 | 12 | 0 | 0 | 1 | 1 | 0 | 4 | 0 | 0 | 6 | 0 |
| CCMB3281 | *hardwickei* | 0 | 4 | 1 | 0 | 0 | 23 | 3 | 0 | 2 | 3 | 1 | 0 | 0 | 0 | 1 | 2 | 14 | 0 | 0 | 1 | 1 | 0 | 4 | 0 | 2 | 10 | 4 |
| CCMB3338 | *hardwickei* | 0 | 6 | 1 | 0 | 0 | 16 | 0 | 0 | 2 | 4 | 0 | 0 | 0 | 0 | 1 | 2 | 13 | 0 | 0 | 1 | 0 | 0 | 3 | 0 | 1 | 6 | 0 |

**Continued**

| ID | species | Haem-binding_protein | Icarapin-like | Ige-ESP-like | Kazal_domain | LDLA | Leucine-rich_repeat | Lipid-transport_protein | Lysosomal_lipase | LysozymeC | M12A | MCSACoa | Neprilysin | NOT_TOXIN_Actin | NOT_TOXIN_Arginine_kinase_ | NOT_TOXIN_ATP_synthase | NOT_TOXIN_Calcium-transporting_ATPase | NOT_TOXIN_Histone_H2A | NOT_TOXIN_Histone_H2B | NOT_TOXIN_INSUFFICIENT_EVIDENCE_Alpha-L-Fucosidase | NOT_TOXIN_INSUFFICIENT_EVIDENCE_Avidin-related | NOT_TOXIN_INSUFFICIENT_EVIDENCE_CUB-tolloid | NOT_TOXIN_INSUFFICIENT_EVIDENCE_NOT_TOXIN_INSUFFICIENT_EVIDENCE_insect_cuticle_uncharacterized_protein_ | NOT_TOXIN_Myosin_heavy_chain_protein | NOT_TOXIN_Myosin_light_chain_protein | NOT_TOXIN_Myosin_tail_1_domain-containing_protein | NOT_TOXIN_NO_SIGNALP_Calponin | NOT_TOXIN_NO_SIGNALP_Pyruvate_kinase |
|---|---|---|---|---|---|---|---|---|---|---|---|---|---|---|---|---|---|---|---|---|---|---|---|---|---|---|---|---|
| CCMB1159 | *morsitans* | 0 | 1 | 0 | 0 | 13 | 0 | 7 | 0 | 2 | 0 | 0 | 0 | 0 | 0 | 0 | 0 | 0 | 0 | 0 | 0 | 3 | 0 | 0 | 0 | 0 | 0 | 0 |
| CCMB1209 | *morsitans* | 0 | 0 | 0 | 0 | 11 | 0 | 1 | 0 | 2 | 0 | 0 | 0 | 0 | 0 | 0 | 1 | 0 | 0 | 0 | 0 | 4 | 0 | 0 | 0 | 0 | 0 | 0 |
| CCMB1325 | *morsitans* | 0 | 0 | 0 | 0 | 15 | 0 | 0 | 0 | 2 | 0 | 0 | 0 | 0 | 0 | 0 | 0 | 0 | 0 | 0 | 0 | 2 | 0 | 0 | 0 | 0 | 0 | 0 |
| CCMB1693 | *morsitans* | 1 | 1 | 0 | 0 | 19 | 0 | 9 | 0 | 3 | 0 | 0 | 0 | 0 | 0 | 0 | 0 | 0 | 1 | 0 | 1 | 6 | 0 | 0 | 0 | 0 | 0 | 0 |
| CCMB1695 | *morsitans* | 0 | 0 | 0 | 0 | 14 | 0 | 0 | 0 | 3 | 0 | 0 | 0 | 0 | 0 | 0 | 0 | 0 | 0 | 1 | 0 | 1 | 0 | 0 | 0 | 0 | 0 | 0 |
| CCMB1696 | *morsitans* | 0 | 1 | 1 | 0 | 17 | 0 | 0 | 0 | 2 | 0 | 0 | 0 | 0 | 0 | 0 | 0 | 0 | 0 | 0 | 0 | 2 | 1 | 0 | 0 | 0 | 0 | 0 |
| CCMB1725 | *morsitans* | 1 | 1 | 0 | 0 | 11 | 0 | 11 | 0 | 2 | 0 | 0 | 1 | 6 | 3 | 0 | 0 | 1 | 1 | 1 | 0 | 1 | 0 | 1 | 2 | 1 | 0 | 1 |
| CCMB1766 | *morsitans* | 0 | 1 | 1 | 0 | 16 | 0 | 0 | 0 | 4 | 0 | 0 | 0 | 0 | 0 | 0 | 0 | 0 | 0 | 0 | 0 | 5 | 0 | 0 | 0 | 0 | 0 | 0 |
| CCMB1769 | *morsitans* | 0 | 1 | 0 | 0 | 17 | 1 | 4 | 0 | 3 | 0 | 0 | 0 | 0 | 0 | 0 | 0 | 0 | 0 | 0 | 0 | 3 | 0 | 0 | 0 | 0 | 0 | 0 |
| CCMB1803 | *morsitans* | 0 | 0 | 0 | 0 | 13 | 0 | 8 | 0 | 2 | 0 | 0 | 0 | 1 | 0 | 0 | 0 | 0 | 0 | 0 | 0 | 1 | 0 | 0 | 0 | 0 | 0 | 0 |
| CCMB2184 | *morsitans* | 0 | 1 | 0 | 0 | 19 | 1 | 1 | 0 | 2 | 0 | 0 | 0 | 2 | 0 | 0 | 0 | 0 | 0 | 0 | 0 | 3 | 0 | 0 | 0 | 0 | 0 | 0 |
| CCMB2198 | *morsitans* | 0 | 1 | 0 | 0 | 14 | 0 | 0 | 0 | 2 | 0 | 0 | 0 | 0 | 0 | 0 | 0 | 0 | 0 | 0 | 0 | 4 | 0 | 0 | 0 | 0 | 0 | 0 |
| CCMB2382 | *morsitans* | 1 | 0 | 0 | 0 | 10 | 0 | 7 | 0 | 2 | 0 | 0 | 1 | 2 | 0 | 0 | 0 | 0 | 0 | 1 | 0 | 2 | 0 | 0 | 0 | 0 | 0 | 0 |
| CCMB3265 | *morsitans* | 1 | 0 | 0 | 1 | 10 | 0 | 7 | 0 | 2 | 0 | 0 | 1 | 3 | 0 | 1 | 0 | 1 | 1 | 1 | 0 | 2 | 0 | 1 | 1 | 0 | 0 | 0 |
| CCMB3772 | *morsitans* | 2 | 0 | 0 | 0 | 9 | 2 | 10 | 0 | 3 | 0 | 0 | 1 | 5 | 3 | 0 | 0 | 0 | 0 | 1 | 0 | 3 | 0 | 0 | 0 | 0 | 0 | 1 |
| CCMB3868 | *morsitans* | 1 | 0 | 1 | 0 | 13 | 0 | 1 | 0 | 3 | 0 | 0 | 0 | 0 | 0 | 0 | 0 | 0 | 0 | 2 | 0 | 1 | 0 | 0 | 0 | 0 | 0 | 0 |
| CCMB41 | *morsitans* | 0 | 0 | 0 | 0 | 7 | 0 | 11 | 0 | 0 | 0 | 0 | 0 | 1 | 0 | 0 | 0 | 1 | 1 | 0 | 0 | 1 | 0 | 0 | 0 | 0 | 0 | 0 |
| CCMB4114 | *morsitans* | 0 | 0 | 2 | 0 | 13 | 0 | 0 | 0 | 3 | 0 | 0 | 0 | 0 | 0 | 0 | 0 | 0 | 0 | 0 | 0 | 3 | 0 | 0 | 0 | 0 | 0 | 0 |
| CCMB4216 | *morsitans* | 1 | 0 | 1 | 0 | 12 | 1 | 9 | 0 | 3 | 0 | 0 | 0 | 3 | 0 | 0 | 0 | 1 | 0 | 4 | 0 | 2 | 0 | 0 | 0 | 0 | 0 | 0 |
| CCMB4224 | *morsitans* | 1 | 0 | 1 | 0 | 12 | 1 | 8 | 0 | 3 | 0 | 0 | 0 | 0 | 0 | 0 | 0 | 0 | 0 | 3 | 0 | 3 | 0 | 0 | 0 | 0 | 0 | 0 |
| CCMB4407 | *morsitans* | 1 | 0 | 0 | 0 | 13 | 0 | 10 | 0 | 2 | 0 | 0 | 0 | 3 | 0 | 0 | 0 | 0 | 0 | 2 | 0 | 2 | 0 | 0 | 0 | 0 | 0 | 0 |
| CCMB4441 | *morsitans* | 0 | 0 | 0 | 0 | 16 | 0 | 4 | 0 | 2 | 0 | 0 | 0 | 0 | 0 | 0 | 0 | 0 | 0 | 3 | 0 | 2 | 0 | 0 | 0 | 0 | 0 | 0 |
| CCMB4454 | *morsitans* | 0 | 0 | 0 | 0 | 10 | 0 | 1 | 0 | 3 | 0 | 0 | 0 | 0 | 0 | 0 | 0 | 0 | 0 | 0 | 0 | 2 | 0 | 0 | 0 | 0 | 0 | 0 |
| CCMB752 | *morsitans* | 0 | 1 | 0 | 0 | 10 | 2 | 9 | 0 | 2 | 0 | 0 | 1 | 1 | 0 | 0 | 0 | 1 | 1 | 0 | 0 | 4 | 0 | 2 | 0 | 0 | 0 | 0 |
| CCMB803 | *morsitans* | 1 | 1 | 1 | 0 | 18 | 0 | 8 | 0 | 2 | 0 | 0 | 0 | 0 | 0 | 0 | 0 | 0 | 0 | 0 | 0 | 3 | 0 | 0 | 0 | 0 | 0 | 0 |
| CCMB805 | *morsitans* | 0 | 1 | 0 | 0 | 14 | 1 | 0 | 0 | 2 | 0 | 0 | 0 | 0 | 0 | 0 | 0 | 0 | 0 | 0 | 0 | 2 | 0 | 0 | 0 | 0 | 0 | 0 |
| CCMB818 | *morsitans* | 0 | 1 | 0 | 0 | 14 | 0 | 0 | 0 | 2 | 1 | 0 | 0 | 0 | 0 | 0 | 0 | 0 | 0 | 0 | 0 | 2 | 0 | 0 | 0 | 0 | 0 | 0 |
| CCMB845 | *morsitans* | 0 | 1 | 0 | 0 | 16 | 0 | 0 | 0 | 2 | 0 | 0 | 0 | 0 | 0 | 0 | 0 | 0 | 0 | 0 | 0 | 1 | 0 | 0 | 0 | 0 | 0 | 0 |
| CCMB1245 | *hardwickei* | 0 | 1 | 0 | 0 | 15 | 0 | 0 | 0 | 3 | 0 | 1 | 0 | 0 | 0 | 0 | 0 | 0 | 1 | 0 | 0 | 2 | 0 | 0 | 0 | 0 | 0 | 0 |
| CCMB1691 | *hardwickei* | 0 | 1 | 0 | 0 | 15 | 0 | 0 | 0 | 3 | 0 | 0 | 0 | 0 | 0 | 0 | 0 | 0 | 0 | 0 | 0 | 0 | 0 | 0 | 0 | 0 | 0 | 0 |
| CCMB1692 | *hardwickei* | 0 | 0 | 0 | 0 | 13 | 0 | 0 | 0 | 3 | 0 | 0 | 0 | 0 | 0 | 0 | 0 | 0 | 0 | 0 | 0 | 0 | 0 | 0 | 0 | 0 | 0 | 0 |
| CCMB1776 | *hardwickei* | 1 | 1 | 1 | 0 | 16 | 0 | 7 | 1 | 3 | 0 | 0 | 1 | 0 | 0 | 0 | 0 | 0 | 0 | 1 | 0 | 0 | 0 | 1 | 0 | 0 | 0 | 0 |
| CCMB2194 | *hardwickei* | 0 | 0 | 1 | 0 | 9 | 0 | 0 | 0 | 0 | 0 | 0 | 0 | 0 | 0 | 0 | 0 | 0 | 0 | 0 | 0 | 0 | 0 | 0 | 0 | 0 | 0 | 0 |
| CCMB2197 | *hardwickei* | 0 | 0 | 0 | 0 | 7 | 0 | 0 | 0 | 0 | 0 | 0 | 0 | 0 | 1 | 0 | 0 | 0 | 0 | 0 | 0 | 0 | 0 | 0 | 0 | 0 | 0 | 0 |
| CCMB3125 | *hardwickei* | 0 | 0 | 0 | 0 | 11 | 0 | 0 | 0 | 3 | 0 | 0 | 0 | 0 | 0 | 0 | 0 | 0 | 0 | 0 | 0 | 0 | 0 | 0 | 0 | 0 | 0 | 0.00E+00 |
| CCMB3279 | *hardwickei* | 0 | 0 | 1 | 0 | 9 | 0 | 0 | 0 | 0 | 0 | 0 | 0 | 0 | 0 | 0 | 0 | 0 | 0 | 0 | 0 | 0 | 0 | 0 | 0 | 0 | 0 | 0 |
| CCMB3280 | *hardwickei* | 0 | 1 | 1 | 0 | 6 | 0 | 9 | 0 | 0 | 0 | 0 | 1 | 3 | 0 | 0 | 0 | 0 | 1 | 0 | 0 | 0 | 0 | 0 | 1 | 0 | 1 | 0 |
| CCMB3281 | *hardwickei* | 1 | 1 | 2 | 0 | 11 | 0 | 9 | 0 | 0 | 0 | 0 | 1 | 5 | 0 | 0 | 0 | 1 | 1 | 0 | 0 | 0 | 0 | 0 | 1 | 0 | 0 | 0 |
| CCMB3338 | *hardwickei* | 0 | 0 | 0 | 0 | 3 | 0 | 7 | 0 | 0 | 0 | 0 | 1 | 0 | 0 | 0 | 0 | 0 | 0 | 0 | 0 | 0 | 0 | 0 | 0 | 0 | 0 | 0 |

**Continued**

| ID | species | NOT_TOXIN_structural_component | NOT_TOXIN_Tropomyosin_ | NOT_TOXIN_Troponin | NOT_TOXIN_tubulin_alpha_chain_ | NOT_TOXIN_tubulin_beta_chain_ | NOT_TOXIN_Vitellogenin-6 | PAM-like | PEBP-like | Peptidase_M13 | Phosphodiesterase2 | PLA2 | Purple acid phosphatase | S1 | S10 | S8 | Saposin-related | SCTX01 | SCTX02 | Serpine | SLPTx-SSD552 | SLPTx-SSD558 | SLPTx-SSD976 | SLPTX01 | SLPTX02 | SLPTX03 | SLPTX04 | SLPTX05 |
|---|---|---|---|---|---|---|---|---|---|---|---|---|---|---|---|---|---|---|---|---|---|---|---|---|---|---|---|---|
| CCMB1159 | *morsitans* | 0 | 0 | 0 | 0 | 0 | 2 | 1 | 2 | 0 | 1 | 1 | 0 | 3 | 2 | 1 | 0 | 0 | 0 | 0 | 0 | 0 | 0 | 2 | 0 | 1 | 4 | 0 |
| CCMB1209 | *morsitans* | 0 | 0 | 0 | 0 | 0 | 2 | 2 | 2 | 0 | 1 | 4 | 0 | 8 | 1 | 2 | 0 | 0 | 0 | 1 | 0 | 0 | 0 | 2 | 0 | 1 | 3 | 0 |
| CCMB1325 | *morsitans* | 0 | 0 | 0 | 0 | 0 | 1 | 2 | 2 | 0 | 6 | 2 | 0 | 7 | 1 | 2 | 0 | 0 | 0 | 0 | 0 | 0 | 1 | 2 | 0 | 2 | 2.00E+00 | 1 |
| CCMB1693 | *morsitans* | 0 | 0 | 0 | 0 | 0 | 7 | 1 | 1 | 1 | 2 | 2 | 0 | 6 | 1 | 2 | 0 | 0 | 1 | 1 | 0 | 0 | 0 | 3 | 0 | 1 | 2 | 0 |
| CCMB1695 | *morsitans* | 0 | 0 | 0 | 0 | 0 | 0 | 1 | 0 | 0 | 1 | 3 | 0 | 7 | 1 | 1 | 0 | 0 | 0 | 0 | 0 | 0 | 0 | 2 | 0 | 1 | 4 | 0 |
| CCMB1696 | *morsitans* | 0 | 0 | 0 | 0 | 0 | 1 | 1 | 1 | 0 | 1 | 2 | 0 | 6 | 1 | 6 | 1 | 0 | 0 | 1 | 0 | 0 | 0 | 2 | 0 | 3 | 4 | 0 |
| CCMB1725 | *morsitans* | 0 | 0 | 0 | 1 | 0 | 8 | 0 | 1 | 0 | 1 | 2 | 0 | 9 | 0 | 0 | 1 | 0 | 0 | 3 | 0 | 0 | 0 | 1 | 0 | 1 | 2 | 0 |
| CCMB1766 | *morsitans* | 0 | 0 | 0 | 0 | 0 | 2 | 1 | 1 | 0 | 1 | 3 | 0 | 6 | 1 | 2 | 0 | 0 | 0 | 1 | 0 | 1 | 0 | 3 | 0 | 1 | 3 | 0 |
| CCMB1769 | *morsitans* | 0 | 0 | 0 | 1 | 0 | 3 | 1 | 1 | 0 | 1 | 2 | 0 | 8 | 1 | 1 | 1 | 0 | 0 | 0 | 0 | 0 | 0 | 2 | 0 | 2 | 4 | 0 |
| CCMB1803 | *morsitans* | 1 | 0 | 0 | 0 | 0 | 6 | 1 | 2 | 0 | 1 | 1 | 0 | 7 | 1 | 1 | 0 | 0 | 0 | 1 | 0 | 0 | 0 | 2 | 1 | 1 | 1 | 0 |
| CCMB2184 | *morsitans* | 0 | 0 | 0.00E+00 | 0 | 0 | 3 | 1 | 2 | 0 | 1 | 2 | 1 | 3 | 2 | 2 | 1 | 0 | 0 | 0 | 0 | 0 | 1 | 3 | 0 | 1 | 2 | 0 |
| CCMB2198 | *morsitans* | 0 | 0 | 0.00E+00 | 0 | 0 | 0 | 1 | 0 | 0 | 1 | 1 | 0 | 6 | 1 | 1 | 0 | 0 | 0 | 1 | 0 | 0 | 1 | 3 | 0 | 1 | 1 | 0 |
| CCMB2382 | *morsitans* | 0 | 0 | 0 | 0 | 0 | 6 | 1 | 0 | 1 | 1 | 2 | 0 | 12 | 2 | 1 | 0 | 0 | 0 | 3 | 0 | 0 | 0 | 2 | 0 | 3 | 4 | 0 |
| CCMB3265 | *morsitans* | 0 | 0 | 0 | 1 | 1 | 9 | 0 | 0 | 1 | 1 | 3 | 0 | 7 | 0 | 0 | 0 | 0 | 0 | 2 | 0 | 0 | 1 | 3 | 1 | 2 | 1 | 0 |
| CCMB3772 | *morsitans* | 0 | 0 | 0 | 0 | 0 | 8 | 1 | 2 | 3 | 2 | 1 | 0 | 11 | 1 | 2 | 0 | 0 | 0 | 3 | 0 | 1 | 0 | 2 | 1 | 1 | 1 | 0 |
| CCMB3868 | *morsitans* | 0 | 0 | 1 | 0 | 0 | 6 | 1 | 2 | 0 | 2 | 2 | 0 | 7 | 2 | 1 | 0 | 0 | 1 | 0 | 0 | 0 | 0 | 4 | 1 | 1 | 1 | 0 |
| CCMB41 | *morsitans* | 0 | 0 | 0 | 0 | 0 | 7 | 0 | 0 | 0 | 0 | 0 | 0 | 4 | 0 | 0 | 0 | 0 | 0 | 1 | 0 | 0 | 0 | 1 | 0 | 1 | 0 | 0 |
| CCMB4114 | *morsitans* | 0 | 0 | 0 | 0 | 0 | 0 | 1 | 3 | 0 | 2 | 2 | 0 | 6 | 2 | 2 | 1 | 0 | 0 | 1 | 0 | 0 | 0 | 2 | 0 | 3 | 2 | 0 |
| CCMB4216 | *morsitans* | 0 | 0 | 0 | 0 | 0 | 9 | 1 | 2 | 1 | 2 | 2 | 0 | 10 | 1 | 1 | 0 | 0 | 0 | 3 | 0 | 0 | 0 | 2 | 0 | 1 | 1 | 1 |
| CCMB4224 | *morsitans* | 0 | 0 | 0 | 0 | 0 | 8 | 1 | 2 | 2 | 2 | 2 | 0 | 9 | 1 | 1 | 0 | 0 | 0 | 4 | 0 | 0 | 0 | 2 | 0 | 1 | 1 | 0 |
| CCMB4407 | *morsitans* | 0 | 0 | 0 | 0 | 0 | 9 | 1 | 2 | 2 | 2 | 2 | 0 | 9 | 1 | 1 | 0 | 0 | 0 | 1 | 0 | 0 | 0 | 2 | 1 | 1 | 0 | 0 |
| CCMB4441 | *morsitans* | 0 | 0 | 0 | 0 | 0 | 5 | 1 | 2 | 0 | 2 | 2 | 0 | 7 | 1 | 1 | 0 | 0 | 0 | 0 | 0 | 0 | 0 | 3 | 0 | 1 | 0 | 1 |
| CCMB4454 | *morsitans* | 0 | 0 | 0 | 0 | 0 | 3 | 1 | 2 | 0 | 2 | 1 | 0 | 5 | 0 | 0 | 0 | 0 | 0 | 0 | 0 | 0 | 0 | 2 | 0 | 1 | 0 | 0 |
| CCMB752 | *morsitans* | 0 | 0 | 1 | 0 | 0 | 5 | 1 | 2 | 3 | 0 | 2 | 0 | 7 | 2 | 1 | 0 | 0 | 0 | 0 | 0 | 0 | 1 | 3 | 0 | 1 | 2 | 0 |
| CCMB803 | *morsitans* | 0 | 0 | 0 | 0 | 0 | 6 | 1 | 1 | 1 | 1 | 3 | 0 | 3 | 1 | 2 | 0 | 0 | 0 | 0 | 0 | 0 | 0 | 2 | 0 | 3 | 4 | 0 |
| CCMB805 | *morsitans* | 0 | 0 | 0 | 0 | 0 | 0 | 2 | 1 | 0 | 1 | 1 | 0 | 6 | 1 | 1 | 1 | 1 | 0 | 1 | 0 | 0 | 0 | 2 | 0 | 1 | 4 | 0 |
| CCMB818 | *morsitans* | 0 | 0 | 0 | 0 | 0 | 2 | 1 | 0 | 0 | 1 | 1 | 0 | 6 | 1 | 2 | 0 | 0 | 2 | 0 | 0 | 0 | 0 | 3 | 0 | 1 | 4 | 0 |
| CCMB845 | *morsitans* | 0 | 0 | 2 | 0 | 0 | 0 | 1 | 0 | 0 | 1 | 2 | 0 | 6 | 1 | 1 | 0 | 0 | 0 | 0 | 1 | 0 | 0 | 2 | 0 | 1 | 3 | 0 |
| CCMB1245 | *hardwickei* | 0 | 0 | 0 | 0 | 0 | 0 | 1 | 1 | 1 | 0 | 7 | 0 | 4 | 1 | 0 | 0 | 0 | 0 | 0 | 0 | 0 | 0 | 2 | 0 | 0 | 3 | 0 |
| CCMB1691 | *hardwickei* | 0 | 0 | 0 | 0 | 0 | 1 | 2 | 1 | 0 | 0 | 6 | 0 | 5 | 1 | 2 | 0 | 0 | 0 | 0 | 0 | 0 | 0 | 2 | 0 | 1 | 0 | 1 |
| CCMB1692 | *hardwickei* | 0 | 0 | 0 | 0 | 0 | 0 | 1 | 1 | 1 | 0 | 4 | 0 | 6 | 1 | 1 | 0 | 0 | 0 | 0 | 0 | 0 | 0 | 2 | 0 | 1 | 3 | 0 |
| CCMB1776 | *hardwickei* | 0 | 0 | 0 | 0 | 0 | 3 | 1 | 1 | 4 | 0 | 6 | 0 | 9 | 1 | 2 | 0 | 0 | 2 | 6 | 0 | 0 | 1 | 2 | 0 | 1 | 0 | 1 |
| CCMB2194 | *hardwickei* | 0 | 0 | 0 | 0 | 0 | 0 | 1 | 1 | 0 | 0 | 4 | 0 | 4 | 1 | 1 | 0 | 0 | 0 | 0 | 0 | 0 | 0 | 2 | 0 | 1 | 0 | 0 |
| CCMB2197 | *hardwickei* | 0 | 0 | 0 | 0 | 0 | 0 | 1 | 1 | 2 | 0 | 2 | 0 | 4 | 0 | 0 | 0 | 0 | 0 | 0 | 0 | 0 | 0 | 2 | 0 | 1 | 0 | 0 |
| CCMB3125 | *hardwickei* | 0 | 0 | 0 | 0 | 0 | 2 | 1 | 1 | 1 | 0 | 5 | 0 | 9 | 1 | 1 | 0 | 0 | 0 | 0 | 0 | 0 | 0 | 2 | 0 | 1 | 1 | 0 |
| CCMB3279 | *hardwickei* | 0 | 0 | 0 | 0 | 0 | 0 | 1 | 1 | 0 | 0 | 3 | 0 | 4 | 1 | 0 | 0 | 0 | 0 | 0 | 0 | 0 | 0 | 1 | 0 | 0 | 0 | 0 |
| CCMB3280 | *hardwickei* | 0 | 0 | 0 | 1 | 1 | 4 | 0 | 0 | 1 | 0 | 3 | 0 | 9 | 0 | 0 | 0 | 0 | 1 | 5 | 0 | 0 | 0 | 1 | 0 | 0 | 0 | 0 |
| CCMB3281 | *hardwickei* | 0 | 1 | 0 | 0 | 1 | 2 | 2 | 0 | 2 | 0 | 3 | 0 | 4 | 1 | 0 | 0 | 0 | 0 | 0 | 0 | 0 | 0 | 1 | 0 | 0 | 0 | 0 |
| CCMB3338 | *hardwickei* | 0 | 0 | 0 | 0 | 0 | 3 | 0 | 0 | 1 | 0 | 2 | 0 | 2 | 0 | 1 | 0 | 0 | 1 | 1 | 0 | 0 | 0 | 1 | 0 | 0 | 0 | 0 |

**Continued**

| ID | species | SLPTX06 | SLPTX07 | SLPTX08 | SLPTX09 | SLPTX10 | SLPTX11 | SLPTX12 | SLPTX13 | SLPTX14 | SLPTX15 | SLPTX16 | SLPTX17 | SLPTX19 | SLPTX20 | SLPTX21 | SLPTX29 | SLPTX30 | SLPTX47 | TGFbeta | Thioester-containing protein 4 | Transferrin | Triacylglycerol_lipase_family | Unchar03 | Unchar05 | Unchar06 | Unchar08 | Unchar10 | Unchar11 | Unchar13 | Unchar30 | Unchar31 | Unchar32 | Unchar33 | Vigilin | WAP |
|---|---|---|---|---|---|---|---|---|---|---|---|---|---|---|---|---|---|---|---|---|---|---|---|---|---|---|---|---|---|---|---|---|---|---|---|---|
| CCMB1159 | *morsitans* | 2 | 1 | 0 | 1 | 5 | 9 | 1 | 2 | 1 | 11 | 5 | 2 | 0 | 1 | 1 | 0 | 0 | 0 | 1 | 0 | 2 | 0 | 0 | 7 | 1 | 0 | 3 | 3 | 0 | 0 | 0 | 0 | 0 | 0 | 1 |
| CCMB1209 | *morsitans* | 0 | 1 | 0 | 1 | 3 | 9 | 1 | 2 | 0 | 9 | 4 | 1 | 1 | 2 | 0 | 0 | 0 | 0 | 1 | 0 | 0 | 1 | 0 | 5 | 1 | 1 | 2 | 0 | 0 | 0 | 0 | 0 | 0 | 0 | 0 |
| CCMB1325 | *morsitans* | 0 | 0 | 0 | 0 | 6 | 17 | 1 | 0 | 8 | 13 | 4 | 1 | 1 | 1 | 2 | 0 | 0 | 0 | 0 | 0 | 0 | 0 | 0 | 4 | 1 | 0 | 2 | 1 | 0 | 1 | 0 | 0 | 0 | 0 | 2 |
| CCMB1693 | *morsitans* | 2 | 0 | 0 | 0 | 4 | 11 | 1 | 3.00E+00 | 7 | 10 | 5 | 4 | 2 | 1 | 1 | 0 | 0 | 0 | 1 | 0 | 7 | 4 | 0 | 4 | 1 | 1 | 2 | 3 | 0 | 0 | 0 | 0 | 0 | 0 | 0 |
| CCMB1695 | *morsitans* | 0 | 1 | 1 | 0 | 4 | 10 | 1 | 3 | 0 | 11 | 6 | 0 | 1 | 1 | 0 | 0 | 1 | 0 | 1 | 0 | 4 | 3 | 0 | 3 | 1 | 0 | 2 | 2 | 0 | 0 | 0 | 0 | 0 | 0 | 0 |
| CCMB1696 | *morsitans* | 0 | 0 | 1 | 0 | 3 | 17 | 0 | 3 | 0 | 13 | 4 | 2 | 2 | 1 | 1 | 0 | 0 | 0 | 1 | 0 | 0 | 1 | 0 | 4 | 1 | 1 | 2 | 1 | 0 | 2 | 0 | 0 | 0 | 0 | 0 |
| CCMB1725 | *morsitans* | 0 | 1 | 0 | 0 | 4 | 13 | 0 | 3 | 0 | 14 | 2 | 2 | 1 | 1 | 0 | 0 | 4 | 0 | 1 | 0 | 9 | 0 | 0 | 1 | 1 | 0 | 2 | 3 | 0 | 1 | 0 | 2 | 1 | 0 | 0 |
| CCMB1766 | *morsitans* | 2 | 0 | 0 | 0 | 5 | 9 | 1 | 0 | 0 | 12 | 5 | 9 | 2 | 1 | 0 | 0 | 0 | 0 | 1 | 0 | 0 | 0 | 0 | 4 | 1 | 1 | 2 | 3 | 0 | 0 | 0 | 0 | 0 | 0 | 0 |
| CCMB1769 | *morsitans* | 2 | 1 | 0 | 0 | 4 | 15 | 1 | 3 | 7 | 15 | 4 | 2 | 1 | 1 | 1 | 0 | 0 | 0 | 1 | 0 | 1 | 0 | 0 | 5 | 1 | 1 | 2 | 1 | 0 | 1 | 0 | 0 | 0 | 0 | 0 |
| CCMB1803 | *morsitans* | 0 | 1 | 0 | 1 | 2 | 9 | 1 | 0 | 0 | 7 | 4 | 0 | 1 | 0 | 0 | 0 | 2 | 0 | 1 | 0 | 1 | 2 | 0 | 4 | 1 | 1 | 3 | 4 | 0 | 0 | 0 | 1 | 0 | 0 | 0 |
| CCMB2184 | *morsitans* | 2 | 1 | 0 | 0 | 2 | 11 | 1 | 0 | 0 | 13 | 5 | 4 | 1 | 0 | 1 | 0 | 1 | 0 | 0 | 0 | 1 | 2 | 0 | 2 | 2 | 1 | 3 | 1 | 0 | 0 | 0 | 0 | 0 | 0 | 0 |
| CCMB2198 | *morsitans* | 0 | 1 | 0 | 0 | 1 | 8 | 1 | 0 | 0 | 7 | 3 | 3 | 1 | 0 | 1 | 0 | 0 | 0 | 0 | 0 | 0 | 1 | 0 | 3 | 1 | 1 | 2 | 2 | 0 | 0 | 0 | 0 | 0 | 0 | 0 |
| CCMB2382 | *morsitans* | 0 | 1 | 0 | 0 | 4 | 15 | 0 | 3 | 9 | 8 | 3 | 0 | 1 | 0 | 0 | 0 | 0 | 0 | 1 | 0 | 7 | 0 | 0 | 2 | 1 | 0 | 2 | 2 | 0 | 1 | 0 | 1 | 0 | 0 | 0 |
| CCMB3265 | *morsitans* | 0 | 1 | 0 | 0 | 1 | 10 | 0 | 0 | 0 | 12 | 6 | 1 | 1 | 1 | 1 | 0 | 2 | 0 | 0 | 1 | 8 | 0 | 1 | 3 | 1 | 1 | 2 | 2 | 0 | 0 | 0 | 2 | 1 | 0 | 0 |
| CCMB3772 | *morsitans* | 0 | 1 | 0 | 0 | 5 | 10 | 1 | 0 | 0 | 8 | 5 | 1 | 1 | 0 | 1 | 0 | 2 | 1 | 0 | 0 | 8 | 0 | 0 | 2 | 1 | 1 | 3 | 4 | 0 | 0 | 0 | 2 | 0 | 0 | 0 |
| CCMB3868 | *morsitans* | 0 | 0 | 1 | 0 | 3 | 6 | 1 | 0 | 0 | 8 | 5 | 0 | 0 | 0 | 0 | 1 | 0 | 0 | 0 | 0 | 2 | 0 | 0 | 3 | 1 | 0 | 1 | 1 | 0 | 0 | 0 | 0 | 0 | 0 | 0 |
| CCMB41 | *morsitans* | 0 | 0 | 0 | 0 | 1 | 2 | 0 | 0 | 0 | 6 | 5 | 0 | 0 | 0 | 0 | 0 | 2 | 0 | 0 | 0 | 3 | 0 | 0 | 0 | 0 | 0 | 1 | 0 | 0 | 0 | 0 | 0 | 2 | 0 | 0 |
| CCMB4114 | *morsitans* | 0 | 0 | 0 | 0 | 1 | 16 | 0 | 0 | 0 | 11 | 3 | 1 | 1 | 0 | 0 | 0 | 1 | 0 | 0 | 0 | 0 | 1 | 0 | 3 | 1 | 0 | 1 | 2 | 0 | 1 | 2 | 0 | 0 | 0 | 0 |
| CCMB4216 | *morsitans* | 0 | 0 | 0 | 0 | 3 | 7 | 0 | 0 | 0 | 10 | 3 | 5 | 1 | 0 | 0 | 0 | 2 | 0 | 0 | 0 | 1 | 1 | 0 | 2 | 1 | 1 | 1 | 4 | 0 | 0 | 0 | 0 | 0 | 0 | 0 |
| CCMB4224 | *morsitans* | 0 | 0 | 0 | 0 | 3 | 8 | 0 | 0 | 0 | 12 | 4 | 7 | 1 | 0 | 0 | 0 | 3 | 0 | 0 | 0 | 1 | 2 | 0 | 3 | 1 | 1 | 1 | 1 | 0 | 0 | 0 | 0 | 0 | 0 | 0 |
| CCMB4407 | *morsitans* | 0 | 0 | 0 | 0 | 3 | 14 | 0 | 0 | 0 | 8 | 1 | 0 | 1 | 0 | 0 | 1 | 1 | 0 | 0 | 0 | 7 | 2 | 0 | 2 | 1 | 0 | 1 | 4 | 0 | 0 | 0 | 0 | 0 | 0 | 0 |
| CCMB4441 | *morsitans* | 0 | 0 | 0 | 0 | 2 | 14 | 0 | 0 | 0 | 8 | 2 | 0 | 1 | 1 | 0 | 0 | 0 | 0 | 0 | 0 | 1 | 2 | 0 | 2 | 1 | 0 | 1 | 1 | 0 | 0 | 1 | 0 | 0 | 0 | 0 |
| CCMB4454 | *morsitans* | 0 | 0 | 0 | 0 | 1 | 6 | 0 | 0 | 0 | 7 | 2 | 0 | 0 | 0 | 0 | 0 | 0 | 0 | 0 | 0 | 0 | 1 | 0 | 2 | 1 | 0 | 1 | 1 | 0 | 0 | 0 | 0 | 0 | 1 | 0 |
| CCMB752 | *morsitans* | 0 | 0 | 0 | 0 | 2 | 9 | 1 | 0 | 1 | 13 | 5 | 3 | 1 | 0 | 1 | 0 | 2 | 0 | 0 | 0 | 2 | 0 | 0 | 4 | 2 | 0 | 2 | 5 | 0 | 0 | 0 | 0 | 0 | 0 | 0 |
| CCMB803 | *morsitans* | 0 | 0 | 0 | 2 | 6 | 18 | 0 | 3 | 3 | 24 | 2 | 1 | 1 | 1 | 1 | 0 | 0 | 0 | 1 | 0 | 5 | 0 | 0 | 2 | 1 | 0 | 2 | 2 | 0 | 1 | 2 | 0 | 0 | 0 | 1 |
| CCMB805 | *morsitans* | 2 | 1 | 0 | 2 | 6 | 12 | 1 | 0 | 1 | 15 | 6 | 0 | 0 | 1 | 2 | 0 | 0 | 0 | 1 | 0 | 0 | 1 | 0 | 4 | 0 | 0 | 2 | 2 | 0 | 0 | 0 | 0 | 0 | 0 | 0 |
| CCMB818 | *morsitans* | 2 | 1 | 0 | 2 | 5 | 12 | 1 | 3 | 0 | 14 | 6 | 0 | 1 | 0 | 2 | 0 | 3 | 0 | 1 | 0 | 5 | 1 | 0 | 6 | 1 | 0 | 1 | 2 | 0 | 0 | 0 | 0 | 0 | 0 | 1 |
| CCMB845 | *morsitans* | 2 | 1 | 0 | 2 | 7 | 10 | 1 | 3 | 1 | 15 | 9 | 0 | 1 | 1 | 2 | 0 | 0 | 0 | 1 | 0 | 0 | 2 | 0 | 5 | 0 | 0 | 1 | 2 | 0 | 0 | 0 | 0 | 0 | 0 | 1 |
| CCMB1245 | *hardwickei* | 0 | 0 | 0 | 0 | 4 | 3 | 1 | 1 | 2 | 1 | 5 | 4 | 0 | 0 | 1 | 0 | 1 | 0 | 0 | 0 | 0 | 0 | 0 | 3 | 1 | 0 | 5 | 2 | 0 | 0 | 0 | 0 | 0 | 0 | 0 |
| CCMB1691 | *hardwickei* | 0 | 0 | 1 | 0 | 6 | 3 | 1 | 1 | 1 | 2 | 5 | 4 | 0 | 0 | 1 | 0 | 1 | 0 | 0 | 0 | 0 | 0 | 0 | 3 | 2 | 0 | 3 | 3 | 0 | 0 | 0 | 1 | 0 | 0 | 0 |
| CCMB1692 | *hardwickei* | 0 | 0 | 0 | 0 | 6 | 3 | 1 | 1 | 1 | 1 | 5 | 5 | 0 | 0 | 1 | 2 | 1 | 0 | 0 | 0 | 0 | 0 | 0 | 1 | 1 | 0 | 5 | 4 | 0 | 0 | 0 | 1 | 0 | 0 | 0 |
| CCMB1776 | *hardwickei* | 0 | 0 | 0 | 0 | 4 | 3 | 0 | 1 | 0 | 1 | 8 | 6 | 0 | 0 | 0 | 0 | 5 | 0 | 0 | 1 | 2 | 0 | 0 | 1 | 1 | 0 | 2 | 3 | 0 | 0 | 0 | 1 | 0 | 0 | 0 |
| CCMB2194 | *hardwickei* | 0 | 0 | 1 | 0 | 3 | 4 | 1 | 0 | 1 | 1 | 6 | 2 | 0 | 0 | 0 | 0 | 1 | 0 | 0 | 0 | 0 | 0 | 0 | 1 | 1 | 0 | 3 | 2 | 0 | 0 | 0 | 1 | 0 | 0 | 0 |
| CCMB2197 | *hardwickei* | 0 | 0 | 0 | 0 | 3 | 4 | 1 | 0 | 1 | 1 | 5 | 2 | 0 | 0 | 0 | 0 | 0 | 0 | 0 | 0 | 0 | 0 | 0 | 1 | 1 | 0 | 3 | 2 | 0 | 0 | 0 | 1 | 0 | 0 | 0 |
| CCMB3125 | *hardwickei* | 0 | 0 | 0 | 0 | 8 | 3 | 1 | 1 | 1 | 1 | 4 | 6 | 0 | 0 | 1 | 0 | 1 | 0 | 0 | 0 | 1 | 0 | 0 | 1 | 1 | 0 | 1 | 4 | 1 | 0 | 0 | 1 | 0 | 0 | 0 |
| CCMB3279 | *hardwickei* | 0 | 0 | 1 | 0 | 3 | 4 | 1 | 0 | 1 | 1 | 5 | 2 | 0 | 0 | 0 | 0 | 0 | 0 | 0 | 0 | 0 | 0 | 0 | 1 | 1 | 0 | 2 | 3 | 0 | 0 | 0 | 0 | 0 | 0 | 0 |
| CCMB3280 | *hardwickei* | 0 | 0 | 0 | 0 | 0 | 3 | 0 | 0 | 1 | 1 | 5 | 0 | 0 | 0 | 0 | 0 | 2 | 0 | 0 | 0 | 3 | 0 | 1 | 1 | 1 | 0 | 1 | 5 | 0 | 0 | 0 | 1 | 0 | 0 | 0 |
| CCMB3281 | *hardwickei* | 0 | 1 | 1 | 0 | 3 | 4 | 1 | 0 | 1 | 1 | 6 | 2 | 1 | 0 | 0 | 0 | 0 | 1 | 0 | 0 | 3 | 0 | 1 | 1 | 1 | 0 | 2 | 3 | 0 | 0 | 0 | 1 | 0 | 0 | 0 |
| CCMB3338 | *hardwickei* | 0 | 0 | 1 | 0 | 4 | 4 | 0 | 0 | 1 | 2 | 4 | 2 | 0 | 0 | 0 | 0 | 0 | 0 | 0 | 0 | 3 | 0 | 0 | 1 | 1 | 0 | 1 | 5 | 0 | 0 | 0 | 1 | 1 | 0 | 0 |

**Loadings**

| | PC 1 | PC 2 | PC 3 |
|---|---|---|---|
| SLPTX15 | 0.23563 | 0.47879 | -0.01874 |
| SLPTX11 | 0.17855 | 0.38616 | 0.023236 |
| BPFTx | -0.56536 | 0.38325 | 0.38671 |
| LDLA | -0.10377 | 0.29234 | 0.14465 |
| CAP | 0.084873 | 0.24504 | 0.01125 |
| DUF3472 | 0.030778 | 0.15935 | -0.1023 |
| Unchar05 | -0.0080535 | 0.12164 | -0.03533 |
| SLPTX04 | -0.0091008 | 0.11956 | -0.0087996 |
| Cubilin_ | 0.051912 | 0.10501 | 0.0097487 |
| NOT_TOXIN_INSUFFICIENT_EVIDENCE_CUB-tolloid | 0.041743 | 0.10362 | 0.0029458 |
| CUB-domai_protein | -0.31367 | 0.092819 | 0.21659 |
| COEsteraseB | 0.013199 | 0.080413 | 0.062156 |
| SLPTX14 | -0.0044898 | 0.074805 | 0.057616 |
| SLPTX13 | -0.0029154 | 0.067032 | 0.040021 |
| LysozymeC | -0.016794 | 0.063264 | 0.067446 |
| SLPTX10 | -0.077768 | 0.06304 | 0.088256 |
| S8 | -0.013185 | 0.060275 | 0.011271 |
| Phosphodiesterase2 | 0.03244 | 0.058866 | -0.0035283 |
| SLPTX06 | -0.0059719 | 0.047322 | -0.0092863 |
| Triacylglycerol_lipase_family | 0.0092413 | 0.043858 | 0.0023724 |
| SLPTX03 | 0.017689 | 0.043614 | -0.0041472 |
| SLPTX19 | 0.017831 | 0.042027 | 0.023971 |

| | | | |
|---|---|---|---|
| Cystatin | 0.045083 | 0.039113 | 0.038721 |
| SLPTX21 | -0.016662 | 0.038978 | 0.0044082 |
| GDH | 0.0062864 | 0.038697 | -0.033619 |
| Chitinase | 0.042926 | 0.034959 | 0.024431 |
| SLPTX20 | 0.0049229 | 0.03371 | -0.004316 |
| TGFbeta | 0.0044858 | 0.032675 | 0.0017007 |
| SLPTX01 | -0.0029222 | 0.032665 | -0.0080529 |
| Chondroitinase | -0.012233 | 0.031424 | 0.0094687 |
| SLPTX09 | -0.00063365 | 0.030144 | -0.015066 |
| ALPHA-MANNOSIDASE | -0.024458 | 0.025684 | 0.018591 |
| S10 | -0.0059758 | 0.025302 | -0.0072004 |
| Unchar30 | 0.0067782 | 0.021494 | 0.001593 |
| PAM-like | -0.024649 | 0.020063 | -0.0024969 |
| PEBP-like | 0.0059459 | 0.019347 | -0.0047191 |
| WAP | 0.00046848 | 0.019188 | -0.0092892 |
| Icarapin-like | -0.010259 | 0.019044 | 0.021418 |
| Unchar08 | 0.0076047 | 0.016974 | 0.015304 |
| S1 | 0.075811 | 0.016381 | 0.20703 |
| SLPTX07 | 0.010477 | 0.015978 | 0.00019076 |
| Saposin-related | 0.0022332 | 0.015009 | -0.003581 |
| Unchar31 | 0.0063003 | 0.011135 | -0.013508 |
| SLPTX12 | -0.026713 | 0.010356 | -0.015261 |
| C-type_Lectin | -0.0021989 | 0.0094938 | 0.07949 |
| Leucine-rich_repeat | 0.012987 | 0.0090881 | 0.017937 |
| NOT_TOXIN_Troponin | -0.00072115 | 0.0080599 | -0.007208 |
| SLPTx-SSD552 | -0.0017839 | 0.0040481 | -0.00076323 |
| Acid-phosphatase | -0.00086567 | 0.0038713 | -0.0015832 |
| NOT_TOXIN_INSUFFICIENT_EVIDENCE_NOT_TOXIN_INSUFFICIENT_EVIDENCE_insect_cuticle_uncharacterized_protein_ | -0.00086567 | 0.0038713 | -0.0015832 |
| SCTX01 | -0.0011058 | 0.0031139 | -0.0023239 |
| SLPTx-SSD976 | 0.001644 | 0.0029309 | 0.0057874 |
| NOT_TOXIN_INSUFFICIENT_EVIDENCE_Avidin-related | 0.00042574 | 0.0028701 | 0.0057095 |
| M12A | 0.00010724 | 0.002735 | -0.0020747 |
| Purple acid phosphatase | -0.00019831 | 0.0025399 | -0.0023349 |
| CAA-like | 0.0017672 | 0.002425 | -0.013202 |
| SLPTx-SSD558 | 0.0020349 | 0.0023011 | 0.0064199 |
| NOT_TOXIN_Calcium-transporting_ATPase | -0.00042812 | 0.0013772 | -0.0037622 |
| NOT_TOXIN_structural_component | 6.76E-05 | 0.00026983 | 0.0010198 |
| SLPTX05 | -0.0057793 | 0.00020779 | 0.018965 |
| NOT_TOXIN_Myosin_tail_1_domain-containing_protein | 0.0039516 | -0.00012697 | 0.0078781 |
| Unchar06 | -0.0064509 | -0.0002644 | 0.012953 |
| Cathepsin | -0.013977 | -0.00065998 | 0.028952 |
| MCSACoa | -0.0062337 | -0.00073132 | 0.0048218 |
| Vigilin | 0.0007449 | -0.0010078 | -0.0074429 |
| NOT_TOXIN_NO_SIGNALP_Pyruvate_kinase | 0.0075617 | -0.0010665 | 0.01385 |
| Cyclase-like_protein | 0.0062243 | -0.0014395 | 0.013401 |
| Kazal_domain | 0.0037601 | -0.001482 | 0.0035459 |
| NOT_TOXIN_ATP_synthase | 0.0037601 | -0.001482 | 0.0035459 |
| NOT_TOXIN_tubulin_alpha_chain_ | 0.0094066 | -0.0025896 | 0.013191 |
| Unchar13 | -0.0031431 | -0.0026427 | 0.00028515 |
| Unchar10 | -0.054173 | -0.0028485 | 0.036889 |
| SLPTX02 | 0.011722 | -0.0029657 | 0.01301 |
| Lysosomal_lipase | -0.0023979 | -0.0032609 | 0.01077 |
| NOT_TOXIN_INSUFFICIENT_EVIDENCE_Alpha-L-Fucosidase | 0.036985 | -0.003603 | 0.059033 |
| NOT_TOXIN_Tropomyosin_ | -0.00066742 | -0.0036676 | 0.0017744 |
| Haem-binding_protein | 0.025239 | -0.0039412 | 0.0572 |
| NOT_TOXIN_Myosin_heavy_chain_protein | 0.0089048 | -0.003955 | 0.02201 |
| Apolipoprotein D | 0.0074011 | -0.0042081 | 0.03414 |
| SLPTX29 | -0.0059585 | -0.004393 | 0.0078953 |
| SLPTX47 | 0.0029427 | -0.0046071 | 0.0077459 |
| Carboxylic ester hydrolase | -0.009081 | -0.0046865 | 0.00098509 |
| Thioester-containing protein 4 | 0.0013622 | -0.0047429 | 0.014316 |
| NOT_TOXIN_NO_SIGNALP_Calponin | 0.0021271 | -0.0053793 | 0.0003836 |
| DUF1397 | -0.00085945 | -0.0056555 | 0.012438 |
| ARYLSULFATASE_ | 0.0087004 | -0.0058521 | 0.033878 |
| SLPTX17 | -0.091927 | -0.0065785 | 0.15052 |
| NOT_TOXIN_Arginine_kinase_ | 0.020283 | -0.0066442 | 0.03725 |
| SCTX02 | 4.97E-05 | -0.0085573 | 0.015013 |
| Fibronectin_or_immunoglobulin-like | 0.098714 | -0.0094905 | 0.2572 |
| NOT_TOXIN_Histone_H2A | 0.016223 | -0.0098212 | 0.013555 |
| Ige-ESP-like | -2.75E-05 | -0.010456 | -0.0039004 |
| NOT_TOXIN_tubulin_beta_chain_ | 0.0052198 | -0.010529 | 0.0057039 |
| Unchar03 | 0.0052198 | -0.010529 | 0.0057039 |
| Cysteine-rich venom protein | 0.010934 | -0.010699 | 0.011064 |
| NOT_TOXIN_Myosin_light_chain_protein | 0.013123 | -0.010783 | 0.02146 |
| SLPTX08 | -0.010395 | -0.011169 | -0.015639 |
| NOT_TOXIN_Histone_H2B | 0.0099278 | -0.012562 | 0.01704 |
| Transferrin | 0.17625 | -0.013104 | 0.22766 |
| GH18 | 0.023308 | -0.014681 | 0.050366 |
| Unchar33 | 0.018276 | -0.015116 | -0.0054203 |
| SLPTX16 | -0.058653 | -0.01562 | 0.0024365 |
| Neprilysin | 0.0161 | -0.018476 | 0.029657 |
| NOT_TOXIN_Vitellogenin-6 | 0.23079 | -0.021989 | 0.25465 |
| Carbon-nitrogen_hydrolase | 0.051072 | -0.024305 | 0.14137 |
| Unchar32 | 0.0060106 | -0.033646 | 0.048338 |
| SLPTX30 | 0.025719 | -0.035805 | 0.11299 |
| PLA2 | -0.10195 | -0.037595 | 0.10623 |
| Peptidase_M13 | 0.0097547 | -0.038093 | 0.1015 |
| Serpine | 0.053926 | -0.043391 | 0.13905 |
| NOT_TOXIN_Actin | 0.085649 | -0.044686 | 0.13815 |
| Calycin-Lipocalin | 0.040347 | -0.049691 | 0.083535 |
| Unchar11 | 0.00034163 | -0.05264 | 0.10433 |
| Lipid-transport_protein | 0.29752 | -0.11522 | 0.35995 |
| Alpha-2-macroglobulin | 0.24566 | -0.15007 | 0.44647 |
| GGT | -0.4013 | -0.36764 | 0.1583 |
